# Errors in peptide synthesis are a source of discrepancies in Aβ42 studies

**DOI:** 10.64898/2026.07.30.741826

**Authors:** Karolina Matulewska-Sobczuk, Katja Bernfur, Dev Thacker, Johan Wallerstein, Elin Stemme, Alexander Dear, Nils Lindblom, Ewa A. Andrzejewska, c Šneiderienė, Tuomas P.J. Knowles, Gunnar Gouras, Ulf Olsson, Sara Linse, Lei Ortigosa-Pascual

## Abstract

Amyloid-β42 (Aβ42) aggregation is highly sensitive to experimental conditions, making reproducibility a persistent challenge in Alzheimer’s disease research. Among the many variables that influence aggregation, the impact of peptide production remains poorly understood. Direct comparison of recombinant and chemically synthesised Aβ42 prepared under carefully controlled conditions reveals that, despite following similar aggregation mechanism and forming the same predominant fibril structures, synthetic Aβ42 aggregates more slowly and exhibits reduced seeding efficiency. Consequently, synthetic Aβ42 produces fewer oligomeric species and displays lower cellular toxicity. Mass spectrometric analyses identify low-abundance sequence imperfections introduced during peptide synthesis as the origin of these differences. By linking synthesis-derived imperfections to variations in Aβ42 behaviour, this work reveals a previously underappreciated source of discrepancies in amyloid studies. In addition, we provide a framework for evaluating the impact of sequence impurities on biophysical studies that are sensitive to peptide composition.

## INTRODUCTION

Alzheimer’s disease (AD) is the most common cause of dementia ^1,2^. The main molecular hallmark of AD is the aggregation of amyloid β peptide (Aβ), an intrinsically disordered protein that self-assembles into oligomeric and fibrillar species ^3^. Aβ is produced from the amyloid-β precursor protein (APP) by the action of β- and γ-secretases, with the former cutting mainly between M671 and D672 of APP and the latter around residues 708-714 in the membrane-embedded part of APP ^4^. This results in a mixture of Aβ peptides, most of which have between 37 and 43 residues. Decades of *in vitro* studies have shown that Aβ aggregation is highly dependent on sample preparation protocol and experimental conditions ^5,6^, but highly reproducible within defined samples having carefully controlled composition. Aggregation is highly sensitive to both intrinsic factors, such as peptide charge, hydrophobicity and length ^7–10^, and extrinsic factors ^11^ that influence aggregation kinetics, including additional peptides and proteins ^12–17^, small molecules ^18,19^, polymers ^12,20^, membranes ^21–23^, salt ^24–26^, pH ^27–29^, temperature ^30,31^, shear forces ^32^ and nanoparticles ^33,34^. Among all variables, peptide purity and sequence homogeneity prove to be crucial, as even small amounts of variant species can substantially alter aggregation kinetics, fibril morphology and oligomer formation ^6^. The level of sequence homogeneity may depend on how the peptide is produced and purified.

The *in vitro* Aβ literature can be divided between the use of recombinant expression or chemical synthesis to produce the Aβ peptide. Recombinant production, usually in bacterial expression systems, benefits from the high fidelity of the ribosome ^35,36^ and cellular quality control systems that minimise sequence errors ^37–40^. Therefore, the product of overexpression is highly sequence homogeneous, and its final purity depends on the purification method rather than on its synthesis route ^41^. Aβ42, the Aβ variant most studied in vitro due to its more pronounced amyloidogenic and toxic properties ^42^, is often produced recombinantly either with an additional N-terminal methionine residue ^11,17,25,43,44^, corresponding to residue 671 in APP, or through expression involving cleavable fusion tags that generate the native Aβ42 peptide starting at Asp672 ^44–46^. Alternatively, peptides can be chemically synthesised by various methods, including solid phase peptide synthesis (SPPS), liquid phase peptide synthesis, and native chemical ligation^47–49^. Among these, SPPS remains the dominant approach due to its flexibility and accessibility ^48,49^. Nevertheless, Aβ42 synthesis through SPPS is viewed as particularly challenging due to the peptide’s high hydrophobicity and aggregation-prone C-terminus ^50,51^. Therefore, the synthesis suffers from low yield ^52–55^, toxic by-products from the use of highly reactive chemicals ^48^, and by-products with closely similar properties that make them hard to remove to obtain a pure sample ^37,56–58^.

Several studies have reported that the source of Aβ42 has a significant impact on its behaviour ^31,59–62^. Aβ42 produced via chemical synthesis, here referred to as synthetic Aβ42 (sAβ42), has been shown to have a lower aggregation rate and neurotoxicity compared to recombinant peptides (rAβ42)^60^. Moreover, synthetic peptides from different manufacturers have been reported to form fibrils of different structures ^61^. In contrast, other studies found no differences in morphology, structure or toxicity between peptide sources ^63,64^. These conflicting observations reflect the sensitivity of Aβ to experimental conditions and sample preparation ^31,41^. The division of the literature into studies using sAβ42 and rAβ42 leads to discrepancies, risking unnecessary disagreement in key aspects of Aβ properties. Understanding the mechanistic basis of these differences may unify previous results and guide the choice of Aβ42 protein origin for future studies.

Here, we systematically compare recombinant and synthetic Aβ42 under carefully controlled monomer preparation and with high reproducibility. Using an array of biophysical methods, we show that while sAβ42 aggregates with a similar mechanism and generates fibrils of similar structure to those of rAβ42, as seen by cryo-EM, its aggregation is slower and its seeding efficiency is weaker. Strikingly, this leads to a lower oligomer production for the synthetic variant, translated into decreased cell toxicity. Detailed mass spectrometry analyses identify low-abundance sequence imperfections present in sAβ42 as a major factor in the described differences and highlight the importance of peptide source and preparation for reproducible amyloid studies.

## RESULTS

### Synthetic Aβ42 aggregates more slowly than recombinant Aβ42

Previous studies have shown that synthetic Aβ42 aggregates more slowly than rAβ42 ^59,60,62,65^. However, some monomer isolation techniques or the lack of their use could lead to the sample not being homogeneous enough to assess the origin of this difference. Here, using a well-established monomer preparation method^41^ that ensures the starting material does not contain any fibrillated species, we observed that rAβ42 aggregated faster than sAβ42 at all tested concentrations as monitored by ThT fluorescence, regardless of the source of the recombinant and synthetic proteins (Fig. 1a). Evaluating the aggregation of synthetic peptides from two different suppliers (here referred to as s1Aβ42 and s2Aβ42) showed that their half-time (t_1/2_) values were, respectively, 1.4 and 2.4 times higher relative to the recombinant peptide. s1Aβ42 was selected for subsequent experiments due to its higher reproducibility.

**Fig. 1.**
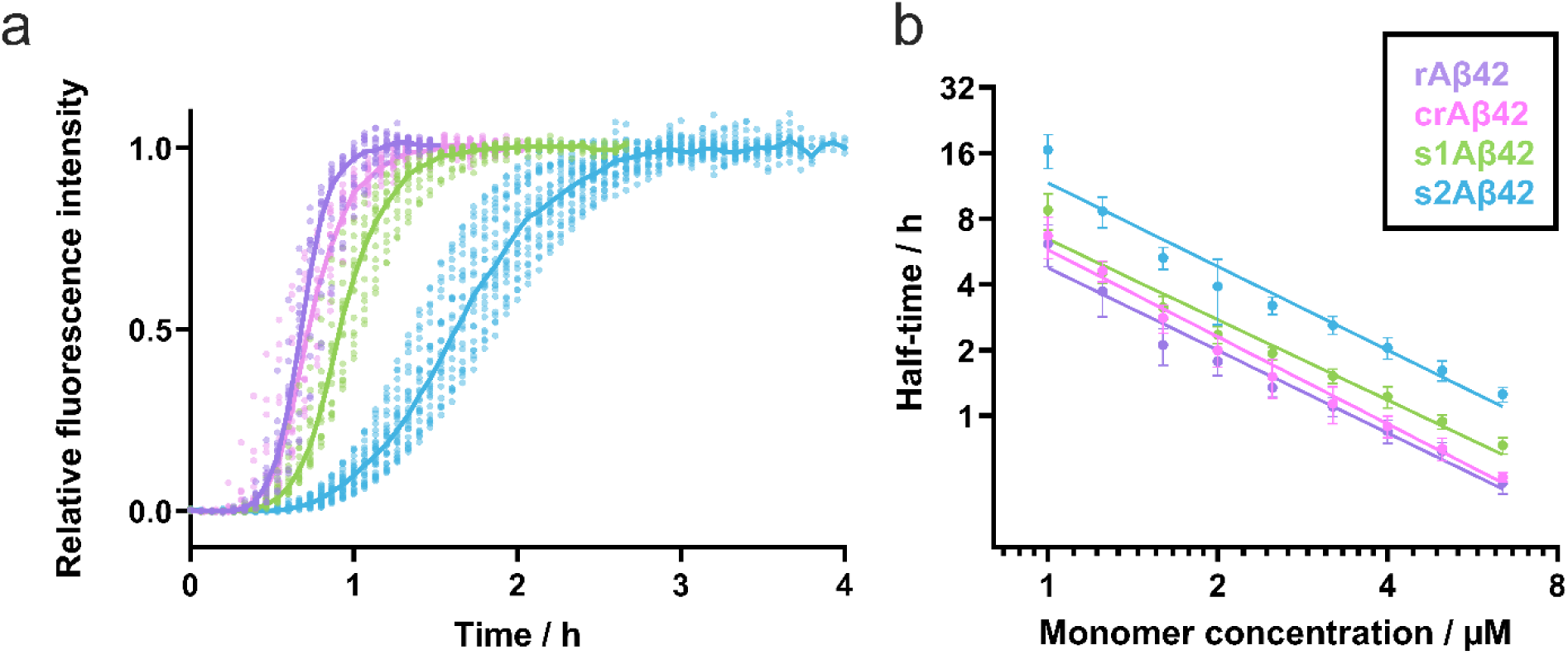
Synthetic Aβ42 aggregates more slowly than recombinant Aβ42 but via the same mechanism. Aggregation kinetics of in-house-produced rAβ42 (purple) versus the three commercial variants, crAβ42 (pink), s1Aβ42 (green), and s2Aβ42 (blue), were monitored by ThT fluorescence for starting monomer concentrations between 0.8 and 6.4 µM. Data were collected across 3 to 5 experiments, with 4-6 replicates per concentration per experiment. (a) Aggregation of Aβ42 at 5 µM initial monomer concentration. Data are plotted as dots and the median of the measurements for each protein is shown as a line. Half-times (t_1/2_) of aggregation were calculated as the time at which relative fluorescence intensity reached 0.5, with values of rAβ42(t_1/2_) = 0.69±0.06 h; crAβ42(t_1/2_) = 0.71±0.09 h; s1Aβ42(t_1/2_) = 0.94±0.08 h and s2Aβ4(t_1/2_) = 1.63±0.17 h for samples with a starting concentration of 5 µM. (b) The t_1/2_ of all replicates at each starting concentration was averaged and plotted as a function of initial monomer concentration. The similarity of the scaling exponent (γ) of the different proteins, obtained from the slope of each function, suggests a similar underlying mechanism for all proteins (rAβ42(γ) = -1.25±0.07; crAβ42(γ) = -1.32±0.08; s1Aβ42(γ) = -1.22±0.05; s2Aβ4(γ) = -1.27±0.05).

To assess whether residual impurities from SPPS ^48,66^ caused the slower aggregation of sAβ42, we subjected the peptides to additional lyophilisation and size exclusion chromatography (SEC) purification ^6,67^. Regardless of the number of SEC cycles applied, the slower aggregation of sAβ42 persisted (Supplementary Fig. 1). Notably, no significant differences were observed between the recombinant protein produced in-house and the commercially sourced recombinant protein (crAβ42) when treated with the same monomer isolation protocol, suggesting that the production method itself underlies the difference in Aβ42 aggregation. Interestingly, the introduction of an extra methionine at the N-terminus of Aβ42 has a lower impact on the aggregation than the production process ^44^ (Supplementary Fig. 2).

### Similar mechanism, different rate

A double logarithmic plot of t_1/2_ as a function of monomer concentration, m_0_, showed a linear (power law) dependence, *t*_1/2_∼*m*_0_*^γ^*, with scaling exponents that are nearly identical across peptide sources (rAβ42 γ = -1.25±0.07, crAβ42 γ = -1.32±0.08, s1Aβ42 γ = -1.22±0.05, s2Aβ42 γ = -1.27±0.05), suggesting that the underlying aggregation mechanisms are conserved despite the differences in aggregation rates (Fig. 1b). Furthermore, scaling exponents γ < -1 indicate that secondary nucleation is the dominating mechanism of aggregation of all systems ^15^. This conclusion was confirmed by detailed kinetic analysis using AmyloFit, where the secondary nucleation dominated model provided the best fit to the experimental data for all peptide variants (Supplementary Fig. 3, 4, 5).

We observed higher difference in the modelled k_+_k_2_ values than k_+_k_n_ (rAβ42 k_+_k_n_ = 1.7 ×10^2^ M^-2^ s^-2^ and k_+_k_2_ = 7.1×10^10^ M^-3^ s^-2^, for s1Aβ42 k_+_k_n_ = 1.5×10^2^ M^-2^ s^-2^ and k_+_k_2_ = 3.1×10^10^ M^-3^ s^-2^, for s2Aβ42 k_+_k_n_ = 0.6×10^2^ M^-2^ s^-2^ and k_+_k_2_ = 0.9×10^10^ M^-3^ s^-2^), while keeping n_c_ = n_2_ = 2 (Supplementary Fig. 6a, Supplementary Table 1). To evaluate the contribution of the kinetic parameter products, k_+_k_n_ and k_+_k_2_, to the model, we examined changes in the mean residual error (MRE) upon constraining individual products (Supplementary Fig. 6b). The broad distributions observed for k_+_k_n_ compared with the well-defined minima of k_+_k_2_ indicates that the model is considerably more sensitive to k_+_k_2_, whereas k_+_k_n_ is less constrained, making the comparison of k_+_k_n_ between the two systems less reliable. Subsequently, we estimated the differences in individual aggregation rate constants with an additional series of reactions (Supplementary Section S5: Supplementary Fig. 7). However, the differences among all the calculated rate constants were smaller than an order of magnitude, making it difficult to pinpoint one mechanism as the sole responsible.

### Peptide source impacts solubility and seeding efficiency with minor changes in polymorph distribution in fibril population

We examined the seeding capacity of the two types of peptides through self-seeding experiments (Fig. 2a). Adding as little as 0.3% (w/w) seeds shortened the aggregation t_1/2_ of rAβ42 approximately 3-fold (t_1/2, 0% seed_ = 0.97 h, t_1/2, 0.3% seed_ = 0.35 h) in line with earlier studies ^15,68^, whereas for s1Aβ42 the same seed concentration only sped up the process by a factor of ∼2 (t_1/2, 0% seed_ = 1.17 h, t_1/2, 0.3% seed_ = 0.67 h). Similarly, higher seed concentrations had a greater effect in decreasing the t_1/2_ of rAβ42 when compared to s1Aβ42. To identify the origin of the stronger seeding capacity of rAβ42, cross-seeding experiments were performed using the same protocol as for the self-seeding, but with the recombinant seed for the synthetic monomer, and *vice versa* (Fig. 2a). The difference in seeding efficiency persisted between the two systems when comparing the monomer source. In both self- and cross-seeding experiments, the reactions with monomeric rAβ42 showed a higher seeding efficiency than those with monomeric sAβ42, regardless of the seed type used (Fig. 2a). Thus, the seeding efficiency is clearly determined by the monomer rather than the fibril.

**Fig. 2.**
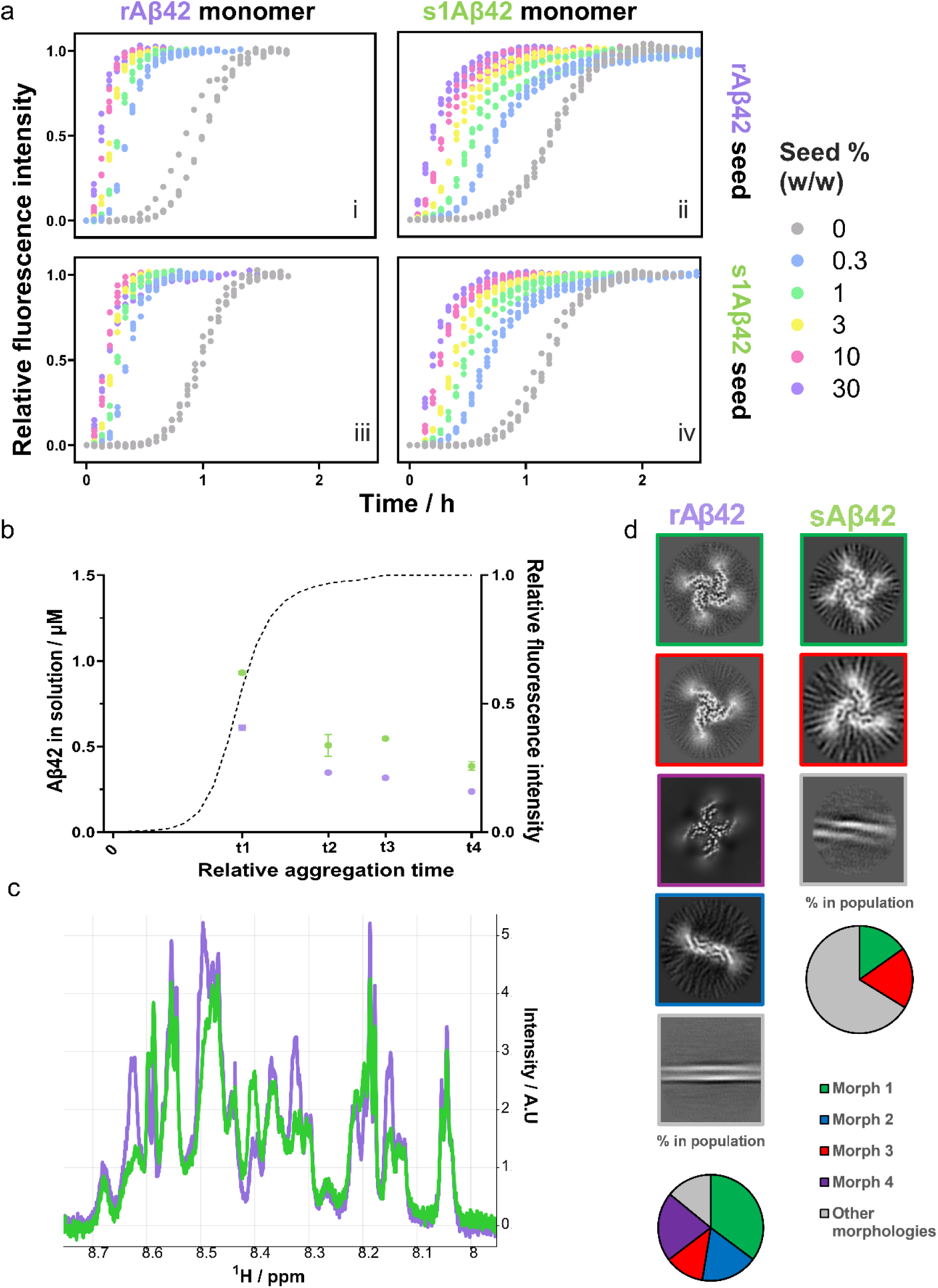
Synthetic Aβ42 has lower seeding efficiency but similar fibril morphologies compared with recombinant Aβ42. (a) Seeded aggregation monitored using ThT fluorescence. 6 µM monomeric rAβ42 (a & c) or s1Aβ42 (b & d) were aggregated in the presence of seeds made with either rAβ42 (a & b) or s1Aβ42 (c & d) at w/w concentrations of 0, 0.3, 1, 3, 10 and 30% (grey, blue, green, yellow, pink, and purple, respectively). (b) Concentration of Aβ42 remaining soluble during non-seeded aggregation of 2 µM rAβ42 (purple) and s1Aβ42 (green). Samples were collected at time points corresponding to different stages of the aggregation reaction, indicated relative to the normalised aggregation curve (black dashed line) and centrifuged (t1 ≈ 3 h at t_1/2_, t2 ≈ 7 h just before the plateau phase, t3 ≈ 20 h plateau, t4 ≈ 36 h later in plateau). The concentration of the supernatant, containing the soluble fraction, was measured with HPLC-MS. Data are presented as mean ± SD. (c) 1D ^1^H NMR spectrum of s1Aβ42 (green) and rAβ42 (purple), zoomed in on the H_N_ regions, recorded at 2 °C.(d) Representative cryo-EM 3D classes identified for rAβ42 (up) fibrils (left) and the relative abundance of each morphology within the classified fibril population (right). Representative cryo-EM 3D classes identified for s1Aβ42 (down) fibrils (left) and the relative abundance of each morphology within the classified fibril population (right).

To compare the apparent solubility of the two peptides, we measured the concentration of free monomer left in solution as a function of aggregation time (Fig. 2b). s1Aβ42 showed a higher monomer concentration in solution than rAβ42 at all the measured points and exhibited around 25% higher apparent solubility than the recombinant peptide after ca. 36 h equilibration (rAβ42 = 0.018 µM, s1Aβ42 = 0.025 µM).

The 1D ^1^H NMR spectra for rAβ42 and sAβ42 monomers showed differences between the species (Supplementary Fig. 8). These differences were present throughout the whole spectra, but particularly pronounced in the Cα, aromatic side chain and H_N_ regions. The 2D ^1^H-^1^H NOESY confirmed the differences and showed that the largest perturbations were related to different conformations at sites Gly9 and Tyr10 (Supplementary Fig. 9).

We performed cryo-EM data collection and processing to compare the fibril structures formed by the two peptides. From the selected particles, iterative 2D classification was performed to separate the polymorphs present in the fibril population, and subsequent 3D classification was performed for each polymorph. Four distinct polymorphs were resolved for rAβ42 (Fig. 2 c, left) and two for s1Aβ42 (Fig. 2d, down). Importantly, both morphologies observed in s1β42 were also present in the recombinant sample, showing that the synthetic peptide has the ability of forming at least some of the structures of rAβ42. The distribution over different structures is based on the dispersed fibrils on the grid only, and therefore may not reflect the true distribution in the sample as a large fraction of the fibrils end up in clumps, and a few on the carbon areas of the grid. Additionally, both datasets contained straight, untwisted fibrils forming a fraction of the population, but this was a much larger fraction of the fibrils in the s1Aβ42 dataset. The structures of these fibrils could not be resolved due to a lack of helical symmetry (Fig. 2d). Therefore, these results do not suggest that s1Aβ42 forms fibrils of fewer morphologies overall, but simply that it gives a larger population of non-helical, heterogeneous fibrils that cannot be classified into 3D classes.

Kinetic parameter evaluation, seeding efficiency and fibril morphology experiments were also performed for s2Aβ42, yielding similar differences relative to rAβ42 as observed for s1Aβ42, albeit more pronounced (Supplementary Fig. 5, 6, 10, Supplementary Table 1).

### Enhanced oligomer formation and LDH release of recombinant Aβ42

To evaluate whether the difference in kinetics led to a difference in oligomer production, we measured the oligomer population of the two peptides using µFFE ^32,69^. At their respective t_1/2_, where Aβ oligomer concentration is at its highest ^70^, rAβ42 showed a 15% higher oligomer count than s1Aβ42 (Fig. 3a, Supplementary Fig. 11). As oligomers are believed to mediate Aβ42’s toxicity ^71–75^, we evaluated whether the difference in oligomer production translated to a difference in cytotoxicity using N2a cells, a commonly used neuroblastoma cell line, and a lactate dehydrogenase (LDH) cytotoxicity assay (Fig. 3b). Treatment with s1Aβ42 did not show an increase in LDH release at any time point or concentration, in agreement with the literature data^59,60,62^. However, treatment with 8 µM rAβ42 for 48 h resulted in a 3.5-fold increase in LDH release after 48 h relative to the same concentration of s1Aβ42.

**Fig. 3.**
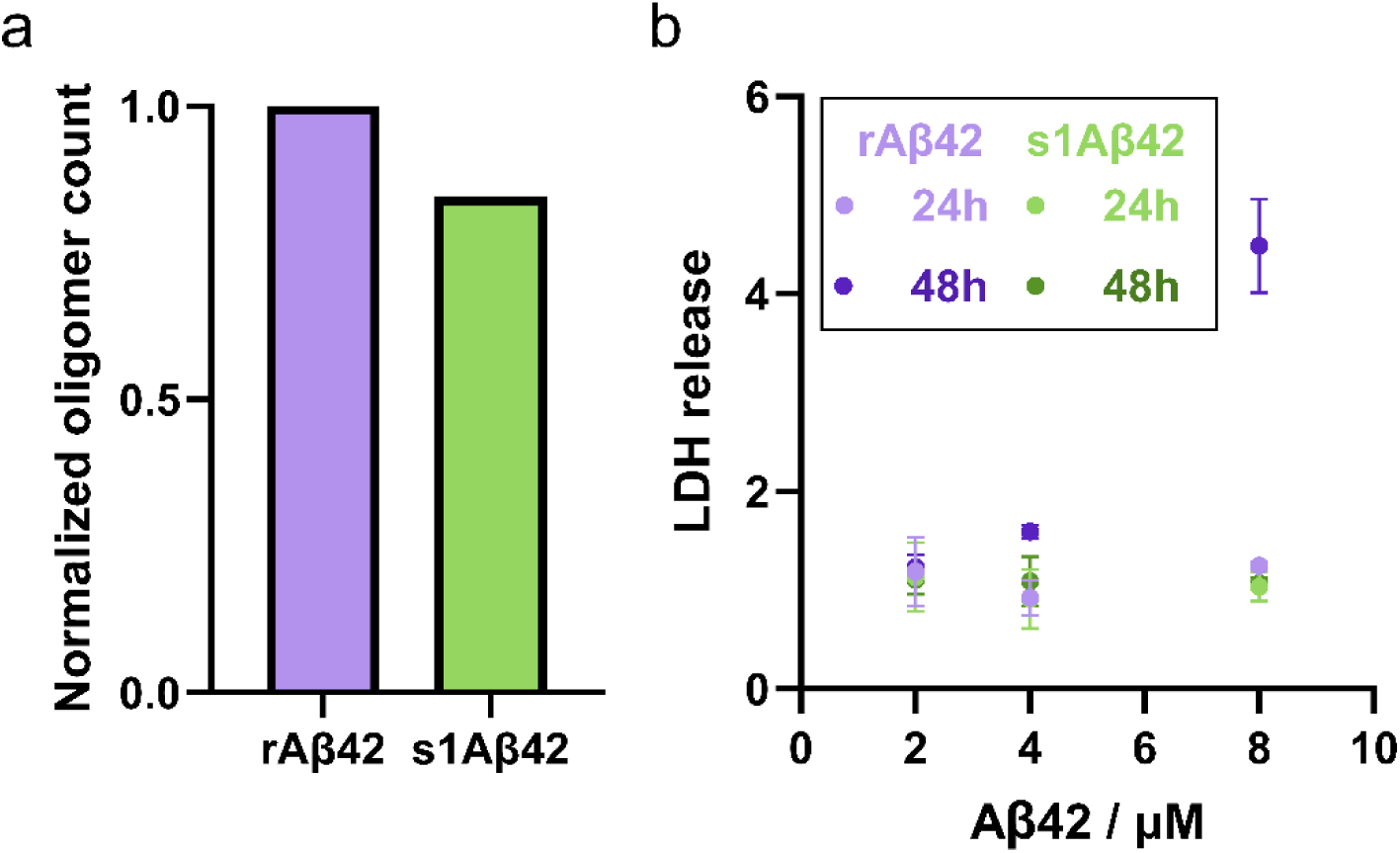
Recombinant Aβ42 forms more oligomers and is more toxic than synthetic Aβ42. (a) Relative oligomer count of rAβ42 (purple) and s1Aβ42 (green) at their respective t_1/2_ (Supplementary Fig. 11), as measured by µFFE ^69,104^. (b) Cytotoxicity of rAβ42 (purple) and s1Aβ42 (green) in N2a cells at peptide concentrations of 2, 4 and 8 µM following 24 (light purple/light green) and 48 h (dark purple/dark green) incubation. Cytotoxicity was assessed by measuring lactate dehydrogenase (LDH) release and normalised to the control (buffer) at each time point. Data are shown as the mean of 3 replicates from 3 experiments ± SD.

### Peptide synthesis produces a higher fraction of imperfect peptides

Contaminants in the sample can profoundly impact the behaviour of Aβ42 ^6,48,67^. However, the persistent differences between rAβ42 and s1Aβ42, even after extensive purification, suggest that the primary source of the discrepancy lies in the molecular origin of the peptides themselves rather than residual contaminants from the purification process. Aβ42 is a particularly difficult peptide to synthesise due to its highly hydrophobic and β-sheet-prone C-terminal region, which promotes aggregation both on-resin and in solution ^59,60,76^. That aggregation reduces reagent accessibility to the growing peptide chain, which can lead to inefficient amino acid coupling, incomplete Fmoc removal and formation of by-products with truncations or deletions. In addition, insufficient washing between couplings can lead to duplications in the sequence ^77,78^. Moreover, the potential presence of racemised amino acids could further influence aggregation kinetics and secondary structure of Aβ42 fibrils^56,58,79^. Finally, the purification of synthetic Aβ42 presents an additional challenge as the peptide can rapidly aggregate upon cleavage from the resin ^51^. Together, these factors make it hard to achieve high-yield synthetic Aβ42 production without compromising the peptide quality ^66^. As a result, Aβ42 produced via peptide synthesis is at increased risk of sequence errors, which could explain the differences reported in this manuscript.

To evaluate this, using mass spectrometry, we searched for deletion and duplication variants in both rAβ42 and s1Aβ42. Multiple peptides with deletions and duplications were detected for both systems with high coverage (Supplementary Table 2, 3; Supplementary Information 12). The relative abundance of each deletion and duplication variant was estimated from their UV absorbance intensity at 280 nm and normalised by the absorbance intensity of the corresponding wild-type variant containing the correct residue. Across all detected amino acid deletion and duplication variants, a higher proportion of peptides with an error was detected for s1Aβ42 than for rAβ42 (Fig. 4a). Considering all detected incorrect peptide variants, ∼8 times more deletions and ∼38 times more duplications were detected for s1Aβ42 relative to rAβ42. Thus, while deletion variants were more abundant than duplications for both peptides, duplications were detected more frequently in s1Aβ42 relative to rAβ42. Notably, all detected sequence alterations were localised within residues 10-27, the end of the N-terminal region of the peptide and a significant part of the fibril core encompassing the charged turn loop, potentially impacting both aggregation kinetics and fibril structure ^8,9,25,79–81^.

**Fig. 4.**
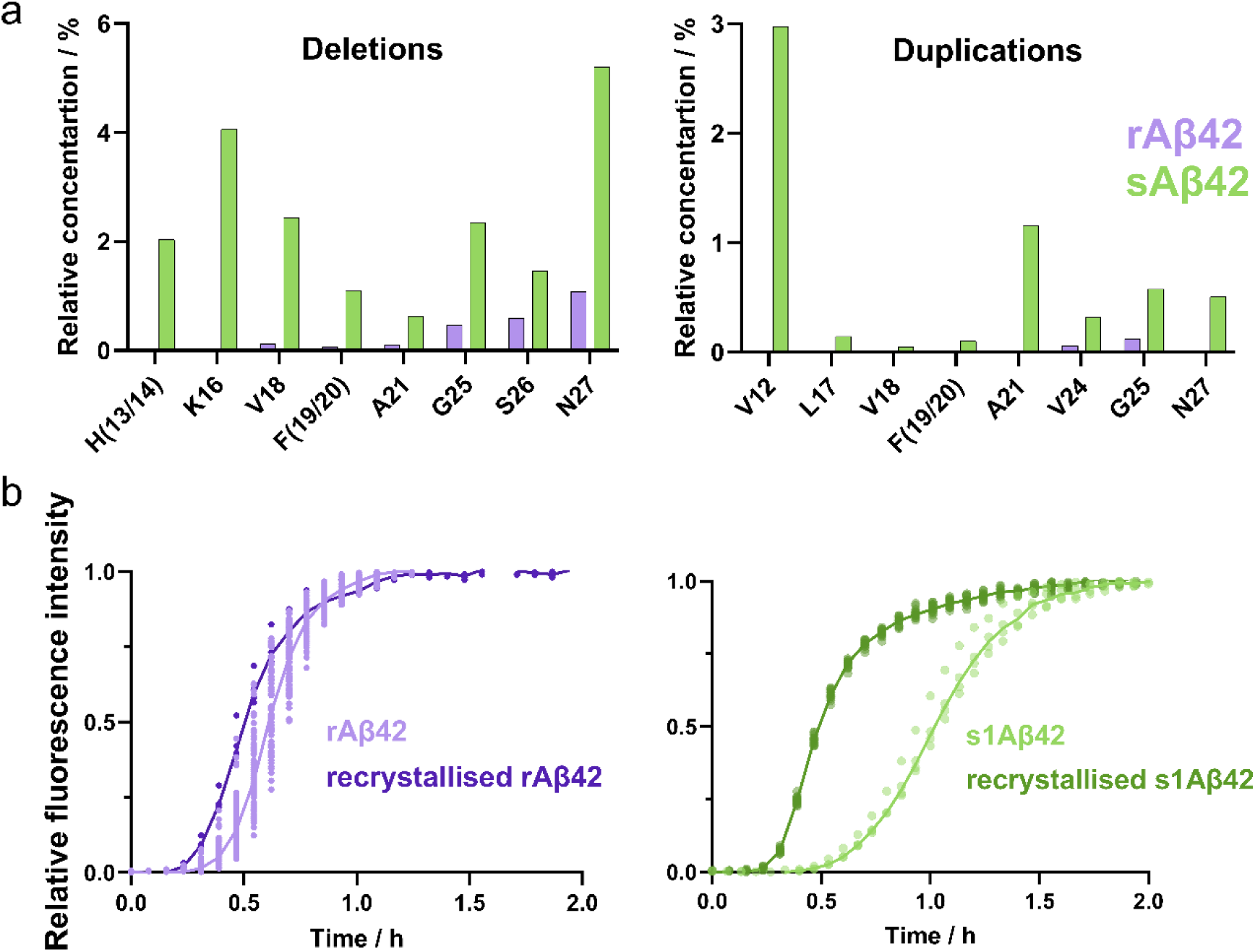
Synthetic Aβ42 contains more sequence variants than rAβ42, which can be removed by recrystallisation. (a) Relative concentration of Aβ42 peptides containing sequence deletions (left) and duplications (right) detected by MS/MS mass spectrometry in rAβ42 (purple) and s1Aβ42 (green). For each identified sequence variant with a single amino acid deletion or duplication, the concentration was measured from its 280 nm absorbance peak intensity and normalised to the concentration of the peptide containing that same amino acid in its wild-type form (Supplementary Table 2, 3). (b) Aggregation kinetics of the original (light purple/light green) and recrystallised (dark purple/dark green) peptide monitored by normalised ThT fluorescence intensity. Recrystallisation was performed by dissolving aggregated fibrils in 8 M GuHCl, subjecting them to SEC and aggregating again, enriching the aggregation-compatible peptide fraction. Lines represent the median of the data points.

Given that there is a small population of imperfect peptides in s1Aβ42 left in the bulk after aggregation, we applied an enrichment approach analogous to recrystallisation, a common purification strategy in organic synthesis ^82^. Assuming that the thermodynamically preferred species is incorporated into an ordered phase while some of the impurities remain in solution, fibrils at the end stage of the aggregation process are expected to consist mainly of wild-type monomers, and monomers of those peptides with sequences compatible with the wild-type fold. Peptides incompatible with this fold are expected to remain in the soluble phase. Following this reasoning, we collected fibrils at the end of an aggregated reaction by centrifugation to enrich aggregation-compatible sequences while excluding incompatible variants. The pelleted fibrils were dissolved in 8 M guanidine hydrochloride (GuHCl) and subjected to SEC and re-aggregated, monitored by ThT fluorescence. In case of rAβ42, the change in the t_1/2_ after the recrystallisation is negligible, whereas s1Aβ42 showed a twofold decrease in t_1/2_ (s1Aβ42 t_1/2_ = 1.06 h before and s1Aβ42 t_1/2_ = 0.5 h after recrystallisation) (Fig. 4b). These results show that the removal of the “incompatible” sequence variants restores faster aggregation kinetics of s1Aβ42, supporting the hypothesis that those variants are responsible for the different behaviour of the two species.

## DISCUSSION

In this study, we compared synthetic and recombinant Aβ42, showing that sAβ42 aggregates more slowly than its recombinant counterpart even after thorough purification and monomer isolation procedures. Moreover, we show that the synthetic peptide is less efficient at seeding, a behaviour governed primarily by the monomer rather than the fibril, and shows a higher apparent solubility than rAβ42. Additionally, sAβ42 produces fewer oligomers, which stands in agreement with its lower toxicity. Despite these differences, both peptides form fibrils of highly similar morphologies.

The observed differences in aggregation kinetics could plausibly arise from inaccuracies in the peptide concentration determination. However, based on the concentration dependence shown in Fig. 1b, we calculated that the concentration error for the synthetic peptides to overlap with rAβ42 would require the actual concentrations of s1Aβ42 and s2Aβ42 being only 77% and 50% of their absorbance-based concentrations, respectively. When estimating the concentration difference by overlapping 1D ^1^H NMR spectra and comparing their signal intensities, the peptide concentrations differed by no more than 15% (Supplementary Fig. 8). Moreover, if the slower aggregation reflected merely on a lower peptide concentration, differences in the final fibril amount should be apparent in the ThT plateau intensity, which were not observed (Supplementary Fig. 12). Together, these observations indicate that the kinetic differences arise from intrinsic properties of the peptides rather than inaccuracies in concentration determination.

To investigate the molecular origin of these differences, we searched for deletion and duplication variants in both peptides using mass spectrometry. Although such variants were detected in both samples, they were consistently more abundant in sAβ42, in line with the fundamentally different synthesis routes of the two peptides. Recombinant expression benefits from the high fidelity of ribosomal translation, whereas SPPS risks accumulating coupling errors during repeated synthesis cycles ^51^. Even assuming high coupling efficiency during SPPS, the cumulative yield of a perfectly synthesised 42-residue peptide is substantially reduced. Many of the imperfect peptide variants may still co-aggregate with wild-type (WT) Aβ42, especially those with variations within the flexible N-terminal region, as reported previously ^8–10^. However, a small fraction, around 1% (Fig. 2b), appears unable to aggregate on the experimental timescale. The acceleration of aggregation following fibril enrichment further suggests that the variants are less compatible with the ordered fibril structure (Fig. 4b). Consequently, fibrils formed from synthetic peptides may consist of both the WT and some imperfect peptides, being kinetic products of aggregation with lower stability than the system with fibrils formed only with the wild-type peptide.

A fraction of the sample containing imperfect sequences successfully explains the various differences reported in this and previous papers. Aside from a reduced concentration of aggregation-competent peptides for sAβ42 leading to slower aggregation kinetics, all stages of the aggregation pathway can be affected by a small population of imperfect peptides: During primary nucleation, imperfect peptides may bind to WT sequences but be unable to nucleate, kidnapping WT monomers and reducing the rate of nuclei formation. During elongation and secondary nucleation, non-WT variants may transiently occupy fibril ends and fibril defects, respectively, blocking the catalytic site for WT peptides while being unable to adopt the correct structure themselves, thereby slowing down both processes ^83^. This behaviour, akin to a competitive inhibitor, would explain the “inhibition-like” effect reported in previous studies when adding sAβ42 to rAβ42 ^59^. Reduced nucleation efficiency of s1Aβ42 would also be expected to produce fewer oligomers ^84^ and, since oligomers are believed to represent the major toxic species in Aβ aggregation ^3,59,60,71–73,85–88^, the higher rAβ42 cytotoxicity observed by us and in previous studies ^3,57,58,70–73,83–85^.

The differences in seeding efficiency between sAβ42 and rAβ42, being dominated by the monomer rather than the fibril identity, also align with imperfect sequences as an explanation. Firstly, while the half-time of a seeded aggregation is directly proportional to the monomer concentration, the seed concentration has a logarithmic effect on it. Thus, the same error in effective concentrations will always be more apparent for monomers. Secondly, if imperfect monomers inhibiting catalytic sites are the main mechanism of aggregation retardation for sAβ42, these species will be present in the reactions with monomeric sAβ42, while their concentration will be reduced in reactions that use sAβ42 fibrils, which would explain monomeric identity dominating the seeding behaviour. Finally, while NMR measurements hinted at differences between the monomeric populations, cryo-EM results indicate that fibril morphologies for both peptide sources are similar, and thus the difference between using either seed type is minimal. It should be noted that there was a larger proportion of imaged sAβ42 fibrils that could not be resolved in cryo-EM when compared to rAβ42, which may reflect increased morphological heterogeneity arising from sequence imperfections. The fact that s2Aβ42, the synthetic variant that aggregated slowest, also shows a higher fraction of non-resolvable fibril morphologies than both s1Aβ42 and rAβ42 supports this (Supplementary Fig. 10).

Amyloid formation is dramatically sensitive to subtle changes in its molecular environment ^5,6^. Therefore, identifying and understanding the variables that govern the aggregation is crucial for interpreting experimental data and producing reliable models. This study explains previous conflicting reports regarding Aβ42 aggregation kinetics and toxicity across different sources of Aβ42, and shows how these factors also affect its seeding behaviour, oligomer production, and fibril morphologies. Since we identify sequence imperfections originating from its synthesis process as the source of the observed differences, it is reasonable to evaluate whether these results apply to other difficult peptides ^89^. However, given Aβ42 is a particularly challenging sequence to synthesise with SPPS ^50,51,76,78^, these findings could potentially apply only to Aβ42. Nevertheless, the role the protein source plays in its *in vitro* behaviour should not be underestimated, especially with amyloid crosstalk studies on the rise ^90–92^, where combining amyloids affected differently by their production process could exponentially complicate the interpretation of their interactions.

The practical question arising from these findings is which Aβ42 source should one use. While synthetic peptides offer advantages in terms of the convenience of purchase, recombinant Aβ42 offers superior sequence homogeneity and reproducibility ^6^. Thus, for studies where precise control of aggregation kinetics is required, we recommend using a recombinant peptide. In that regard, we observe AβM1-42 to behave nearly identically to rAβ42 (Supplementary Fig. 2), being a convenient alternative to the wild-type peptide, and easier to produce and to purify. Furthermore, our results show that commercially available recombinant Aβ42 (crAβ42) can behave comparably to in-house purified peptide as long as it is thoroughly monomerised prior to use, i.e. dissolved in GuHCl and purified by SEC (Fig. 1). Reproducing key findings using properly purified recombinant peptide in an ensured monomeric state may help reduce sample heterogeneity and improve reproducibility across studies.

It should be noted that, while this study only focuses on the difference between sAβ42 and rAβ42, other protein sources present more complex differences. Aβ expressed in a mammalian cell system or directly extracted from the brain has been shown to aggregate faster and to be more toxic than either recombinant or synthetic protein^62,93^. Such differences may contribute to some of the variability and reproducibility issues that have long characterised the amyloid field ^41,94,95^. Thus, this study should not be taken as an indication of rAβ42 being the ideal model for *in vivo* Aβ42, but rather as an example of the potential bias that comes from the choice of system, and a stepping stone for the full characterisation of these.

Taken together, our study identifies sequence heterogeneity introduced during peptide synthesis as a major determinant of Aβ42 aggregation and toxicity, and it establishes peptide source as a critical experimental parameter in amyloid research. At the same time, we provide a template for evaluation of the role of imperfections generated during peptide synthesis, which can be applied to not only other amyloid systems, but to any study involving complex peptides produced via SPPS.

## METHODS

### Chemicals and Consumables

Synthetic Aβ42 peptides were purchased from AnaSpec (Fremont, CA, USA, 0.5 g AS – 24224, Lot # 2055180), here referred to as s1Aβ42 and from Bachem (California Peptide Research, Inc., Napa, CA, 5 mg 4014447.5000 Lot # 1071350), here referred to as s2Aβ42. Commercial recombinant Aβ42 was purchased from rPeptide (batch), here referred to as crAβ42.

### Expression and purification of recombinant proteins

Recombinant Aβ42 (rAβ42) was expressed using the EDDIE mutant of the self-cleavable nPro tag fused to Aβ42 ^41,46^. The fusion protein sequence was: MELNHFELLYKTSKQKPVGVEEPVYDTAGRPLFGNPSEVHPQSTLKLPHDRGEDDIETTLRDLPRKGDCRSGNHLGPVSGIYIKPGPVYYQDYTGPVYHRAPLEFFDETQFEETTKRIGRVTGSDGKLYHIYVEVDGEILLKQAKRGTPRTLKWTRNTTNCPLWVTSCDAEFRHDSGYEVHHQKLVFFAEDVGSNKGAIIGLMVGGVVIA. *E. coli* BL21 DE3 pLysS* cells were transformed with a pET3a plasmid (Genscript) containing the EDDIE-Aβ42 construct. Expression was carried out in overnight expression medium (2.5 mM Na_2_HPO_4_, 2.5 mM KH_2_PO_4_, 12 mM (NH_4_)_2_SO_4_, 1 mM MgSO_4_, 0.1 g/L glucose, 0.4 g/L lactose, 1 g/L glycerol, 10 g/L NaCl, 10 g/L tryptone, 5 g/L Bacto yeast extract, 50 mg/L ampicillin, and 30 mg/L chloramphenicol). Following overexpression, the cell pellet from 4 L culture was sonicated five times in 80 mL of buffer A (10 mM Tris/HCl, 1 mM EDTA, pH 8.5), with centrifugation at 18 000 g for 7 min after each cycle. The pellet was dissolved in 150 mL of 10 M urea with 10 mM dithiothreitol (DTT) in buffer A, using a combination of sonication, grinding, and stirring. The resulting solution was then diluted with 150 mL of buffer A and applied to tandem 20 mL DEAE-Sepharose FF columns (GE Healthcare). The columns were washed with 100 mL of buffer A containing 4 M urea and 1 mM DTT. Elution was achieved with a 0-0.4 M NaCl gradient in the same buffer. Fractions containing EDDIE-Aβ42 were diluted twofold with 1 M Tris, 1 mM EDTA, 5 mM DTT, pH 7.9, and dialysed against the same buffer in three shifts at 4 °C for 60 h to allow EDDIE to fold, resulting in its auto-cleavage and release of rAβ42. Dialysis bags (3.5 kDa MW cutoff) were boiled four times in Milli-Q water prior to use. The solution was then dialysed three times against 10 L of 5 mM Tris/HCl, 0.5 mM EDTA, pH 8.5. After the third dialysis, the solution was passed through 25 mL SP Sepharose to remove EDDIE and supplemented with 50 g Q-Sepharose Big Beads (GE Healthcare, equilibrated in buffer A), followed by incubation for 0.5 h on ice while stirring intermittently with a glass rod. The beads were collected using a Büchner funnel. Washing was performed with 200 mL buffer A, and elution was carried out in eight steps of 100 mL of buffer A containing 75 mM NaCl. The eluted fractions were analysed by SDS-PAGE. Fractions primarily containing rAβ42 were lyophilised and dissolved in 6 M guanidine hydrochloride (GuHCl), 20 mM sodium phosphate, pH 8.5, and subjected to size exclusion chromatography (SEC) in 20 mM sodium phosphate, 0.2 mM EDTA, pH 8.5 on a Superdex^TM^ 75 26/600 column at 4 °C. This step aimed to isolate the protein from remaining *E. coli* proteins, EDDIE, and aggregated species. The eluted fractions were examined using UV absorbance, agarose gel electrophoresis, and SDS-PAGE. Fractions corresponding to the centre of the rAβ42 monomer peak were pooled, lyophilised, and redissolved in 10 mL of 6 M GuHCl, 20 mM sodium phosphate, pH 8.5. Subsequently, a second round of SEC was then performed under identical conditions. The resulting monomer fractions were evaluated by mass spectrometry, pooled, aliquoted, lyophilised, and kept at -20 °C.

Previous publications have used a variant of Aβ42 containing the starting methionine, here referred to as Aβ(M1-42). This variant shows similar fibril formation kinetics as rAβ42, while being easier to produce ^44^. We expressed and purified Aβ(M1-42) as previously described^11,41^ and subjected it to the same kinetic analysis as performed for rAβ42 and sAβ42 (Fig. 1), confirming that the kinetic behaviour of Aβ(M1-42) and rAβ42 are indeed indistinguishable (Supplementary Fig. 2). Given the ease of production and handling of Aβ(M1-42), this variant was used for the recrystallisation experiment and NMR. A fluorescently labelled S8C mutant of Aβ(M1-42) was used for oligomer counting experiments. The Aβ(M1-42)-S8C mutant was expressed and purified as previously described^96^.

The Brichos domain of human pro-SPC (residues 88-197), here referred to as Brichos, was expressed and purified as previously described by Willander *et al.* ^17^.

### Monomeric Aβ42 preparation

To ensure a fully monomeric sample at the start of each experiment, all recombinant and synthetic Aβ42 variants were subjected to the protocol described by Cohen *et al* ^11^ and Silvers *et al.* ^44^. Samples were dissolved in 6 M GuHCl, pH 8.5, incubated for at least 20 min at room temperature, and purified by SEC on a GE Superdex^TM^ 10/300 75 Increase column in 20 mM sodium phosphate, 0.2 mM EDTA, pH 8.0 (reaction buffer). Elution was monitored by absorbance at 280 nm. The central part of the peak corresponding to monomeric Aβ42 was collected and maintained on ice until use. The concentration of Aβ42 was determined using the absorbance at 280 nm and a molar extinction coefficient of 1 400 M^-1^cm^-1^. For s2Aβ42, a cloudy precipitate formed upon dissolution in 6 M GuHCl. To remove the precipitate, the sample was centrifuged at 10 000 g for 10 min, and only the supernatant was used for one more round of SEC.

### Unseeded aggregation kinetics

Aggregation kinetics of Aβ42 were studied in the reaction buffer and monitored using 6 μM thioflavin T (ThT) fluorescence. Measurements were performed in a 96-well half-area PEGylated polystyrene plate (Corning 3881). The plate was carefully sealed to prevent evaporation or contamination and incubated in a plate reader at 37 °C (FLUOstar Omega, BMG Labtech, Offenburg, Germany). ThT fluorescence (excitation/emission: 448/480 nm) was recorded from the bottom of the plate continuously and without shaking. Cycle time was set to be the same as the minimum cycle time for all experiments to reduce idle time between cycles and increase reproducibility ^32^. Unseeded experiments were conducted with initial Aβ42 monomer concentrations ranging from 0.8 μM to 6.4 μM.

Aggregation of Aβ42 in the presence of Brichos was studied by following the aggregation of 5 µM monomeric Aβ42 in the absence or the presence of 7.5 µM Brichos, with the kinetics monitored by ThT fluorescence as described above.

### Seeded aggregation kinetics

Seeds were prepared under the same conditions as those used for aggregation kinetics, but with an initial monomer concentration of 10 µM. Control samples with ThT were used to monitor the aggregation. Once the control samples reached the plateau phase, seeds without ThT were collected and used immediately. Seeded experiments were run by mixing 6 µM monomeric Aβ42 with seeds at w/w concentrations of 0, 0.3, 1, 3, 10, and 30%.

Elongation-dominated kinetics experiments were performed using seeds sonicated prior to use to maximise the number of fibril ends and ensure a uniform size distribution. The sonication was performed using a tip sonicator at 4.3% amplitude for 90 s at 50% duty cycle, 1 s on, 1 s off. The aggregation of 3 µM monomeric Aβ42 in the presence of 8% sonicated seeds was monitored in a plate reader via ThT fluorescence as described above.

### Kinetic data analysis

ThT fluorescence intensity traces were normalised and aggregation curves across all concentrations (1 µM -6.4 µM) were globally fitted using AmyloFit ^68^. The model yielding the lowest mean residual error (MRE) was considered to best describe the data. The secondary nucleation-dominated unseeded aggregation model provided the best fit (Supplementary Fig. 3, 4, 5), in agreement with previous studies of Aβ42 aggregation ^9,11,23,97^. This was consistent across all experiments involving all Aβ42 species in the current study. The kinetic parameters derived from the fitting process with the lowest MRE were geometrically averaged to prevent amplification of variability in the resulting parameters. The MRE was evaluated as a function of a fixed kinetic parameter combination (k_+_k_n_ or k_+_k_2_) to generate a parabolic error landscape using AmyloFit (Supplementary Fig. 6).

Fibril elongation rates were assessed from the initial slopes of the fluorescence intensity versus time in seeded aggregation reactions (Supplementary Fig. 7b). For monomeric rAβ42, the linear region was identified between 0-0.13 h, whereas for monomeric s1Aβ42, it corresponded to 0-0.47 h. These experiments provided the ratio of the elongation rate constants (k_+_) of rAβ42 and sAβ42. Data from the experiment in the presence of Brichos were fitted to the elongation-*plus*-primary-nucleation model based on the assumption that Brichos completely blocks secondary nucleation ^32,98^, and allowing for a calculation of the difference in k_+_k_n_ between the two systems (Supplementary Fig. 7a). Using the k_+_ value determined from the elongation-dominated aggregation and the difference in k_+_k_n_ between rAβ42 and sAβ42, we calculated the difference in k_n_. Finally, the difference in k_2_ was estimated by fitting the experimental data in Amylofit while setting the proportions of k_n_ and k_+_ to those obtained above (full analysis in section S5 of the Supplementary Information).

### Solubility measurements

Monomeric rAβ42 and sAβ42 were aggregated at 2 µM and monitored via ThT fluorescence using a plate reader as described above. Samples were collected at time points corresponding to t_1/2_, the end of the exponential growth and the plateau of the aggregation reaction, as well as after 3 weeks of further incubation. The samples were centrifuged at 20 000 g for 25 min at room temperature, and the top of the supernatant was collected for analysis. The supernatant, containing the soluble fraction of Aβ42, was placed in a 96-well half-area PEGylated polystyrene surface plate (Corning 3686) and its concentration was determined using HPLC-MS (Shimadzu Nexera XR with LCMS-2020). Samples were injected with an autosampler onto a reverse-phase column (BIOshell A160 Peptide CN column 66966-U, Sigma-Aldrich). The elution had a constant flow rate of 0.5 mL/min for 10 min, with a linear gradient (5% to 95%) of acetonitrile in water, supplemented with 0.1% TFA. The absorbance was measured at 205 and 280 nm, see Supplementary Fig. 13 for examples of chromatograms. Aβ42 concentrations were determined based on the area of the peak identified as Aβ42 using a calibration curve (Supplementary Fig. 14). Note that the calibration curve was constructed from measurements on samples with known concentrations of recombinant Aβ(M1-42).

### CryoEM

Fibrils were prepared as described above, except without ThT. Grids were prepared for cryo-EM following standard vitrification procedures. Solutions containing fibrils were applied to glow-discharged 300 mesh lacey Formvar carbon-coated copper grids (Agar Scientific). Excess liquid was blotted away with a 2 s blot time and 0.5 s post-blot time before the grids were rapidly plunged into liquid ethane at approx. -184 °C using an automatic plunge freezer (Leica EM GP) under controlled temperature and humidity conditions. This vitrification process ensured the formation of thin liquid films in a glass-like state, preventing ice crystal formation and preserving the native microstructure of the samples.

Cryo-EM datasets for sAβ42 were collected at SciLife lab, Umeå, Sweden, using a Glacios electron microscope (Thermo Fisher) operated at 200 kV with a Falcon4 detector. Datasets were collected at a nominal magnification of 150 000x and set, yielding a pixel size of 0.95 Å. A defocus range of -1.4 to -2.6 µm and a total dose of 40 e-/Å^2^ was used over an exposure time of 3.67 s. The data were collected in TIFF format, and motion correction was performed in RELION-5 ^99^. Contrast Transfer Function (CTF) estimation was done for micrographs using CTFFIND4 ^100^. Manual picking of fibrils was done until roughly 50 000 particles of a 400-pixel box size were picked. This was used to train a Topaz dataset, and autopicking was performed using this trained Topaz dataset, which is incorporated in Relion-5. Different morphs were then separated out and processed with iterative rounds of 2D classification, and subsequent 2D classifications were performed separately for each set of morphs. The first round of 3D classification for each polymorph was done with initial models obtained from 2D class averages using the relion_helix_inimodel2d command ^101^. Multiple 3D classifications were performed to refine the helical rise and twist parameters for each polymorph until the peptide backbone was visible in the 3D classes.

Cryo-EM datasets for rAβ42 were collected at the University of Leeds Astbury Biostructure Laboratory. Data were collected using a Titan Krios electron microscope (Thermo Fisher) operated at 300 kV with a Falcon4 detector and Selectris energy filter (Thermo Fisher) operating with a 10 e-V slit. A nominal magnification of 130,000x was set yielding a pixel size of 0.95 Å. Movies were collected with a nominal defocus range of -1.4 to -2.6 µm and a total dose of 40 e-/Å^2^ was used over an exposure time of 4.48 s. The subsequent processing steps were performed as described above.

### Solution-state NMR spectroscopy

To compare rAβ42 and s1Aβ42 structural properties in the monomeric state, both peptides were purified by SEC as described before and supplemented with 3% (v/v) D_2_O to each sample. Concentrations of rAβ42 and s1Aβ42 were estimated by measuring absorbance at 280 nm to 20 and 40 µM, respectively. 1D ^1^H spectra were recorded on a Bruker Avance Neo 800 MHz 4-channel spectrometer with a TCI 3 mm CryoProbe (Bruker Biospin, Rheinstetten, Germany) using the standard water-suppression pulse program zgesgppe. Spectra were processed using TopSpin 4.3.0 (Bruker). Signal intensities were used to estimate differences in the peptides’ concentrations. Because of the limited overlap between 1D spectra, a precise quantitative comparison could not be performed. However, the concentration difference was estimated to be no greater than 15%. 2D NOESY and TOCSY spectra were also acquired to determine the differences between the peptides in more structural detail. The experiments were run at 2 °C for 18-20 h each.

### Oligomer measurements via µFFE

The reactions for liquid-electrode microchip free-flow electrophoresis (µFFE) were run at a labelled-to-unlabelled Aβ42 molar ratio of 1:3.5, based on previous reports showing that the aggregation and fibril morphology at this ratio are similar to those of fully unlabelled Aβ42 ^96^. The fluorescently labelled Aβ(M1-42)-S8C was mixed with either rAβ42 or s1Aβ42 at a 1:3.5 ratio to a final concentration of 2 µM. The aggregation was monitored following the quenching of the fluorescence of the Alexa Fluor 488 (Supplementary Fig. 11). Samples were collected at the respective half-times of their reactions and analysed by µFFE. Microfluidic devices were designed with AutoCAD software (Autodesk), and acetate transparencies (Micro Lithography Services) were used for photolithographic masks. SU-8 moulds were made via photolithographic processes previously described ^102^, and the features were evaluated by a profilometer (Dektak, Bruker). Polydimethylsiloxane (PDMS) (Dow Corning) was prepared by mixing a curing agent and base ratio of 1:10, poured on the mould, and degassed. After baking for 1.5 h at 65 °C, devices were cut and holed with biopsy punches for electrode insertion (1.5 mm) and tubing connection (0.75 mm). Devices were then cleaned in a sonication bath with isopropyl alcohol and bonded to glass slides with oxygen plasma. An additional prolonged exposure to oxygen plasma rendered the devices hydrophobic prior to their use ^103^. µFFE was run as previously reported ^104^. The liquid flow rate was regulated by syringe pumps (Cetoni neMESYS). Glass syringes (Gas Tight, Hamilton) were connected to the device with PTFE tubing, 0.012“ID x 0.030”OD (Cole-Parmer). Auxiliary buffer, electrolyte and sample were flown into the device at flow rates of 1000, 200 and 10 µL h^-1^, respectively. Bent syringe tips were inserted in the electrolyte outlets to allow application of electric potential via a programmable benchtop power supply (Elektro-Automatik EA-PS 9500-06). A custom-built single-molecule confocal fluorescence spectroscopy setup was used for all measurements. The setup was equipped with a 488 nm laser beam (Cobolt 06-MLD 488 nm 200 mW diode laser, Cobolt) and a time-correlated single photon counting module (TimeHarp 260 PICO, PicoQuant) with a 25 ps time resolution. To discern single-molecule bursts as discrete events from the acquired photon stream, a Lee filter of 3 was used, with fluorescence bursts characterised by a minimum of 0.001 µs inter-photon time and containing at least 22 photons. The single-molecule events were then recorded as a function of their position in the microfluidic device, with the degree of deflection corresponding to the charge-mobility relationship ^19^. Measurements of monomeric and fibrillar samples were done as a reference. Using their fluorescence as a lower and upper threshold, respectively, species between monomer and fibril size were counted as intermediate species, here referred to as oligomers.

### LDH cytotoxicity assay

10 000 N2a cells (neuroblastoma cell line) were seeded on a 96-well plate. 24 h later, the media were changed, and the cells were treated with 2, 4 or 8 μM of either rAβ42 or s1Aβ42. Media were collected after 24 and 48 h, and lactate dehydrogenase (LDH) activity was assessed by the CyQUANT™ LDH Cytotoxicity Assay Kit (Invitrogen, C20300). All experiments were done in triplicate, and each separate experiment was counted as one biological replicate (n). The assay was performed in accordance with the manufacturer’s instructions. In brief, 50 µL of media was added to a 96-well plate followed by 50 µL of reaction mixture, mixed well on a plate-shaker and then incubated at room temperature for 30 min, protected from light. The absorbance was then measured using the iMark™ Microplate Absorbance Reader (Bio-Rad 1681130XTU) at 680 nm and 490 nm. LDH activity was calculated as the 490-680 nm signal. LDH release of samples containing Aβ42 was normalised to a control containing only buffer for each of the recorded time points.

### Mass Spectrometry analysis

Prior to digestion of proteins, the pH of the samples was adjusted to 7.8 by adding ammonium bicarbonate to a final concentration of 100 mM. Digestion was performed by adding sequencing-grade modified trypsin (Promega, Madison, WI, USA) to the protein samples at a 1:50 (w/w) protease-to-protein ratio, followed by overnight incubation at 37 °C. The next day, formic acid was added to a final concentration of 0.5 %. Samples were then injected into an ultra-high-pressure nanoflow chromatography system (nanoElute, Bruker Daltonics). The peptides were loaded onto an Acclaim PepMap C18 (5 mm, 300 μm id, 5 μm particle diameter, 100 Å pore size) trap column (Thermo Fisher Scientific) and separated on a Bruker Pepsep Ten C18 (75 µm × 10 cm, 1.9 µm particle size) analytical column (Bruker Daltonics). Mobile phase A (2% acetonitrile, 0.1% formic acid) was used with the mobile phase B (0.1% formic acid in acetonitrile) for 45 min to create a gradient (from 2 to 17% B in 20 min, from 17 to 34% B in 10 min, from 34 to 95% B in 3 min, at 95%B for 12 min) at a flow rate of 300 nL/min and a column oven temperature of 50 °C. The peptides were analysed on a quadrupole time-of-flight mass spectrometer (timsTOF Pro, Bruker Daltonics) via a nano electrospray ion source (Captive Spray Source, Bruker Daltonics) in positive mode, controlled by the OtofControl 5.1 software (Bruker Daltonics). The temperature of the ion transfer capillary was 180 °C. A data-dependent acquisition method was used to select precursor ions for fragmentation with one TIMS-MS scan and 10 PASEF MS/MS scans. The TIMS-MS scan was acquired between 0.60–1.6 V s/cm^2^ and 100–1 700 m/z with a ramp time of 100 ms. The 10 PASEF scans contained a maximum of 10 MS/MS scans per PASEF scan with a collision energy of 10 eV. Precursors with a maximum of 5 charges with an intensity threshold of 5 000 a.u. and a dynamic exclusion of 0.4 s were used.

Raw data files were processed using Mascot Distiller (version 2.8, Matrix Science) to perform peak picking and generate Mascot generic files (MGF). Database searching was performed by submitting the processed data to a local Mascot Server (version 2.8.0). Custom in-house FASTA databases were constructed to identify specific Aβ42 variants. For the deletion library, 42 unique sequences were generated by systematically removing one amino acid at a time from each position of the wild-type sequence. Similarly, a duplication library was created consisting of 42 sequences, where each amino acid was systematically duplicated at its respective position. The following settings were used: precursor ion tolerance 10 ppm, MS/MS fragment mass tolerance 0.015 Da, trypsin as protease, 0 missed cleavage sites, and Methionine oxidation as variable modification. Label-free quantification (LFQ) of identified tryptic peptides containing either amino acid deletions or duplications was performed using the Average LFQ protocol within the Mascot Distiller Quantitation Toolbox. Peptide abundances were determined by the integration of precursor MS1 Extracted Ion Chromatograms (XICs). To derive relative protein abundance ratios, the total intensities of each variant-bearing peptide were normalised against their respective WT counterparts. To ensure high-confidence quantification, only peptides with a Mascot individual ion score > 33 and a minimum of two independent identifications per peptide were considered. Furthermore, XIC peak integrity was validated using a correlation threshold at 0.9 and a fraction threshold at 0.8. These thresholds were established through empirical optimisation, followed by manual inspection of the final spectra and data to ensure rigorous reproducibility (Supplementary Table 2, 3; Supplementary Section S12).

### Recrystallisation experiment

Monomeric Aβ42 was aggregated at 6.4 µM under the conditions described above for aggregation kinetics. After reaching the plateau phase, fibrils were pelleted by centrifugation at 20 000 g for 20 min, and the supernatant was discarded. The pellet was resuspended in 8 M GuHCl, pH 8.5, to dissociate the fibrils back to monomers. The sample was subjected to SEC to re-isolate monomeric Aβ42 as described above and subsequently aggregated at 6.4 µM, monitored by ThT fluorescence.

## ACKNOWLEDGEMENTS

We thank Max Lindberg for stimulating discussions and support with HPLC measurements, Karin Åkerfeldt for helpful discussions and Erik Golovtchenko for purifying AβM1-42 and for support in the lab.

The authors would like to acknowledge Umeå Centre for Electron Microscopy (UCEM) for technical assistance and access to electron microscopy. Support was provided by SciLife Lab national Cryo-EM Unit at Umeå University. The authors would like to acknowledge the University of Leeds Astbury Biostructure Laboratory for technical assistance and access to electron microscopy.

## AUTHOR CONTRIBUTIONS

Conceptualisation: S.L., L.O.P.

Methodology: K.M.S., K.B., D.T., E.S., J.W., N.L., E.A.A., G.S., L.O.P.

Data curation: K.M.S., K.B., D.T., E.S., J.W., N.L., E.A.A., L.O.P.

Formal analysis: K.M.S., U.O., S.L., L.O.P.

Resources: T.P.J.K., G.G., U.O., S.L.

Project Administration: U.O., S.L., L.O.P Supervision: T.P.J.K., G.G., U.O., S.L., L.O.P.

Writing - original draft: K.M.S., U.O., S.L., L.O.P.

Writing - review & editing: K.M.S., K.B., D.T., E.S., J.W., A.D., N.L., E.A.A., G.S., T.P.J.K., G.G., U.O., S.L. and L.O.P.

## REFERENCES

1. Silva, M. V. F. et al. Alzheimer’s disease: Risk factors and potentially protective measures. J. Biomed. Sci. 26, 1–11 (2019).

2. World Health Organization. Global action plan on the public health response to dementia 2017 - 2025. Geneva: World Health Organization 52 (2017).

3. Hampel, H. et al. The Amyloid-β Pathway in Alzheimer’s Disease. Mol. Psychiatry 26, 5481–5503 (2021).

4. Vassar, R. et al. β-Secretase cleavage of Alzheimer’s amyloid precursor protein by the transmembrane aspartic protease BACE. Science 286, 735–741 (1999).

5. Hortschansky, P., Schroeckh, V., Christopeit, T., Zandomeneghi, G. & Fändrich, M. The aggregation kinetics of Alzheimer’s β-amyloid peptide is controlled by stochastic nucleation. Protein Science 14, 1753–1759 (2005).

6. Hellstrand, E., Boland, B., Walsh, D. M. & Linse, S. Amyloid β-protein aggregation produces highly reproducible kinetic data and occurs by a two-phase process. ACS Chem. Neurosci. 1, 13–18 (2010).

7. Meisl, G. et al. Differences in nucleation behavior underlie the contrasting aggregation kinetics of the Aβ40 and Aβ42 peptides. Proc. Natl. Acad. Sci. U. S. A. 111, 9384–9389 (2014).

8. Szczepankiewicz, O. et al. N-Terminal Extensions Retard Aβ42 Fibril Formation but Allow Cross-Seeding and Coaggregation with Aβ42. J. Am. Chem. Soc. 137, 14673– 14685 (2015).

9. Thacker, D., Willas, A., Dear, A. J. & Linse, S. Role of Hydrophobicity at the N-Terminal Region of Aβ42 in Secondary Nucleation. ACS Chem. Neurosci. 13, 3477– 3487 (2022).

10. Weiffert, T. et al. Increased Secondary Nucleation Underlies Accelerated Aggregation of the Four-Residue N-Terminally Truncated Aβ42 Species Aβ5-42. ACS Chem. Neurosci. 10, 2374–2384 (2019).

11. Cohen, S. I. A. et al. Proliferation of amyloid-β42 aggregates occurs through a secondary nucleation mechanism. Proc. Natl. Acad. Sci. U. S. A. 110, 9758–9763 (2013).

12. Assarsson, A., Linse, S. & Cabaleiro-Lago, C. Effects of polyamino acids and polyelectrolytes on amyloid β fibril formation. Langmuir 30, 8812–8818 (2014).

13. Baine, M. et al. Inhibition of Aβ42 aggregation using peptides selected from combinatorial libraries. Journal of Peptide Science 15, 499–503 (2009).

14. Carrotta, R. et al. Inhibiting effect of α s1 -casein on Aβ 1-40 fibrillogenesis. Biochim. Biophys. Acta Gen. Subj. 1820, 124–132 (2012).

15. Cohen, S. I. A., Vendruscolo, M., Dobson, C. M. & Knowles, T. P. J. From macroscopic measurements to microscopic mechanisms of protein aggregation. Journal of Molecular Biology 421 160–171 (2012).

16. Månsson, C. et al. Conserved S/T Residues of the Human Chaperone DNAJB6 Are Required for Effective Inhibition of Aβ42 Amyloid Fibril Formation. Biochemistry 57, 4891–4902 (2018).

17. Willander, H. et al. BRICHOS domains efficiently delay fibrillation of amyloid β-peptide. Journal of Biological Chemistry 287, 31608–31617 (2012).

18. Habchi, J. et al. Systematic development of small molecules to inhibit specific microscopic steps of Aβ42 aggregation in Alzheimer’s disease. Proc. Natl. Acad. Sci. U. S. A. 114, E200–E208 (2017).

19. Härd, T. & Lendel, C. Inhibition of amyloid formation. Journal of Molecular Biology 421 441–465 (2012).

20. Heegaard, P. M. H., Boas, U. & Otzen, D. E. Dendrimer Effects on Peptide and Protein Fibrillation. Macromol. Biosci. 7, 1047–1059 (2007).

21. Baumann, K. N. et al. A Kinetic Map of the Influence of Biomimetic Lipid Model Membranes on Aβ42 Aggregation. ACS Chem. Neurosci. 14, 323–329 (2023).

22. Grey, M., Linse, S., Nilsson, H., Brundin, P. & Sparr, E. Membrane interaction of α-synuclein in different aggregation states. J. Parkinsons Dis. 1, 359–371 (2011).

23. Lindberg, D. J., Wesén, E., Björkeroth, J., Rocha, S. & Esbjörner, E. K. Lipid membranes catalyse the fibril formation of the amyloid-β (1–42) peptide through lipid-fibril interactions that reinforce secondary pathways. Biochim. Biophys. Acta Biomembr. 1859, 1921–1929 (2017).

24. Abelein, A., Jarvet, J., Barth, A., Gräslund, A. & Danielsson, J. Ionic Strength Modulation of the Free Energy Landscape of Aβ40 Peptide Fibril Formation. J. Am. Chem. Soc. 138, 6893–6902 (2016).

25. Meisl, G., Yang, X., Dobson, C. M., Linse, S. & Knowles, T. P. J. Modulation of electrostatic interactions to reveal a reaction network unifying the aggregation behaviour of the Aβ42 peptide and its variants. Chem. Sci. 8, 4352–4362 (2017).

26. Owczarz, M. & Arosio, P. Sulfate anion delays the self-assembly of human insulin by modifying the aggregation pathway. Biophys. J. 107, 197–207 (2014).

27. Buell, A. K. et al. Solution conditions determine the relative importance of nucleation and growth processes in α-synuclein aggregation. Proc. Natl. Acad. Sci. U. S. A. 111, 7671–7676 (2014).

28. Meisl, G., Yang, X., Frohm, B., Knowles, T. P. J. & Linse, S. Quantitative analysis of intrinsic and extrinsic factors in the aggregation mechanism of Alzheimer-associated Aβ-peptide. Sci. Rep. 6, 1–12 (2016).

29. Schützmann, M. P. et al. Endo-lysosomal Aβ concentration and pH trigger formation of Aβ oligomers that potently induce Tau missorting. Nat. Commun. 12, (2021).

30. Cohen, S. I. A. et al. Distinct thermodynamic signatures of oligomer generation in the aggregation of the amyloid-β peptide. Nat. Chem. 10, 523–531 (2018).

31. Foley, A. R. & Raskatov, J. A. Assessing Reproducibility in Amyloid β Research: Impact of Aβ Sources on Experimental Outcomes. ChemBioChem 21 2425–2430 (2020).

32. Axell, E. et al. The role of shear forces in primary and secondary nucleation of amyloid fibrils. Proc. Natl. Acad. Sci. U. S. A. 121 e2322572121 (2024).

33. Cabaleiro-Lago, C., Lynch, I., Dawson, K. A. & Linse, S. Inhibition of IAPP and IAPP(20−29) Fibrillation by Polymeric Nanoparticles. Langmuir 26, 3453–3461 (2010).

34. Vácha, R., Linse, S. & Lund, M. Surface effects on aggregation kinetics of amyloidogenic peptides. J. Am. Chem. Soc. 136, 11776–11782 (2014).

35. Maniatis, T., Fritsch, E. F. & Sambrook, J. Molecular Cloning: A Laboratory Manual.*No Title*. (Cold Spring Harbor Laboratory, Cold Spring Harbor, NY, 1982).

36. Yarus, M. The Accuracy of Translation. Prog. Nucleic Acid Res. Mol. Biol. 23, 195– 225 (1980).

37. Dyakin, V. V., Wisniewski, T. M. & Lajtha, A. Racemization in post-translational modifications relevance to protein aging, aggregation and neurodegeneration: Tip of the iceberg. Symmetry 13 455 (2021).

38. Ceccaldi, R., Rondinelli, B. & D’Andrea, A. D. Repair Pathway Choices and Consequences at the Double-Strand Break. Trends in Cell Biology 26 52–64 (2016).

39. Krokan, H. E. & Bjørås, M. Base excision repair. Cold Spring Harb. Perspect. Biol. 5, 1–22 (2013).

40. Myung, J., Kim, K. B. & Crews, C. M. The ubiquitin-proteasome pathway and proteasome inhibitors. Med. Res. Rev. 21, 245–273 (2001).

41. Linse, S. Expression and Purification of Intrinsically Disordered Aβ Peptide and Setup of Reproducible Aggregation Kinetics Experiment. in Intrinsically Disordered Proteins: Methods and Protocols (eds. Kragelund, B. B. & Skriver, K.) 731–754 (Springer US, New York, NY, 2020).

42. Jarrett, J. T., Berger, E. P. & Lansbury, P. T. Jr. The carboxy terminus of the β-amyloid protein is critical for the seeding of amyloid formation: Implications for the pathogenesis of Alzheimer’s disease. Biochemistry 32, 4693–4697 (1993).

43. Linse, S. et al. Nucleation of protein fibrillation by nanoparticles. Proc. Natl. Acad. Sci. U. S. A. 104, 8691–8696 (2007).

44. Silvers, R. et al. Aggregation and Fibril Structure of AβM01–42 and Aβ1–42. Biochemistry 56, 4850–4859 (2017).

45. Abelein, A. et al. High-yield Production of Amyloid-β Peptide Enabled by a Customized Spider Silk Domain. Sci. Rep. 10, 1–10 (2020).

46. Achmüller, C. et al. Npro fusion technology to produce proteins with authentic N termini in E. coli. Nat. Methods 4, 1037–1043 (2007).

47. Dawson, P. E., Muir, T. W., Clark-Lewis, I. & Kent, S. B. H. Synthesis of Proteins by Native Chemical Ligation. Science (1979). 266, 776–779 (1994).

48. Jaradat, D. M. M. Thirteen decades of peptide synthesis: key developments in solid phase peptide synthesis and amide bond formation utilized in peptide ligation. Amino Acids 50 39–68 (2018).

49. Merrifield, R. B. Solid Phase Peptide Synthesis. I. The Synthesis of a Tetrapeptide. J. Am. Chem. Soc. 85, 2149–2154 (1963).

50. Condron, M. M., Monien, B. H. & Bitan, G. Synthesis and purification of highly hydrophobic peptides derived from the C-terminus of amyloid β-protein. The Open Biotechnology Journal 2 87 (2008).

51. Paradís-Bas, M., Tulla-Puche, J. & Albericio, F. The road to the synthesis of ‘difficult peptides’. Chemical Society Reviews 45 631–654 (2016).

52. El-Faham, A. & Albericio, F. Peptide Coupling Reagents, More than a Letter Soup. Chem. Rev. 111, 6557–6602 (2011).

53. Lam, P. L., Wu, Y. & Wong, K. L. Incorporation of Fmoc-Dab(Mtt)-OH during solid-phase peptide synthesis: a word of caution. Org. Biomol. Chem. 20, 2601–2604 (2022).

54. Larionov, V. A., Stoletova, N. V & Maleev, V. I. Advances in Asymmetric Amino Acid Synthesis Enabled by Radical Chemistry. Adv. Synth. Catal. 362, 4325–4367 (2020).

55. Stráner, P., Taricska, N., Szabó, M., Tóth, G. K. & Perczel, A. Bacterial expression and/or solid phase peptide synthesis of 20-40 amino acid long polypeptides and miniproteins, the case study of Class B GPCR ligands. Curr. Protein Pept. Sci. 17, 147–155 (2016).

56. Angeletti, R. H. et al. Analysis of racemization during ‘Standard’ solid phase peptide synthesis: a multicenter study. Techniques in Protein Chemistry 8, 875–890 (1997).

57. Riester, D., Wiesmüller, K.-H., Stoll, D. & Kuhn, R. Racemization of Amino Acids in Solid-Phase Peptide Synthesis Investigated by Capillary Electrophoresis. Anal. Chem. 68, 2361–2365 (1996).

58. Zhou, Y. et al. Suppression of alpha-carbon racemization in peptide synthesis based on a thiol-labile amino protecting group. Nat. Commun. 14, 5324 (2023).

59. Adams, D. J., Nemkov, T. G., Mayer, J. P., Old, W. M. & Stowell, M. H. B. Identification of the primary peptide contaminant that inhibits fibrillation and toxicity in synthetic amyloid-β42. PLoS One 12, e0182804 (2017).

60. Finder, V. H., Vodopivec, I., Nitsch, R. M. & Glockshuber, R. The Recombinant Amyloid-β Peptide Aβ1-42 Aggregates Faster and Is More Neurotoxic than Synthetic Aβ1-42. J. Mol. Biol. 396, 9–18 (2010).

61. Suvorina, M. Y. et al. Studies of Polymorphism of Amyloid-β42 Peptide from Different Suppliers. Journal of Alzheimer’s Disease 47, 583–593 (2015).

62. Varshavskaya, K. B., Mitkevich, V. A., Makarov, A. A. & Barykin, E. P. Synthetic, Cell-Derived, Brain-Derived, and Recombinant β-Amyloid: Modelling Alzheimer’s Disease for Research and Drug Development. International Journal of Molecular Sciences 23 15036 (2022).

63. Walsh, D. M. et al. A facile method for expression and purification of the Alzheimer’s disease-associated amyloid β-peptide. FEBS Journal 276, 1266–1281 (2009).

64. Lieblein, T. et al. Structural rearrangement of amyloid-β upon inhibitor binding suppresses formation of Alzheimer’s disease related oligomers. Elife 9, 1–30 (2020).

65. Moore, B. D., Rangachari, V., Tay, W. M., Milkovic, N. M. & Rosenberry, T. L. Biophysical Analyses of Synthetic Amyloid-β(1−42) Aggregates before and after Covalent Cross-Linking. Implications for Deducing the Structure of Endogenous Amyloid-β Oligomers. Biochemistry 48, 11796–11806 (2009).

66. Teplow, D. B. Preparation of Amyloid β-Protein for Structural and Functional Studies. Methods Enzymol. 413, 20–33 (2006).

67. Martins, P. M. et al. MIRRAGGE – Minimum Information Required for Reproducible AGGregation Experiments. Front. Mol. Neurosci. 13, 1–18 (2020).

68. Meisl, G. et al. Molecular mechanisms of protein aggregation from global fitting of kinetic models. Nat. Protoc. 11, 252–272 (2016).

69. Dear, A. J. et al. Aβ Oligomer Dissociation Is Catalyzed by Fibril Surfaces. ACS Chem. Neurosci. 15 2296-2307 (2024)

70. Michaels, T. C. T. et al. Dynamics of oligomer populations formed during the aggregation of Alzheimer’s Aβ42 peptide. Nat. Chem. 12, 445–451 (2020).

71. Cleary, J. P. et al. Natural oligomers of the amyloid-β protein specifically disrupt cognitive function. Nat. Neurosci. 8, 79–84 (2005).

72. Kayed, R. et al. Common structure of soluble amyloid oligomers implies common mechanism of pathogenesis. Science 300, 486–489 (2003).

73. Sian, A. K. et al. Oligomerization of β-amyloid of the Alzheimer’s and the Dutch-cerebral-haemorrhage types. Biochemical Journal 349, 299–308 (2000).

74. van Dyck, C. H. et al. Lecanemab in Early Alzheimer’s Disease. New England Journal of Medicine 388, 9–21 (2023).

75. Nilsberth, C. et al. The ‘Arctic’ APP mutation (E693G) causes Alzheimer’s disease by enhanced Aβ protofibril formation. Nat Neurosci 4 887–893 (2001).

76. Tickler, A., Clippingdale, A. & Wade, J. Amyloid-B as a ‘difficult sequence’ in Solid Phase Peptide Synthesis. Protein Pept. Lett. 11, 377–384 (2005).

77. Behrendt, R., White, P. & Offer, J. Advances in Fmoc solid-phase peptide synthesis. Journal of Peptide Science 22 4–27 (2016).

78. Stawikowski, M. & Fields, G. B. Introduction to peptide synthesis. Curr. Protoc. Protein Sci. 69, 18–1 (2012).

79. Hayden, E. Y. et al. Identification of key regions and residues controlling Aβ folding and assembly. Sci. Rep. 7, (2017).

80. Seuma, M., Lehner, B. & Bolognesi, B. An atlas of amyloid aggregation: the impact of substitutions, insertions, deletions and truncations on amyloid beta fibril nucleation. Nat. Commun. 13, (2022).

81. Thacker, D. et al. The role of fibril structure and surface hydrophobicity in secondary nucleation of amyloid fibrils. Proc. Natl. Acad. Sci. U. S. A. 117 25272–25283 (2020).

82. Furniss, B. S. Vogel’s Textbook of Practical Organic Chemistry. (Pearson Education, 2011).

83. Hu, J. et al. Structural defects in amyloid-β fibrils drive secondary nucleation. Nat. Commun. 17, 1933 (2026).

84. Michaels, T. C. T. et al. Dynamics of oligomer populations formed during the aggregation of Alzheimer’s Aβ42 peptide. Nat. Chem. 12, 445–451 (2020).

85. Guerrero-Muñoz, M. J. et al. Amyloid-β oligomers as a template for secondary amyloidosis in Alzheimer’s disease. Neurobiol. Dis. 71, 14–23 (2014).

86. Lambert, M. P. et al. Diffusible, nonfibrillar ligands derived from Aβ1–42 are potent central nervous system neurotoxins. Neurobiology 95, 6448–6453 (1998).

87. Roher, A. E. et al. Morphology and toxicity of Aβ-(1-42) dimer derived from neuritic and vascular amyloid deposits of Alzheimer’s disease. Journal of Biological Chemistry 271, 20631–20635 (1996).

88. Walsh, D. M., Klyubin, I., Fadeeva, J. V., Rowan, M. J. & Selkoe, D. J. Amyloid-β oligomers: Their production, toxicity and therapeutic inhibition. Biochem. Soc. Trans. 30, 552–557 (2002).

89. Abedini, A., Singh, G. & Raleigh, D. P. Recovery and purification of highly aggregation-prone disulfide-containing peptides: Application to islet amyloid polypeptide. Anal. Biochem. 351, 181–186 (2006).

90. Mosconi, M. et al. Tau catalyzes amyloid-β aggregation and toxicity in a polymorph-dependent manner. 123, 2532775123 (2026).

91. Koloteva-Levine, N. et al. Amyloid particles facilitate surface-catalyzed cross-seeding by acting as promiscuous nanoparticles. Proc. Natl. Acad. Sci. U. S. A. 118, e2104148118 (2021).

92. Vaneyck, J., Segers-Nolten, I., Broersen, K. & Claessens, M. M. A. E. Cross-seeding of alpha-synuclein aggregation by amyloid fibrils of food proteins. Journal of Biological Chemistry 296, 100358 (2021).

93. Seubert, P. et al. Isolation and quantification of soluble Alzheimer’s β-peptide from biological fluids. Nature 359, 325–327 (1992).

94. Brinkmalm, A. et al. Fluid-based proteomics targeted on pathophysiological processes and pathologies in neurodegenerative diseases. J. Neurochem. 151, 417–434 (2019).

95. Cukalevski, R. et al. The Aβ40 and Aβ42 peptides self-assemble into separate homomolecular fibrils in binary mixtures but cross-react during primary nucleation. Chem. Sci. 6, 4215–4233 (2015).

96. Thacker, D., Bless, M., Barghouth, M., Zhang, E. & Linse, S. A Palette of Fluorescent Aβ42 Peptides Labelled at a Range of Surface-Exposed Sites. Int. J. Mol. Sci. 23, (2022).

97. Törnquist, M. et al. Secondary nucleation in amyloid formation. Chemical Communications 54, 8667–8684 (2018).

98. Cohen, S. I. A. et al. A molecular chaperone breaks the catalytic cycle that generates toxic Aβ oligomers. Nat. Struct. Mol. Biol. 22, 207–213 (2015).

99. Burt, A. et al. An image processing pipeline for electron cryo-tomography in RELION-5. FEBS Open Bio 14, 1788–1804 (2024).

100. Rohou, A. & Grigorieff, N. CTFFIND4: Fast and Accurate Defocus Estimation from Electron Micrographs. Journal of structural biology, 192, 216–221 (2015).

101. He, S. & Scheres, S. H. W. Helical reconstruction in RELION. J. Struct. Biol. 198, 163–176 (2017).

102. Mcdonald, J. C., Duffy, D. C., Anderson, J. R. & Chiu, D. T. Review General Fabrication of microfluidic systems in poly (dimethylsiloxane). Electrophoresis 21, 27–40 (2000).

103. Tan, S. H., Nguyen, N. T., Chua, Y. C. & Kang, T. G. Oxygen plasma treatment for reducing hydrophobicity of a sealed polydimethylsiloxane microchannel. Biomicrofluidics 4, 1–8 (2010).

104. Saar, K. L. et al. On-chip label-free protein analysis with downstream electrodes for direct removal of electrolysis products. Lab Chip 18, 162–170 (2018).

