## Supplementary Information for "Errors in peptide synthesis are a source of discrepancies in Aβ42 studies"

#### S1. Batch-to-batch differences in s2Aβ42 persist after multiple SEC rounds

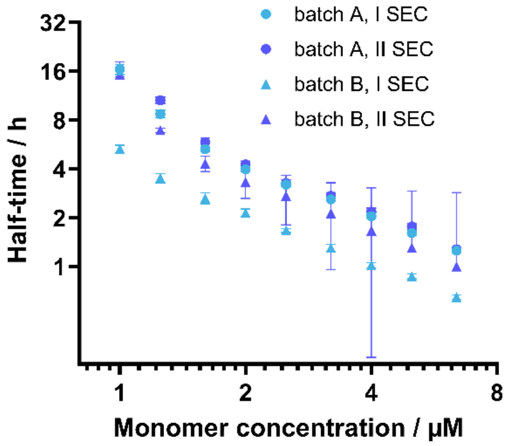


**Supplementary Fig. 1** Two different batches of s2Aβ42 were subjected to either 1 or 2 steps of SEC purification to evaluate the effect of purification steps in the kinetic behaviour of sAβ42. The half-time ($t_{1/2}$) of all s2Aβ42 replicates for each concentration was averaged and plotted as a function of initial monomer concentration. Light blue circles: batch A of s2Aβ42 subjected to one SEC; dark blue circles: batch A of s2Aβ42 subjected to two rounds of SEC; light blue triangles: batch B of s2Aβ42 subjected to one SEC; dark blue triangles: batch B of s2Aβ42 subjected to two rounds of SEC. Error bars represent standard deviation. Results show that, while different batches of the same synthetic protein differ in their aggregation behaviour, this difference is reduced by additional SEC purification steps.

**S2. AβM1-42 exhibits nearly identical kinetic behaviour to rAβ42**

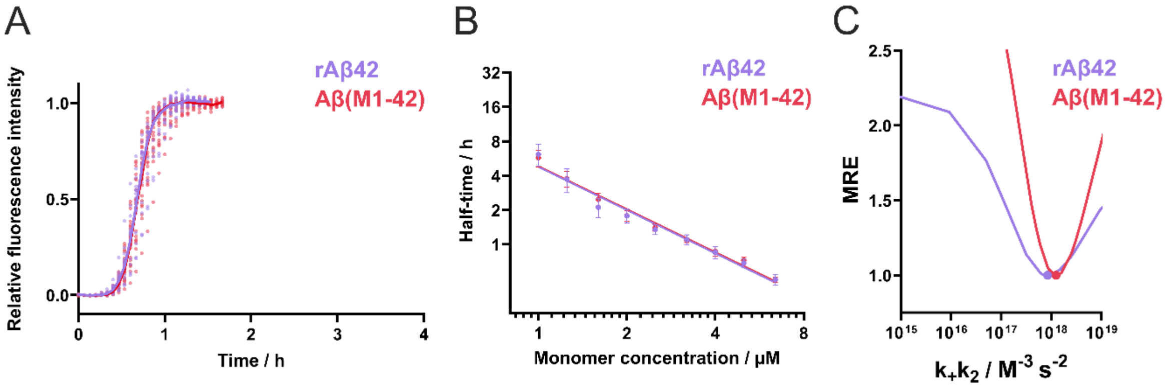


**Supplementary Fig. 2** Comparative study of rAβ42 and Aβ(M1-42). (A) Aggregation of rAβ42 (purple) and Aβ(M1-42) (red) at 5 μM. Data collected in 5 different experiments with 3-6 different replicates in each were put together and plotted as dots. The median of the obtained measurements for each protein is shown with a line. The x-axis is chosen for ease of comparison to Fig. 1a. (B) Initial monomer concentration of each experiment was plotted against the half-time ($t_{1/2}$) of the aggregation obtained for rAβ42 (purple) and Aβ(M1-42) (red). (C) Relative MRE as a function of $k_+k_2$ measured for rAβ42 (purple) and Aβ(M1-42) (red). Primary nucleus size ($n_c$) and secondary nucleus size ($n_2$) were fixed at 2.

**S3. Fitting of concentration-dependant aggregation kinetics of all peptides using AmyloFit.**

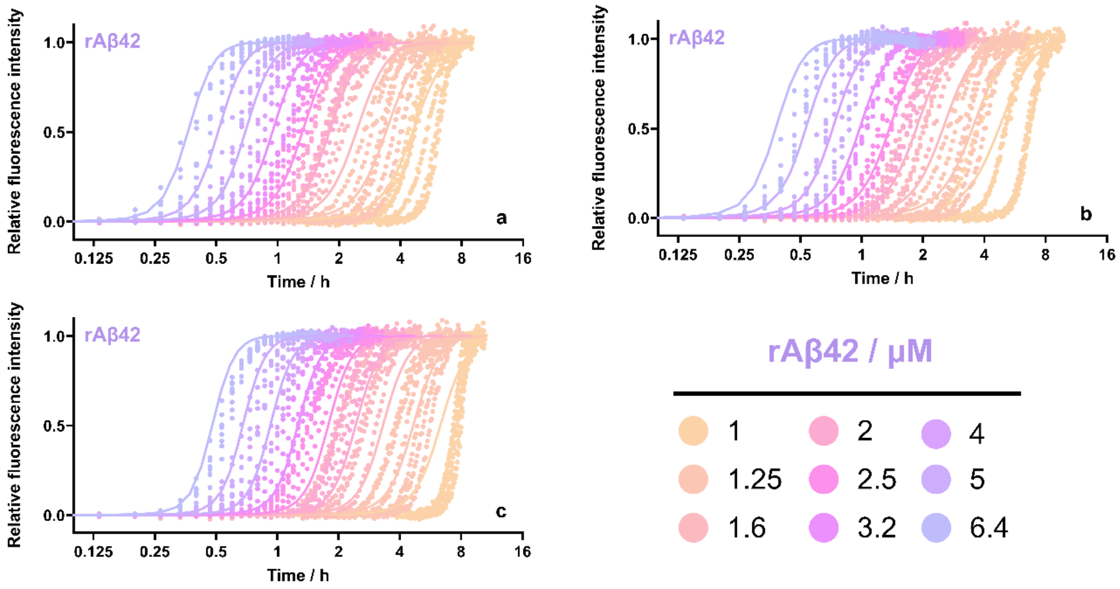


**Supplementary Fig. 3 Concentration-dependent** aggregation of rAβ42 (a-c). All proteins were studied using initial monomer concentrations between 1-6.4 µM, with data represented as dots. AmyloFit[1] was used to find the best global fit to each data set, while fixing primary nucleation reaction order ($n_c$) and secondary nucleation reaction order ($n_2$) to 2 (fit represented by lines), identifying secondary nucleation dominated model as the best fitting mechanism for all peptides.

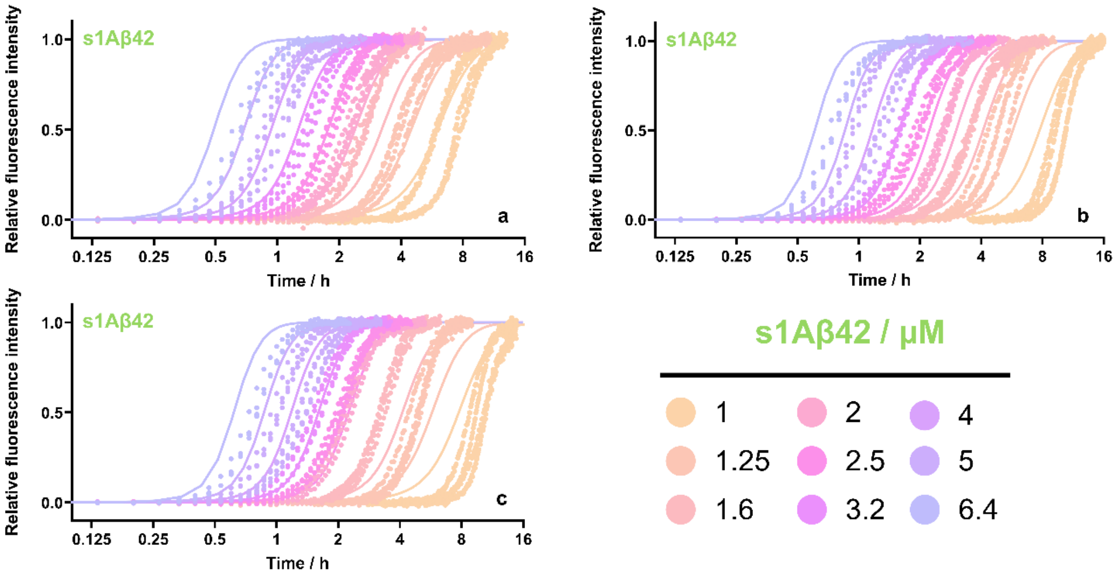


**Supplementary Fig. 4** Concentration dependent aggregation of s1Aβ42 (a-c). All proteins were studied using initial monomer concentrations between 1-6.4 μM, with data represented as dots. AmyloFit[1] was used to find the best global fit to each data set, while fixing primary nucleation reaction order ($n_c$) and secondary nucleation reaction order ($n_2$) to 2 (fit represented by lines), identifying secondary nucleation dominated model as the best fitting mechanism for all peptides.

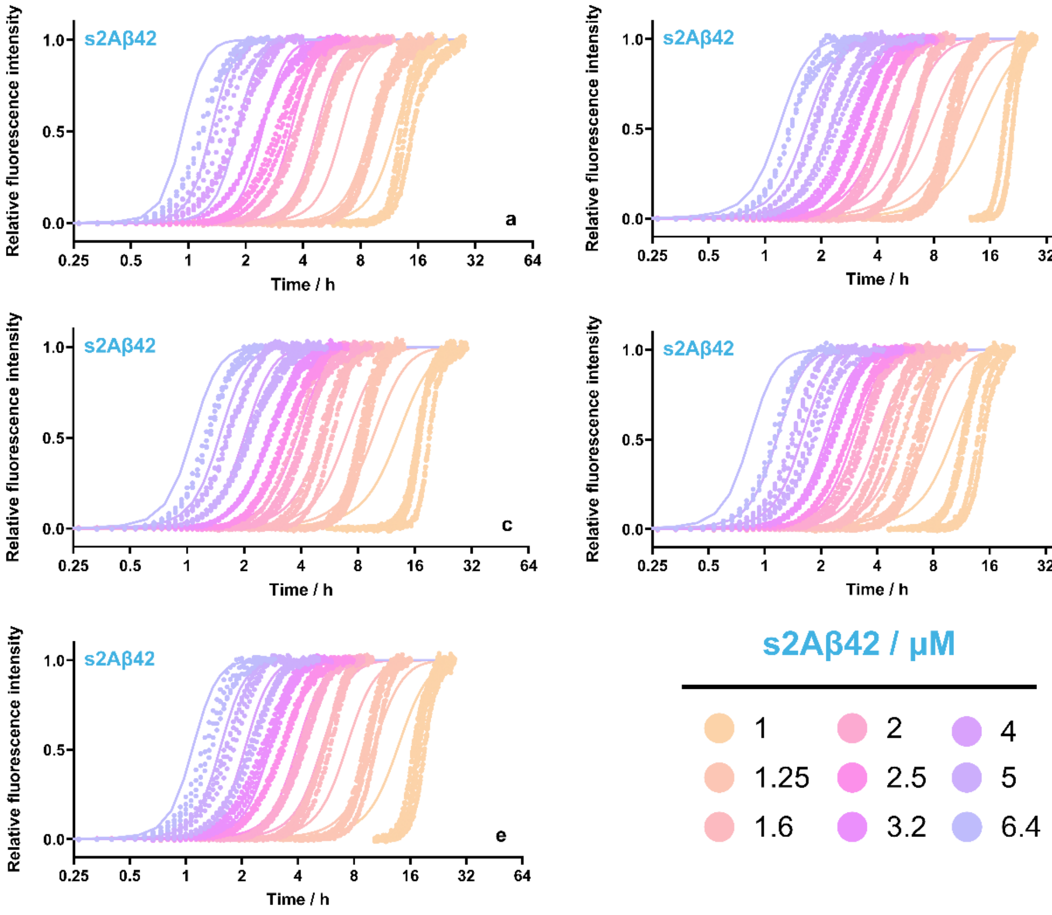


**Supplementary Fig. 5** Concentration dependent aggregation of s2Aβ42 (a-c). All proteins were studied using initial monomer concentrations between 1-6.4 µM, with data represented as dots. AmyloFit[1] was used to find the best global fit to each data set, while fixing primary nucleation reaction order ($n_c$) and secondary nucleation reaction order ($n_2$) to 2 (fit represented by lines), identifying secondary nucleation dominated model as the best fitting mechanism for all peptides.

#### S4. Comparison of rate constants and quality of fitting for all Aβ42 variants.

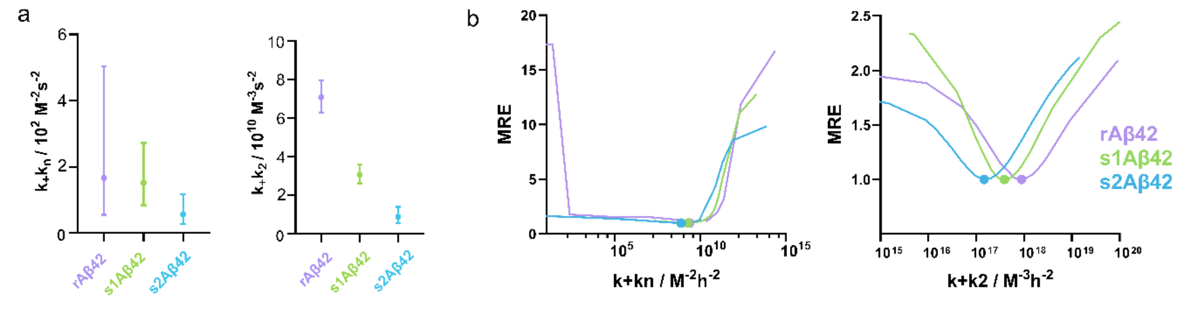


**Supplementary Fig. 6** The kinetic parameters $k_+k_n$ (a, left) and $k_+k_2$ (a, right) as obtained from the fits shown in Supplementary Fig. 3, 4 and 5. These values were measured independently for 3-5 different experiments for each protein. The data plotted in (a) represent the geometric mean and geometric standard deviation of $k_+k_n$ and $k_+k_2$ obtained for rAβ42 (purple), s1Aβ42 (green), and s2Aβ42 (blue). The full data obtained can be found in Supplementary Table 1. To analyse the reliability of the parameters, relative MRE was measured as a function of $k_+k_n$ (b, left) and $k_+k_2$ (b, right). The same fit used to obtain (a) was done, but fixing the value of either $k_+k_n$ (b, left) or k+k2 (B, right) and allowing the other one to vary as much as needed for the program to find the best fit to the data. The MRE values obtained for those fits are reported relative to the MRE of the best fit.

**Supplementary Table 1** $k_+k_n$, $k_+k_2$, MRE of fit and slope of double-logarithmic plot calculated for each experiment using AmyloFit [1]. Individual slopes were averaged and plotted on Fig. 1 with respective SD. For each peptide type, 3-5 independent replicates were performed, each consisting of 4-6 technical replicates across Aβ42 concentration range of 0.8 – 6.4 µM. The last two columns show the geometric mean of the kinetic parameters for each protein, with the geometric standard deviation between brackets.

| | No. | k+ kn ($M^{-2}$ $s^{-2}$) | k+ k2 ($M^{-3}$ $s^{-2}$) | MRE | Slope | k+ kn ($M^{-2}$ $s^{-2}$) | k+ k2 ($M^{-3}$ $s^{-2}$) |
|---|---|---|---|---|---|---|---|
| rAβ42 | 1 | 2.77E+02 | 8.1E+10 | 0.0247 | -1.209 | 1.67E+02 (3) | 7.09E+10 (1.1) |
| | 2 | 3.58E+02 | 6.76E+10 | 0.0296 | -1.201 | | |
| | 3 | 0.47E+02 | 6.53E+10 | 0.0358 | -1.331 | | |
| s1Aβ42 | 1 | 2.8E+02 | 3.65E+10 | 0.0125 | -1.166 | 1.52E+02 (1.8) | 3.08E+10 (1.2) |
| | 2 | 0.87E+02 | 3.01E+10 | 0.0227 | -1.274 | | |
| | 3 | 1.43E+02 | 2.66E+10 | 0.0303 | -1.215 | | |
| s2Aβ42 | 1 | 0.19E+02 | 1.54E+10 | 0.0169 | -1.298 | 0.57E+02 (2.1) | 0.89E+10 (1.6) |
| | 2 | 0.997E+02 | 0.48E+10 | 0.0474 | -1.303 | | |
| | 3 | 0.69E+02 | 0.75E+10 | 0.0407 | -1.262 | | |
| | 4 | 1.15E+02 | 1.21E+10 | 0.0339 | -1.283 | | |
| | 5 | 0.37E+02 | 0.85E+10 | 0.0397 | -1.293 | | |
| Aβ(M1-42) | 1 | 0.28E+02 | 9.72E+10 | 0.0138 | -1.312 | 0.55E+02 (3.7) | 9.87E+10 (1.2) |
| | 2 | 0.77E+02 | 9.65E+10 | 0.0133 | -1.244 | | |
| | 3 | 0.14E+02 | 0.12E+10 | 0.0209 | -1.298 | | |
| | 4 | 2.95E+02 | 8.26E+10 | 0.0267 | -1.195 | | |

**S5. Evaluating the contribution of individual microscopic rate constants to the difference between s1Aβ42 and rAβ42.**

While synthetic Aβ42 shows a clearly slower aggregation than its recombinant counterpart, the double logarithmic plot and fitting of kinetic models to the data suggest both rAβ42 and s1Aβ42 aggregate following a similar mechanism, one dominated by secondary nucleation. To elucidate the role of each individual microscopic step of the aggregation in the observed difference, we performed a series of experiments and calculations.

Firstly, to isolate the role of secondary nucleation, we used the chaperone Brichos, shown to block secondary nucleation by binding to defects on the fibril surface (Supplementary Fig. 7a) [2]. Interestingly, blocking secondary nucleation accentuated the difference between s1Aβ42 and rAβ42, indicating that a mechanism other than secondary nucleation plays a role in this difference. The original reaction being secondary nucleation dominated meant effects on other microscopic steps were masked, thus the bigger error in determination of $k_+k_n$ (Supplementary Fig. 6). Thus, by fitting the data obtained in the presence of Brichos to an elongation+primary nucleation model in AmyloFit[1], we obtained a more accurate comparison between the $k_+k_n$ of the two systems, that being that **rAβ42 has a $k_+k_n$ rate ~8.7 times bigger than that of s1Aβ42**. With that knowledge, we fitted the original data again using AmyloFit, but this time forcing the program to make the $k_+k_n$ of rAβ42 8.7 times larger than that of s1Aβ42. Doing this gave us a more informed comparison of their $k_+k_2$ values, which showed **rAβ42 to have 2.1 times higher $k_+k_2$ relative to s1Aβ42**.

Following that, we isolated the effect of elongation by setting up reactions where Aβ42 monomers were seeded with highly sonicated fibrils, creating conditions where elongation was the dominant aggregation mechanism (Supplementary Fig. 7b). Under these conditions, the elongation rate $k_+$ can be approximated from the initial slopes of the aggregation traces. More specifically, dividing the slope obtained with recombinant monomer by that obtained with synthetic monomer shows the difference in $k_+$ between the two systems. Given how the monomer dominates the seeding speed, calculating this with either seed type gave a similar value, that being that **rAβ42 shows a $k_+$ ~3.8 times bigger than that of s1Aβ42**.

Finally, using the relative differences obtained for $k_+k_n$, $k_+k_2$ and $k_+$ allowed us to calculate the difference between all the individual microscopic rate constants:

- **$k_+$ --> rAβ42 = 3.8*s1Aβ42**
- **$k_n$ --> rAβ42 = 2.3*s1Aβ42**
- **$k_2$ --> 1.8*rAβ42 = s1Aβ42**

Given that the differences between all the calculated rate constants are within a factor of four, no clear conclusions can be drawn from these results, other than that all three steps in the mechanism could be affected by the protein source.

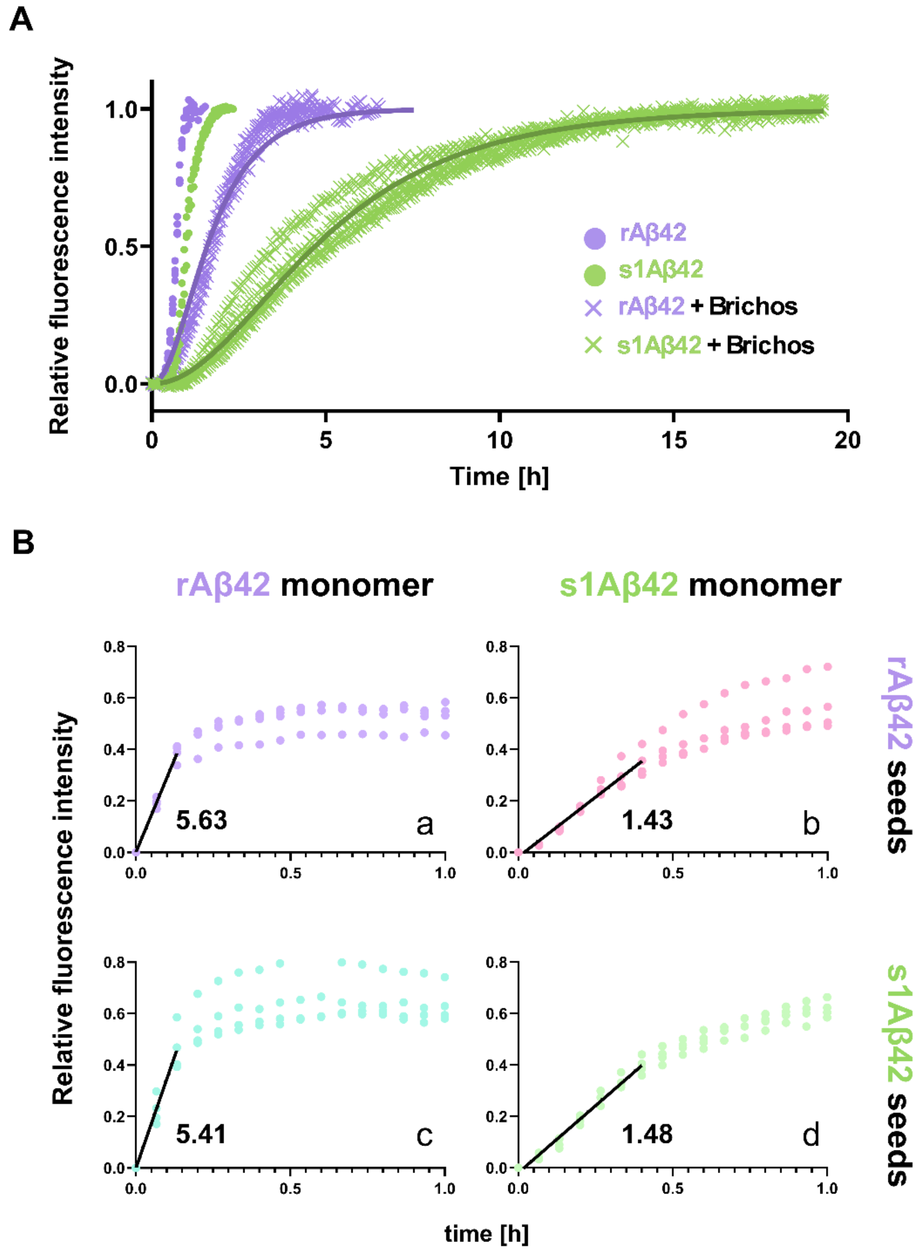


**Supplementary Fig. 7** (A) Aggregation of rAβ42 (purple) and s1Aβ42 (green) in the absence (dots) or presence (crosses) of Brichos chaperone. Each data point of 3 replicates is plotted as a symbol. The data obtained in the presence of Brichos was fitted to an elongation+primary nucleation model (setting the reaction orders $n_c = n_2 = 2$) using AmyloFit[1]. The best fit obtained, here represented by the lines, gave $k_+k_n$ values of $1.37x10^{10}$ and $1.58x10^{9}$ $M^{-2}$ $h^{-2}$ for rAβ42 and s1Aβ42, respectively. (B) Elongation dominated aggregation reactions. 8% sonicated fibrils were added to monomeric 3 µM Aβ42. Seeds were formed with rAβ42 (a, b) or s1Aβ42 (c, d) and mixed with monomeric rAβ42 (a, c) or s1Aβ42 (b, d). Data were plotted as dots. Lines show linear regression fit performed using GraphPad Prism version 11.0.2 for Windows, GraphPad Software, Boston, Massachusetts USA, www.graphpad.com. Numbers inside each graph correspond to the slope obtained for each experiment. When comparing the effect that the monomer has on the slope of the linear regime of the elongation-dominated aggregation, using rAβ42 seeds (a, b) and using s1Aβ42 seeds (c, d) led to differences of 3.94 and 3.65, respectively. This indicates $k_+$ being ~3.8 times bigger for rAβ42 than for s1Aβ42.

#### S6. NMR analysis of monomeric rAβ42 and s1Aβ42

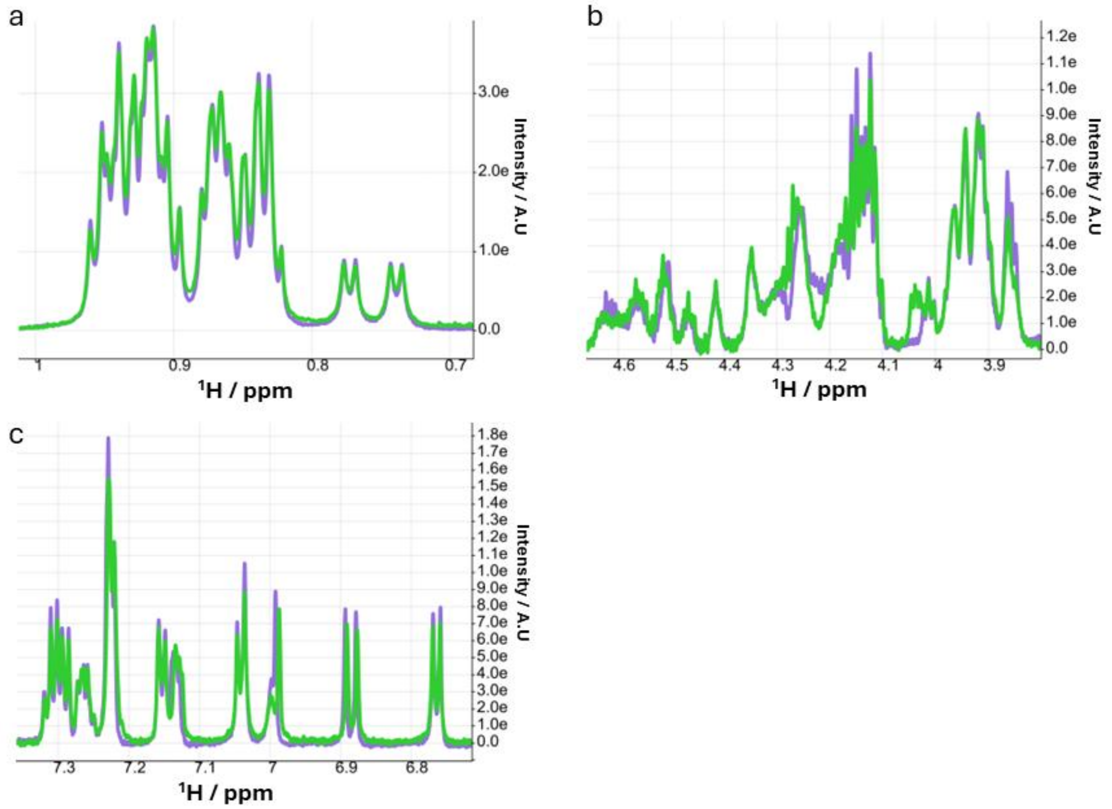


**Supplementary Fig. 8** 1D $^1$H NMR spectra of s1Aβ42 (green) and rAβ42 (purple), zoomed in on the methyl (a), Cα (b) and aromatic side chain (c) regions.

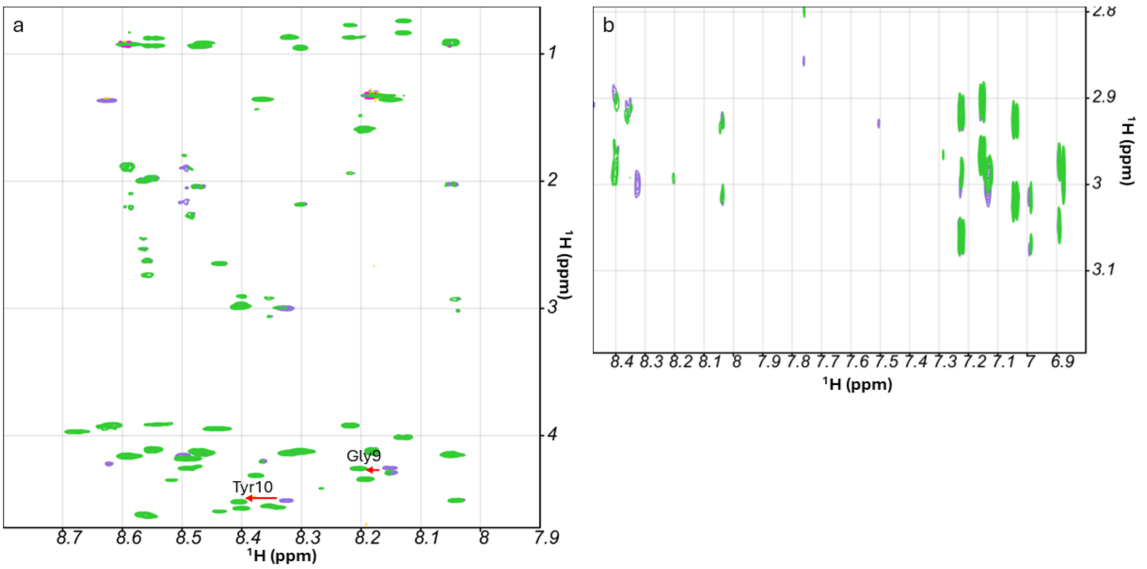


**Supplementary Fig. 9** (a) Section of a 2D $^1$H-$^1$H TOCSY spectrum recorded at 2°C for s1Aβ42 (green) and rAβ42 (purple). Red arrows depict a change in chemical shift corresponding to

Gly9 and Tyr10. (b) Section of a 2D $^{1}H$-$^{1}H$ NOESY spectrum recorded at 2°C for s1Aβ42 (green) and rAβ42 (purple).

#### S7. Seeding efficiency and cryo-EM imaging of s2Aβ42

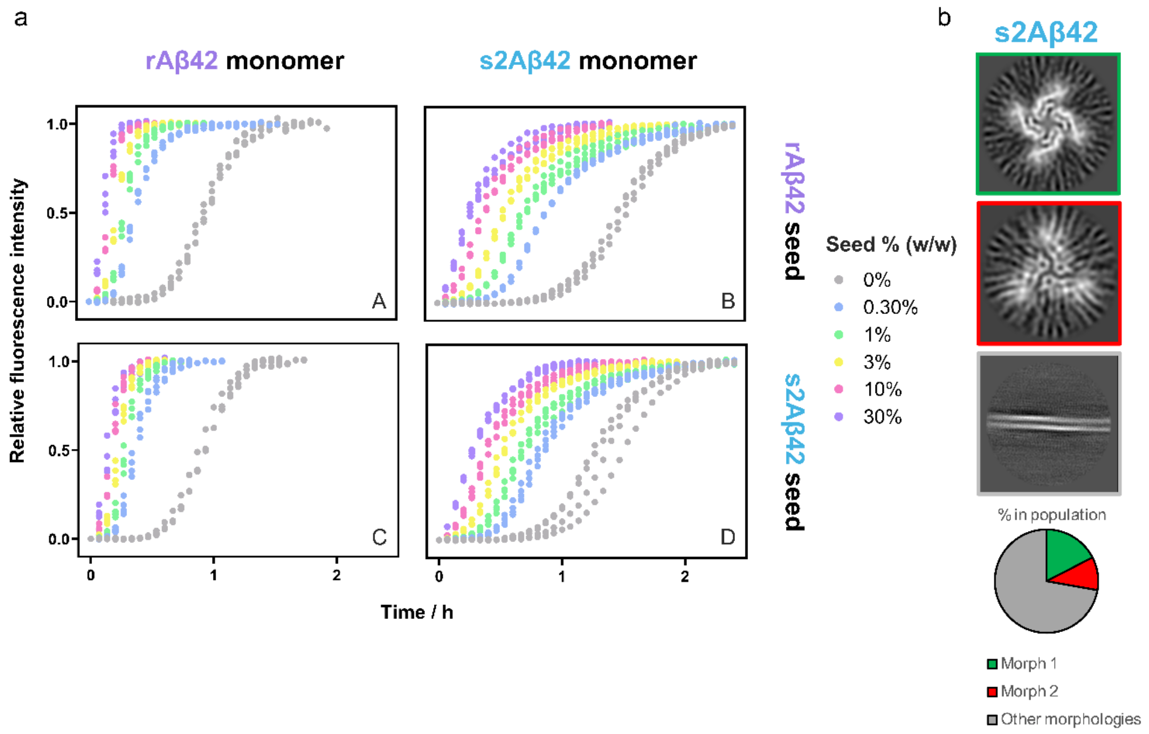


**Supplementary Fig. 10** (A) Seeded aggregation monitored using ThT fluorescence. 6 µM monomeric rAβ42 (a & c) or s2Aβ42 (b & d) were aggregated in the presence of seeds made with either rAβ42 (a & b) or s2Aβ42 (c & d) at w/w concentrations of 0, 0.3, 1, 3, 10 and 30% (grey, blue, green, yellow, pink, and purple, respectively). (D) Representative cryo-EM 3D classes identified for s2Aβ42 fibrils (left) and the relative abundance of each morphology within the classified fibril population of isolated fibrils outside large clusters (right).

#### S8. MS detection of deletion and duplication variants of Aβ42.

**Supplementary Table 2.** Deletion variants identified for s1Aβ42 and rAβ42. MS data of s1Aβ42 and rAβ42 samples were analysed using a custom in-house FASTA file containing 42 unique duplication variants of the Aβ42 sequence. Using the integration of precursor MS1 Extracted Ion Chromatograms, each detected peptide's 280 nm peak intensity was measured. For each identified deletion variant, the same analysis was done for the WT equivalent of that peptide detected in the same sample. Finally, the fraction of peptides detected with deletion was calculated by dividing the intensity of the WT peptide peak by the sum of the intensity of the WT and deletion peaks. An example MS/MS spectrum of each of the peptides in the table can be seen in S12. Data plotted in Fig. 4a, left.

| Amino acid | Detected WT sequence | Detected **DELETION** sequence | Synthetic (s1Aβ42) | | | Recombinant (rAβ42) | | |
|---|---|---|---|---|---|---|---|---|
| | | | Peak intensity (a.u.) | | Fraction of peptides detected | Peak intensity (a.u.) | | Fraction of peptides detected |
| | | | WT | DELETION | with **DELETION** (%) | WT | DELETION | with **DELETION** (%) |
| H(13/14) | HDSGYEVHHQK | HDSGYEVHQK | 292208 | 6069 | 2,03 | 45170 | Not detected | - |
| K16 | HDSGYEVHHQK | HDSGYEVHHQLVFFAEDVGSNK | 292208 | 12360 | 4,06 | 45170 | Not detected | - |
| V18 | LVFFAEDVGSNK | LFFAEDVGSNK | 1321000 | 32960 | 2,43 | 295600 | 358 | 0,12 |
| F(19/20) | LVFFAEDVGSNK | LVFAEDVGSNK | 1321000 | 14740 | 1,10 | 295600 | 198 | 0,07 |
| A21 | LVFFAEDVGSNK | LVFFEDVGSNK | 1321000 | 8311 | 0,63 | 295600 | 280 | 0,09 |
| G25 | LVFFAEDVGSNK | LVFFAEDVSNK | 1321000 | 31720 | 2,34 | 295600 | 1377 | 0,46 |
| S26 | LVFFAEDVGSNK | LVFFAEDVGNK | 1321000 | 19510 | 1,46 | 295600 | 1767 | 0,59 |
| N27 | LVFFAEDVGSNK | LVFFAEDVGSK | 1321000 | 72540 | 5,21 | 295600 | 3221 | 1,08 |

**Supplementary Table 3.** Duplication variants identified for s1Aβ42 and rAβ42. MS data of s1Aβ42 and rAβ42 samples were analysed using a custom in-house FASTA file containing 42 unique duplication variants of the Aβ42 sequence. Using the integration of precursor MS1 Extracted Ion Chromatograms, each detected peptide's 280 nm peak intensity was measured. For each identified duplication variant, the same analysis was done for the WT equivalent of that peptide detected for the same sample. Finally, the fraction of peptides detected with duplication was calculated by dividing the intensity of the WT peptide peak by the sum of the intensity of the WT and duplication peaks. An example MS/MS spectrum of each of the peptides in the table can be seen in S12. Data plotted in Fig. 4a, right.

| Amino acid | Detected WT sequence | Detected **DUPLICATION** sequence | Synthetic (s1Aβ42) | | | Recombinant (rAβ42) | | |
|---|---|---|---|---|---|---|---|---|
| | | | Peak intensity (a.u.) | | Fraction of peptides detected | Peak intensity (a.u.) | | Fraction of peptides detected |
| | | | WT | DUPLICATION | with **DUPLICATION** (%) | WT | DUPLICATION | with **DUPLICATION** (%) |
| V12 | HDSGYEVHHQK | HDSGYEVVHHQK | 292208 | 8944 | 2,97 | 45170 | Not detected | - |
| L17 | LVFFAEDVGSNK | LLVFFAEDVGSNK | 1321000 | 1913 | 0,14 | 295600 | Not detected | - |
| V18 | LVFFAEDVGSNK | LVVFFAEDVGSNK | 1321000 | 678 | 0,05 | 295600 | Not detected | - |
| F(19/20) | LVFFAEDVGSNK | LVFFFAEDVGSNK | 1321000 | 1343 | 0,10 | 295600 | Not detected | - |
| A21 | LVFFAEDVGSNK | LVFFAAEDVGSNK | 1321000 | 29106 | 2,16 | 295600 | Not detected | - |
| V24 | LVFFAEDVGSNK | LVFFAEDVVGSNK | 1321000 | 4222 | 0,32 | 295600 | 176 | 0,06 |
| G25 | LVFFAEDVGSNK | LVFFAEDVGGSNK | 1321000 | 7654 | 0,58 | 295600 | 357 | 0,12 |
| N27 | LVFFAEDVGSNK | LVFFAEDVGSNNK | 1321000 | 6714 | 0,51 | 295600 | Not detected | - |

#### S9. Aggregation of Aβ42 used for the oligomer population measurement via μFFE

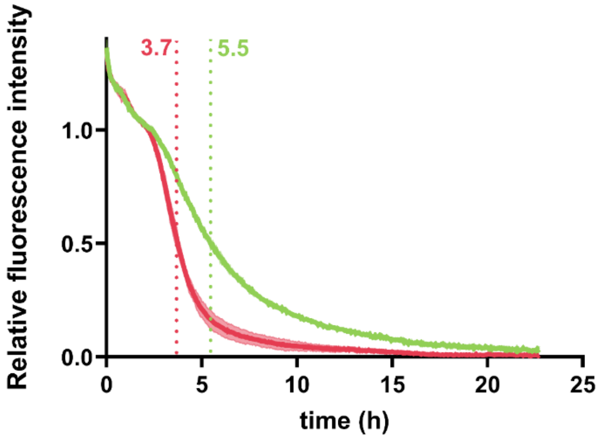


**Supplementary Fig. 11** Aggregation kinetics of 2 μM Aβ(M1-42) (red) and s1Aβ42 (green) in a mixture with Alexa 488 labelled S8C-Aβ(M1-42) at a 3.5:1 ratio. The aggregation is followed by monitoring the quenching of the fluorescence of Alexa 488 upon aggregation. Given that the lag phase of the aggregation is not flat when monitored with fluorescence quenching, the fluorescence intensity is normalised to the start of the exponential phase rather than the fluorescence at t = 0. Samples were collected at $t_{1/2}$ for Aβ(M1-42) (3.7 h) and s1Aβ42 (5.5 h) and analysed with μFFE as indicated in the methods section.

**S10. ThT fluorescence intensity at plateau of aggregation indicates negligible concentration errors for s1Aβ42 relative to rAβ42.**

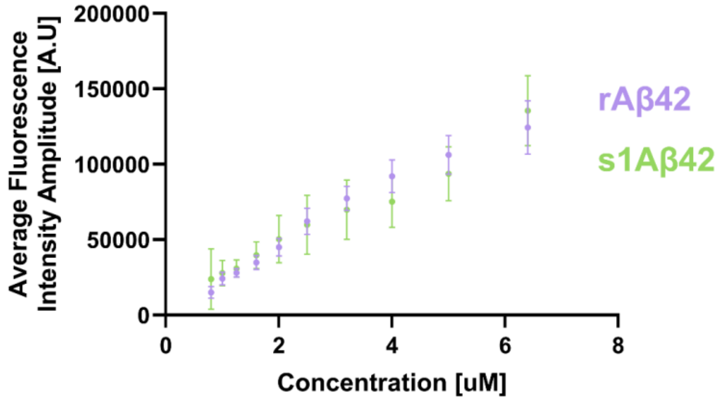


**Supplementary Fig. 12** End-point fluorescence intensity of rAβ42 (purple) and s1Aβ42 (green) as a function of initial monomer concentration. Data were collected across 3 to 5 experiments, with 4-6 replicates per concentration per experiment. Data are plotted as the median of the measurements for each concentration, with error bars representing standard deviation.

#### S11. HPLC data analysis of the soluble Aβ42 fraction

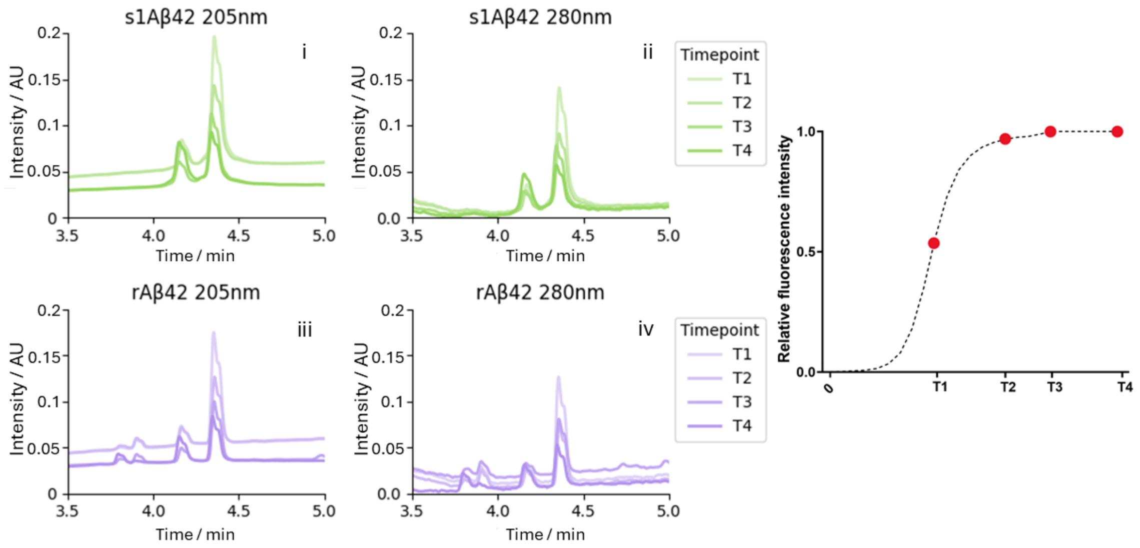


**Supplementary Fig. 13** Chromatograms from the HPLC-MS experiments at 205 nm (I, iii) and 280 nm (ii, iv). A single replicate from each time point (indicated on the right) is shown for s1Aβ42 (green) and rAβ42 (purple). The peak at approximately 4.3 min was identified as Aβ42 and its area was used to calculate the Aβ42 concentration (Fig. 2b).

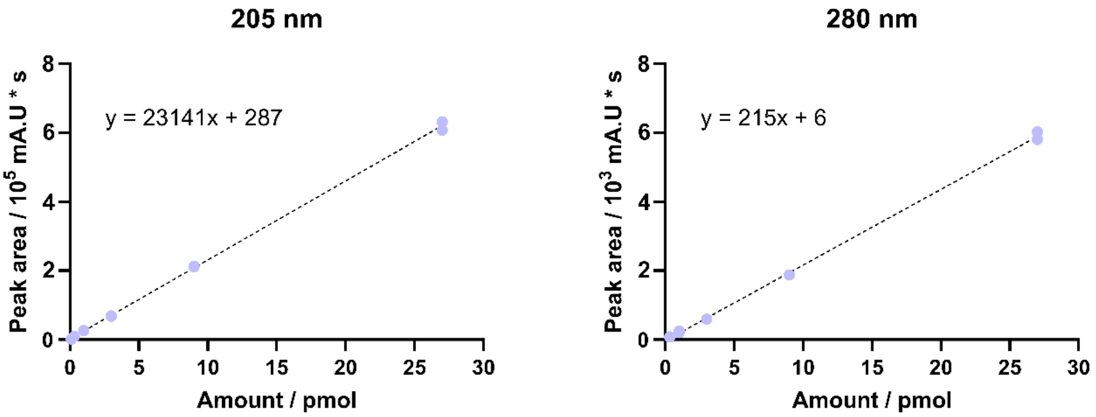


**Supplementary Fig. 14** Calibration curves for absorbance at 205 nm and 280 nm in HPLC-MS. The curve was constructed with a weighted (1/X) least-square linear regression. Measurements on standard samples (N =2) of recombinant Aβ(M1-42) were used as calibration points. Sample concentration was obtained by dividing the molar amount by the injected sample volume.

#### S12. MS/MS spectra of each peptide identification of rAβ42 and s1Aβ42 used for the MS analysis of imperfections

Pure monomeric rAβ42 and s1Aβ42 were digested with trypsin overnight and analysed on a quadrupole time-of-flight mass spectrometer. Selected precursor peptides were fragmented to generate MS/MS spectra. The mass of the precursor peptides, as well as the b and y ions (corresponding to N- and C-terminal fragments, respectively) formed upon their fragmentation, were compared with a custom-made database with duplication and deletion variants of the Aβ42 sequence.

This section contains the MS/MS spectra (and corresponding b and y ions) of the peptides used to analyse the propensity of duplication and deletion variants of rAβ42 and s1Aβ42, represented in Fig. 4 and Supplementary Tables 2 and 3.

### WT Aβ42 peptide detections

## In s1Aβ42

#### HDSGYEVHHQK

MS/MS Fragmentation of **HDSGYEVHHQK**

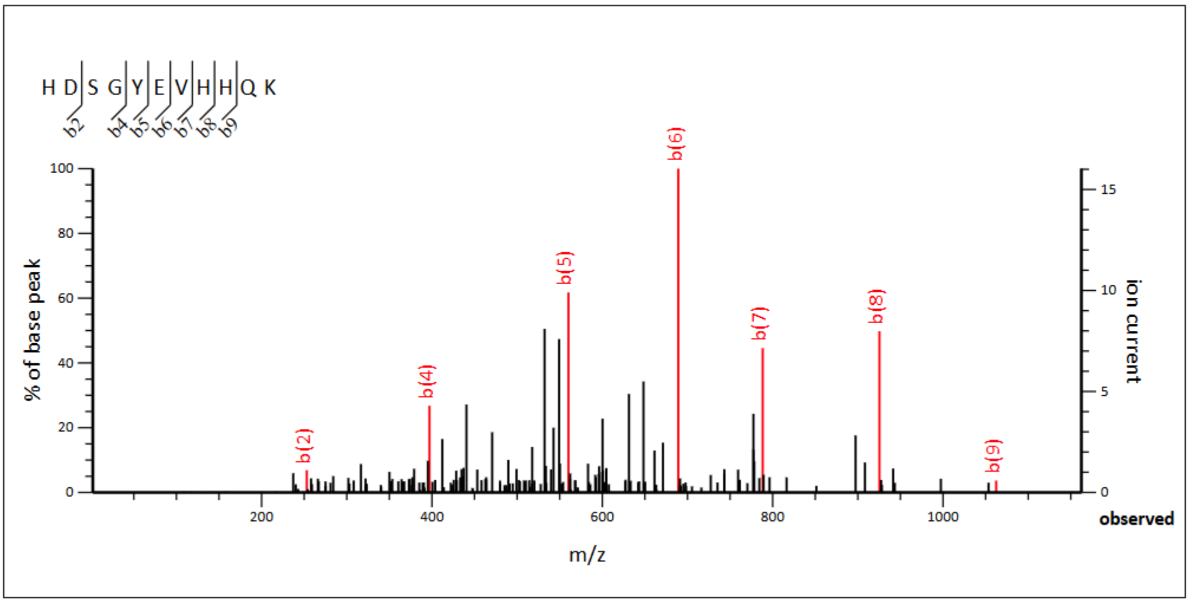


**Monoisotopic mass of neutral peptide Mr(calc):** 1335.5956
**Ions Score:** 55 **Expect:** 2.9e-06
**Matches :** 7/90 fragment ions using 10 most intense peaks

| # | b | $b^{++}$ | $b^{*}$ | $b^{*++}$ | $b^{0}$ | $b^{0++}$ | Seq. | y | $y^{++}$ | $y^{*}$ | $y^{*++}$ | $y^{0}$ | $y^{0++}$ | # |
|---|---|---|---|---|---|---|---|---|---|---|---|---|---|---|
| 1 | 138.0662 | 69.5367 | | | | | H | | | | | | | 11 |
| 2 | ***253.0931*** | 127.0502 | | | 235.0826 | 118.0449 | D | 1199.5440 | 600.2756 | 1182.5174 | 591.7624 | 1181.5334 | 591.2703 | 10 |
| 3 | 340.1252 | 170.5662 | | | 322.1146 | 161.5609 | S | 1084.5170 | 542.7622 | 1067.4905 | 534.2489 | 1066.5065 | 533.7569 | 9 |
| 4 | ***397.1466*** | 199.0769 | | | 379.1361 | 190.0717 | G | 997.4850 | 499.2461 | 980.4585 | 490.7329 | 979.4744 | 490.2409 | 8 |
| 5 | ***560.2100*** | 280.6086 | | | 542.1994 | 271.6033 | Y | 940.4635 | 470.7354 | 923.4370 | 462.2221 | 922.4530 | 461.7301 | 7 |
| 6 | ***689.2525*** | 345.1299 | | | 671.2420 | 336.1246 | E | 777.4002 | 389.2037 | 760.3737 | 380.6905 | 759.3896 | 380.1985 | 6 |
| 7 | ***788.3210*** | 394.6641 | | | 770.3104 | 385.6588 | V | 648.3576 | 324.6824 | 631.3311 | 316.1692 | | | 5 |
| 8 | ***925.3799*** | 463.1936 | | | 907.3693 | 454.1883 | H | 549.2892 | 275.1482 | 532.2627 | 266.6350 | | | 4 |
| 9 | ***1062.4388*** | 531.7230 | | | 1044.4282 | 522.7177 | H | 412.2303 | 206.6188 | 395.2037 | 198.1055 | | | 3 |
| 10 | 1190.4974 | 595.7523 | 1173.4708 | 587.2390 | 1172.4868 | 586.7470 | Q | 275.1714 | 138.0893 | 258.1448 | 129.5761 | | | 2 |
| 11 | | | | | | | K | 147.1128 | 74.0600 | 130.0863 | 65.5468 | | | 1 |

#### LVFFAEDVGSNK

MS/MS Fragmentation of **LVFFAEDVGSNK**

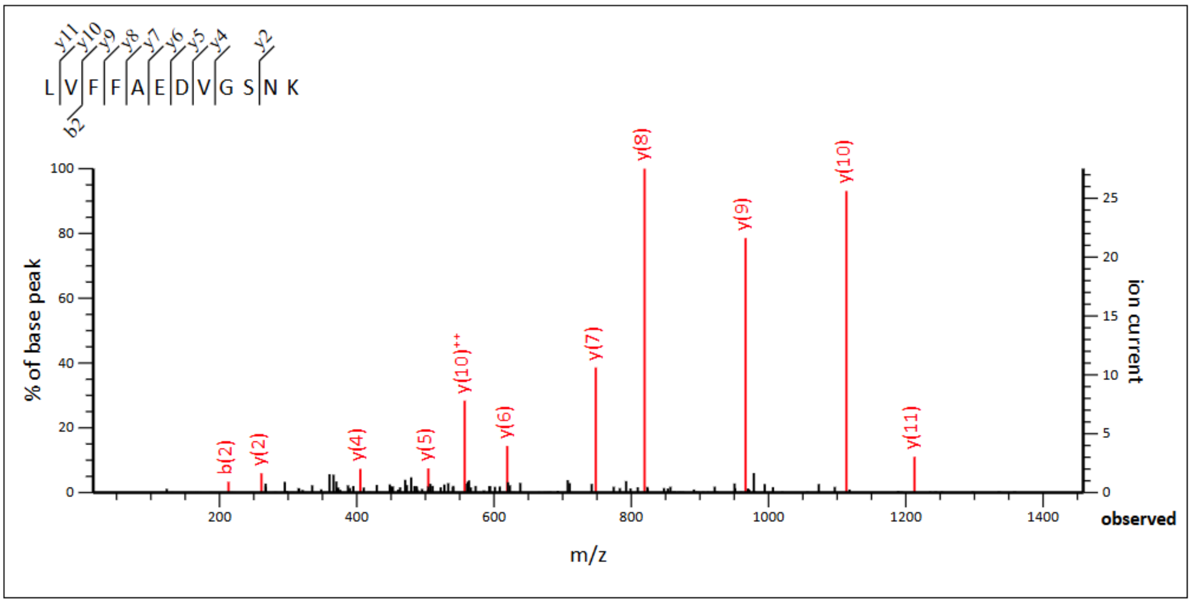


**Monoisotopic mass of neutral peptide Mr(calc):** 1324.6663
**Ions Score:** 98 **Expect:** 1.7e-10
**Matches :** 11/98 fragment ions using 12 most intense peaks

| # | b | $b^{++}$ | b* | $b^{*++}$ | $b^0$ | $b^{0++}$ | Seq. | y | $y^{++}$ | y* | $y^{*++}$ | $y^0$ | $y^{0++}$ | # |
|---|---|---|---|---|---|---|---|---|---|---|---|---|---|---|
| 1 | 114.0913 | 57.5493 | | | | | L | | | | | | | 12 |
| 2 | 213.1598 | 107.0835 | | | | | V | 1212.5895 | 606.7984 | 1195.5630 | 598.2851 | 1194.5790 | 597.7931 | 11 |
| 3 | 360.2282 | 180.6177 | | | | | F | 1113.5211 | 557.2642 | 1096.4946 | 548.7509 | 1095.5106 | 548.2589 | 10 |
| 4 | 507.2966 | 254.1519 | | | | | F | 966.4527 | 483.7300 | 949.4262 | 475.2167 | 948.4421 | 474.7247 | 9 |
| 5 | 578.3337 | 289.6705 | | | | | A | 819.3843 | 410.1958 | 802.3577 | 401.6825 | 801.3737 | 401.1905 | 8 |
| 6 | 707.3763 | 354.1918 | | | 689.3657 | 345.1865 | E | 748.3472 | 374.6772 | 731.3206 | 366.1639 | 730.3366 | 365.6719 | 7 |
| 7 | 822.4032 | 411.7053 | | | 804.3927 | 402.7000 | D | 619.3046 | 310.1559 | 602.2780 | 301.6427 | 601.2940 | 301.1506 | 6 |
| 8 | 921.4716 | 461.2395 | | | 903.4611 | 452.2342 | V | 504.2776 | 252.6425 | 487.2511 | 244.1292 | 486.2671 | 243.6372 | 5 |
| 9 | 978.4931 | 489.7502 | | | 960.4825 | 480.7449 | G | 405.2092 | 203.1082 | 388.1827 | 194.5950 | 387.1987 | 194.1030 | 4 |
| 10 | 1065.5251 | 533.2662 | | | 1047.5146 | 524.2609 | S | 348.1878 | 174.5975 | 331.1612 | 166.0842 | 330.1772 | 165.5922 | 3 |
| 11 | 1179.5681 | 590.2877 | 1162.5415 | 581.7744 | 1161.5575 | 581.2824 | N | 261.1557 | 131.0815 | 244.1292 | 122.5682 | | | 2 |
| 12 | | | | | | | K | 147.1128 | 74.0600 | 130.0863 | 65.5468 | | | 1 |

#### GAIIGLMVGGVVIA

MS/MS Fragmentation of **GAIIGLMVGGVVIA**

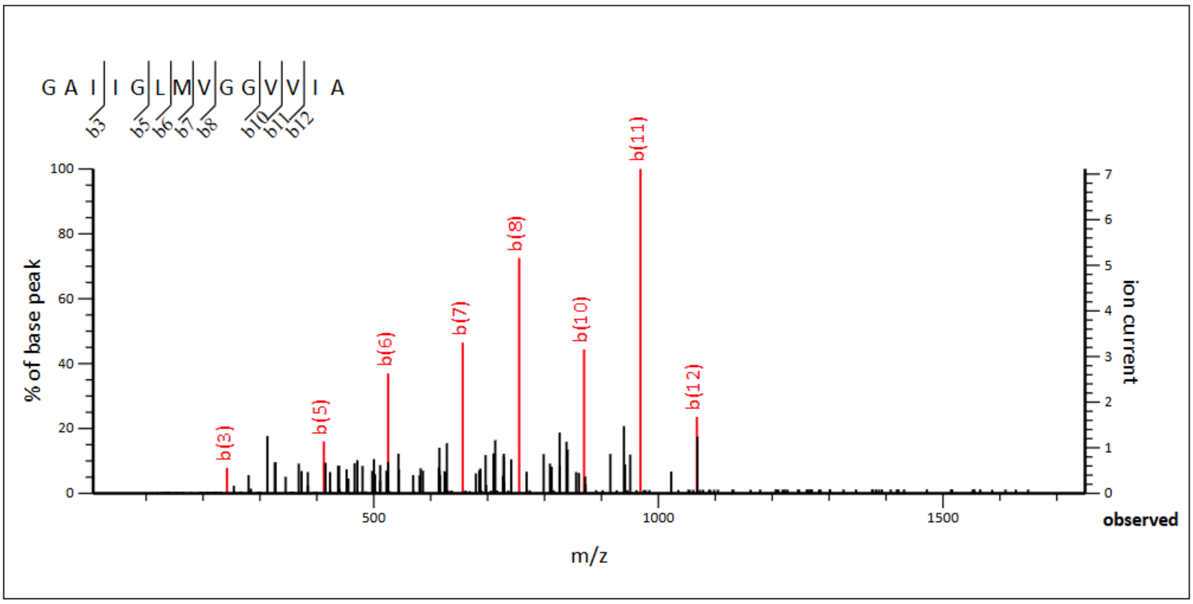


**Monoisotopic mass of neutral peptide Mr(calc):** 1268.7526
**Ions Score:** 63 **Expect:** 4.8e-07
**Matches :** 8/52 fragment ions using 12 most intense peaks

| # | b | $b^{++}$ | Seq. | y | $y^{++}$ | # |
|---|---|---|---|---|---|---|
| 1 | 58.0287 | 29.5180 | G | | | 14 |
| 2 | 129.0659 | 65.0366 | A | 1212.7384 | 606.8729 | 13 |
| 3 | *242.1499* | 121.5786 | I | 1141.7013 | 571.3543 | 12 |
| 4 | 355.2340 | 178.1206 | I | 1028.6173 | 514.8123 | 11 |
| 5 | *412.2554* | 206.6314 | G | 915.5332 | 458.2702 | 10 |
| 6 | *525.3395* | 263.1734 | L | 858.5117 | 429.7595 | 9 |
| 7 | *656.3800* | 328.6936 | M | 745.4277 | 373.2175 | 8 |
| 8 | *755.4484* | 378.2278 | V | 614.3872 | 307.6972 | 7 |
| 9 | 812.4699 | 406.7386 | G | 515.3188 | 258.1630 | 6 |
| 10 | *869.4913* | 435.2493 | G | 458.2973 | 229.6523 | 5 |
| 11 | *968.5598* | 484.7835 | V | 401.2758 | 201.1416 | 4 |
| 12 | *1067.6282* | 534.3177 | V | 302.2074 | 151.6074 | 3 |
| 13 | 1180.7122 | 590.8598 | I | 203.1390 | 102.0731 | 2 |
| 14 | | | A | 90.0550 | 45.5311 | 1 |

#### In rAβ42

#### HDSGYEVHHQK

MS/MS Fragmentation of **HDSGYEVHHQK**

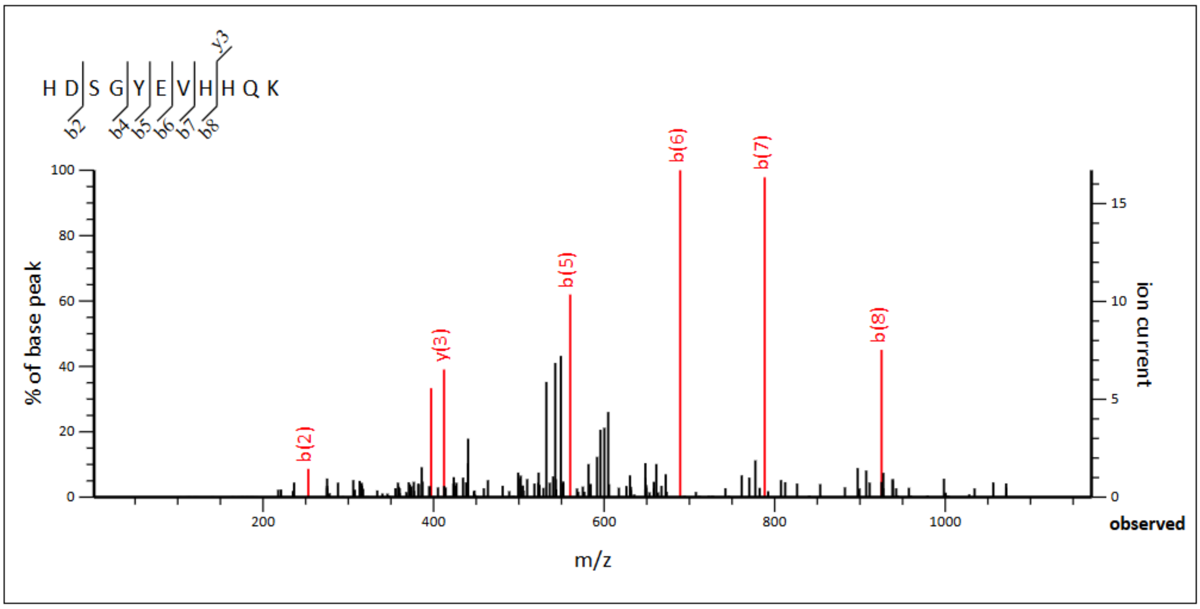


**Monoisotopic mass of neutral peptide Mr(calc):** 1335.5956
**Ions Score:** 45 **Expect:** 3e-05
**Matches :** 7/90 fragment ions using 10 most intense peaks

| # | b | $b^{++}$ | b* | $b^{*++}$ | $b^0$ | $b^{0++}$ | Seq. | y | $y^{++}$ | y* | $y^{*++}$ | $y^0$ | $y^{0++}$ | # |
|---|---|---|---|---|---|---|---|---|---|---|---|---|---|---|
| 1 | 138.0662 | 69.5367 | | | | | H | | | | | | | 11 |
| 2 | 253.0931 | 127.0502 | | | 235.0826 | 118.0449 | D | 1199.5440 | 600.2756 | 1182.5174 | 591.7624 | 1181.5334 | 591.2703 | 10 |
| 3 | 340.1252 | 170.5662 | | | 322.1146 | 161.5609 | S | 1084.5170 | 542.7622 | 1067.4905 | 534.2489 | 1066.5065 | 533.7569 | 9 |
| 4 | 397.1466 | 199.0769 | | | 379.1361 | 190.0717 | G | 997.4850 | 499.2461 | 980.4585 | 490.7329 | 979.4744 | 490.2409 | 8 |
| 5 | 560.2100 | 280.6086 | | | 542.1994 | 271.6033 | Y | 940.4635 | 470.7354 | 923.4370 | 462.2221 | 922.4530 | 461.7301 | 7 |
| 6 | 689.2525 | 345.1299 | | | 671.2420 | 336.1246 | E | 777.4002 | 389.2037 | 760.3737 | 380.6905 | 759.3896 | 380.1985 | 6 |
| 7 | 788.3210 | 394.6641 | | | 770.3104 | 385.6588 | V | 648.3576 | 324.6824 | 631.3311 | 316.1692 | | | 5 |
| 8 | 925.3799 | 463.1936 | | | 907.3693 | 454.1883 | H | 549.2892 | 275.1482 | 532.2627 | 266.6350 | | | 4 |
| 9 | 1062.4388 | 531.7230 | | | 1044.4282 | 522.7177 | H | 412.2303 | 206.6188 | 395.2037 | 198.1055 | | | 3 |
| 10 | 1190.4974 | 595.7523 | 1173.4708 | 587.2390 | 1172.4868 | 586.7470 | Q | 275.1714 | 138.0893 | 258.1448 | 129.5761 | | | 2 |
| 11 | | | | | | | K | 147.1128 | 74.0600 | 130.0863 | 65.5468 | | | 1 |

### LVFFAEDVGSNK

MS/MS Fragmentation of **LVFFAEDVGSNK**

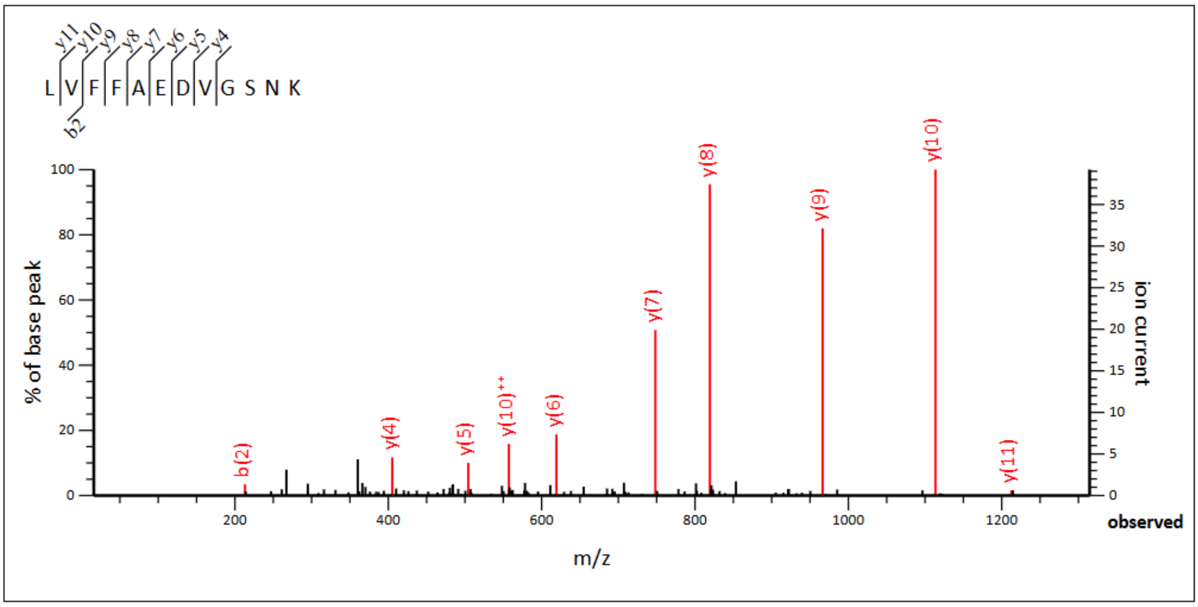


**Monoisotopic mass of neutral peptide Mr(calc):** 1324.6663
**Ions Score:** 83 **Expect:** 4.5e-09
**Matches :** 10/98 fragment ions using 11 most intense peaks

| # | b | $b^{++}$ | b* | $b^{*++}$ | $b^{0}$ | $b^{0++}$ | Seq. | y | $y^{++}$ | y* | $y^{*++}$ | $y^{0}$ | $y^{0++}$ | # |
|---|---|---|---|---|---|---|---|---|---|---|---|---|---|---|
| 1 | 114.0913 | 57.5493 | | | | | L | | | | | | | 12 |
| 2 | **213.1598** | 107.0835 | | | | | V | ***1212.5895*** | 606.7984 | 1195.5630 | 598.2851 | 1194.5790 | 597.7931 | 11 |
| 3 | 360.2282 | 180.6177 | | | | | F | ***1113.5211*** | **557.2642** | 1096.4946 | 548.7509 | 1095.5106 | 548.2589 | 10 |
| 4 | 507.2966 | 254.1519 | | | | | F | ***966.4527*** | 483.7300 | 949.4262 | 475.2167 | 948.4421 | 474.7247 | 9 |
| 5 | 578.3337 | 289.6705 | | | | | A | ***819.3843*** | 410.1958 | 802.3577 | 401.6825 | 801.3737 | 401.1905 | 8 |
| 6 | 707.3763 | 354.1918 | | | 689.3657 | 345.1865 | E | ***748.3472*** | 374.6772 | 731.3206 | 366.1639 | 730.3366 | 365.6719 | 7 |
| 7 | 822.4032 | 411.7053 | | | 804.3927 | 402.7000 | D | ***619.3046*** | 310.1559 | 602.2780 | 301.6427 | 601.2940 | 301.1506 | 6 |
| 8 | 921.4716 | 461.2395 | | | 903.4611 | 452.2342 | V | ***504.2776*** | 252.6425 | 487.2511 | 244.1292 | 486.2671 | 243.6372 | 5 |
| 9 | 978.4931 | 489.7502 | | | 960.4825 | 480.7449 | G | ***405.2092*** | 203.1082 | 388.1827 | 194.5950 | 387.1987 | 194.1030 | 4 |
| 10 | 1065.5251 | 533.2662 | | | 1047.5146 | 524.2609 | S | 348.1878 | 174.5975 | 331.1612 | 166.0842 | 330.1772 | 165.5922 | 3 |
| 11 | 1179.5681 | 590.2877 | 1162.5415 | 581.7744 | 1161.5575 | 581.2824 | N | 261.1557 | 131.0815 | 244.1292 | 122.5682 | | | 2 |
| 12 | | | | | | | K | 147.1128 | 74.0600 | 130.0863 | 65.5468 | | | 1 |

#### GAIIGLMVGGVVIA

MS/MS Fragmentation of **GAIIGLMVGGVVIA**

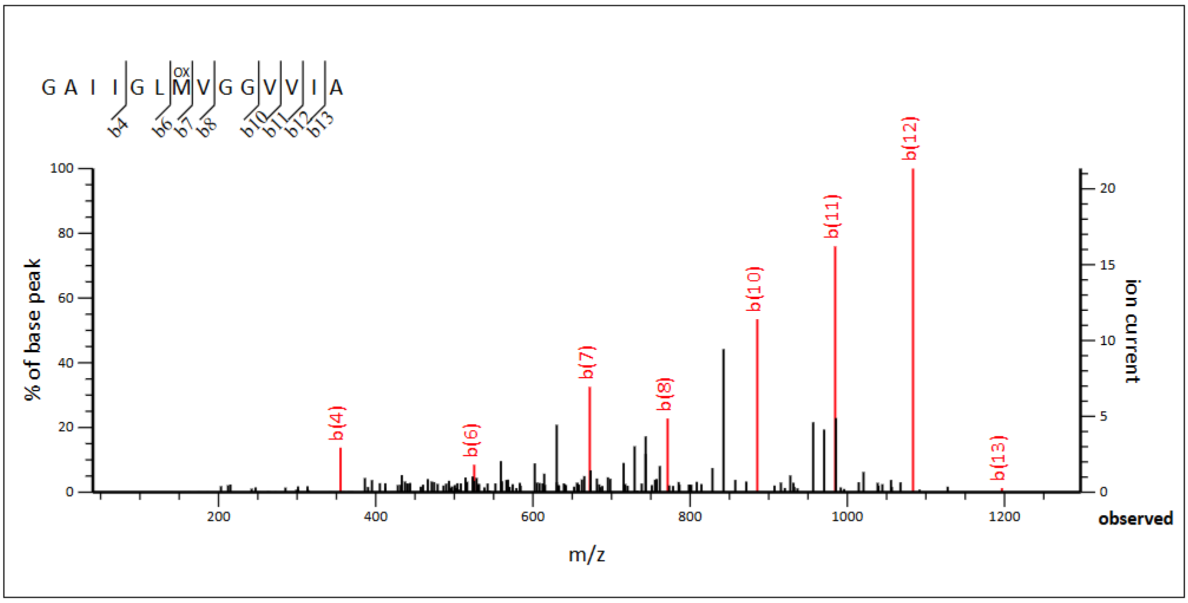


**Monoisotopic mass of neutral peptide Mr(calc):** 1284.7476
**Variable modifications:**
M7 : Oxidation (M), with neutral losses 0.0000(shown in table), 63.9983
**Ions Score:** 68 **Expect:** 1.7e-07
**Matches :** 8/78 fragment ions using 11 most intense peaks

| # | b | $b^{++}$ | Seq. | y | $y^{++}$ | # |
|---|---|---|---|---|---|---|
| 1 | 58.0287 | 29.5180 | G | | | 14 |
| 2 | 129.0659 | 65.0366 | A | 1228.7334 | 614.8703 | 13 |
| 3 | 242.1499 | 121.5786 | I | 1157.6962 | 579.3518 | 12 |
| 4 | ***355.2340*** | 178.1206 | I | 1044.6122 | 522.8097 | 11 |
| 5 | 412.2554 | 206.6314 | G | 931.5281 | 466.2677 | 10 |
| 6 | ***525.3395*** | 263.1734 | L | 874.5067 | 437.7570 | 9 |
| 7 | ***672.3749*** | 336.6911 | M | 761.4226 | 381.2149 | 8 |
| 8 | ***771.4433*** | 386.2253 | V | 614.3872 | 307.6972 | 7 |
| 9 | 828.4648 | 414.7360 | G | 515.3188 | 258.1630 | 6 |
| 10 | ***885.4863*** | 443.2468 | G | 458.2973 | 229.6523 | 5 |
| 11 | ***984.5547*** | 492.7810 | V | 401.2758 | 201.1416 | 4 |
| 12 | ***1083.6231*** | 542.3152 | V | 302.2074 | 151.6074 | 3 |
| 13 | ***1196.7071*** | 598.8572 | I | 203.1390 | 102.0731 | 2 |
| 14 | | | A | 90.0550 | 45.5311 | 1 |

### DELETION Aβ42 peptide detections

## In s1Aβ42

### H6

MS/MS Fragmentation of peptide **DSGYEVHHQK**

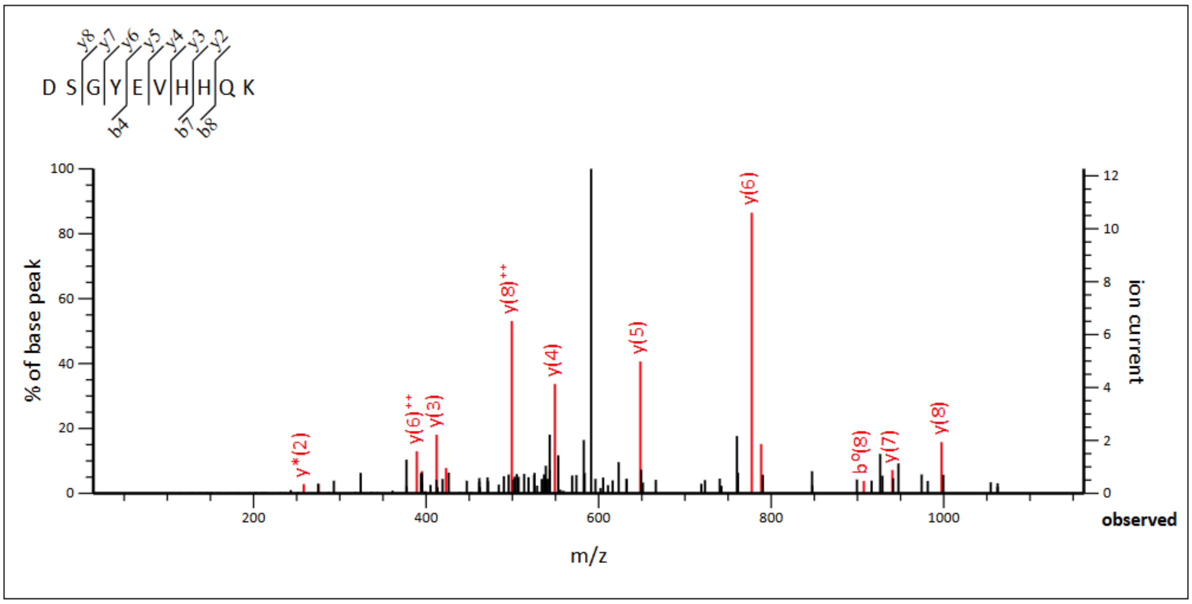


**Monoisotopic mass of neutral peptide Mr(calc):** 1198.5367
**Ions Score:** 35 **Expect:** 0.00034
**Matches :** 14/82 fragment ions using 39 most intense peaks

| # | b | $b^{++}$ | b* | $b^{*++}$ | $b^0$ | $b^{0++}$ | Seq. | y | $y^{++}$ | y* | $y^{*++}$ | $y^0$ | $y^{0++}$ | # |
|---|---|---|---|---|---|---|---|---|---|---|---|---|---|---|
| 1 | 116.0342 | 58.5207 | | | 98.0237 | 49.5155 | D | | | | | | | 10 |
| 2 | 203.0662 | 102.0368 | | | 185.0557 | 93.0315 | S | 1084.5170 | 542.7622 | 1067.4905 | 534.2489 | 1066.5065 | 533.7569 | 9 |
| 3 | 260.0877 | 130.5475 | | | 242.0771 | 121.5422 | G | ***997.4850*** | **499.2461** | 980.4585 | 490.7329 | 979.4744 | 490.2409 | 8 |
| 4 | **423.1510** | 212.0792 | | | 405.1405 | 203.0739 | Y | ***940.4635*** | 470.7354 | 923.4370 | 462.2221 | 922.4530 | 461.7301 | 7 |
| 5 | 552.1936 | 276.6005 | | | 534.1831 | 267.5952 | E | ***777.4002*** | **389.2037** | 760.3737 | 380.6905 | 759.3896 | 380.1985 | 6 |
| 6 | 651.2620 | 326.1347 | | | 633.2515 | 317.1294 | V | ***648.3576*** | 324.6824 | 631.3311 | 316.1692 | | | 5 |
| 7 | **788.3210** | 394.6641 | | | 770.3104 | 385.6588 | H | ***549.2892*** | 275.1482 | 532.2627 | 266.6350 | | | 4 |
| 8 | 925.3799 | 463.1936 | | | 907.3693 | 454.1883 | H | ***412.2303*** | 206.6188 | 395.2037 | 198.1055 | | | 3 |
| 9 | 1053.4384 | 527.2229 | 1036.4119 | 518.7096 | 1035.4279 | 518.2176 | Q | ***275.1714*** | 138.0893 | 258.1448 | 129.5761 | | | 2 |
| 10 | | | | | | | K | 147.1128 | 74.0600 | 130.0863 | 65.5468 | | | 1 |

## H13/14

MS/MS Fragmentation of peptide **HDSGYEVHQK**

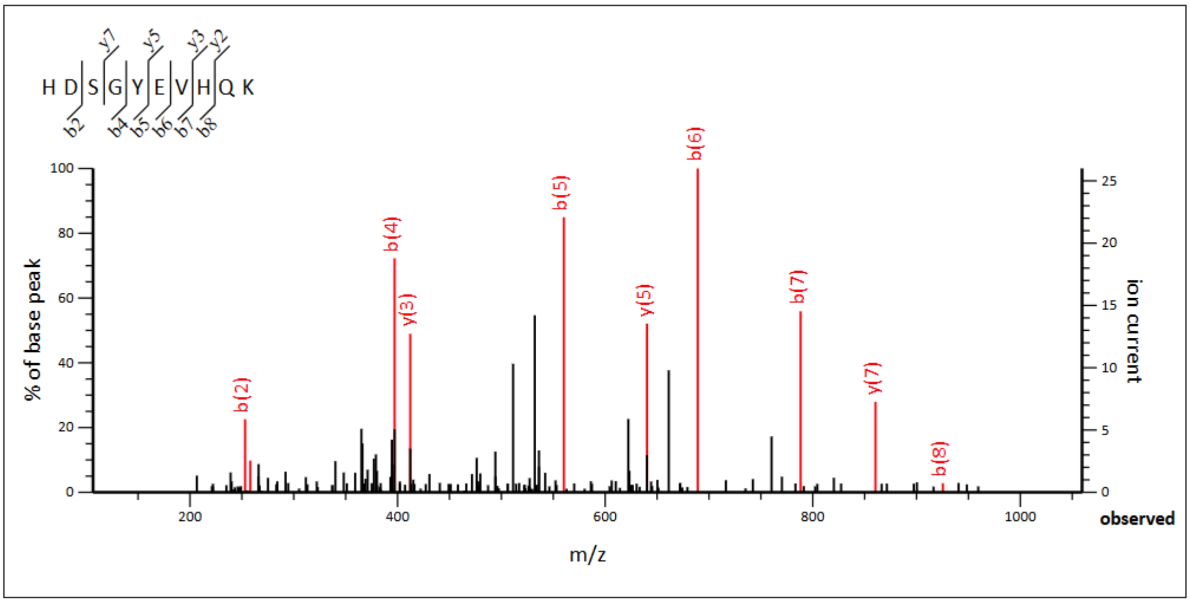


**Monoisotopic mass of neutral peptide Mr(calc):** 1198.5367
**Ions Score:** 45 **Expect:** 3.3e-05
**Matches :** 10/82 fragment ions using 16 most intense peaks

| # | b | $b^{++}$ | b* | $b^{*++}$ | $b^0$ | $b^{0++}$ | Seq. | y | $y^{++}$ | y* | $y^{*++}$ | $y^0$ | $y^{0++}$ | # |
|---|---|---|---|---|---|---|---|---|---|---|---|---|---|---|
| 1 | 138.0662 | 69.5367 | | | | | H | | | | | | | 10 |
| 2 | ***253.0931*** | 127.0502 | | | 235.0826 | 118.0449 | D | 1062.4851 | 531.7462 | 1045.4585 | 523.2329 | 1044.4745 | 522.7409 | 9 |
| 3 | 340.1252 | 170.5662 | | | 322.1146 | 161.5609 | S | 947.4581 | 474.2327 | 930.4316 | 465.7194 | 929.4476 | 465.2274 | 8 |
| 4 | ***397.1466*** | 199.0769 | | | 379.1361 | 190.0717 | G | **860.4261** | 430.7167 | 843.3995 | 422.2034 | 842.4155 | 421.7114 | 7 |
| 5 | ***560.2100*** | 280.6086 | | | 542.1994 | 271.6033 | Y | 803.4046 | 402.2060 | 786.3781 | 393.6927 | 785.3941 | 393.2007 | 6 |
| 6 | ***689.2525*** | 345.1299 | | | 671.2420 | 336.1246 | E | **640.3413** | 320.6743 | 623.3148 | 312.1610 | 622.3307 | 311.6690 | 5 |
| 7 | ***788.3210*** | 394.6641 | | | 770.3104 | 385.6588 | V | 511.2987 | 256.1530 | 494.2722 | 247.6397 | | | 4 |
| 8 | ***925.3799*** | 463.1936 | | | 907.3693 | 454.1883 | H | **412.2303** | 206.6188 | 395.2037 | 198.1055 | | | 3 |
| 9 | 1053.4384 | 527.2229 | 1036.4119 | 518.7096 | 1035.4279 | 518.2176 | Q | 275.1714 | 138.0893 | 258.1448 | 129.5761 | | | 2 |
| 10 | | | | | | | K | 147.1128 | 74.0600 | 130.0863 | 65.5468 | | | 1 |

## K16

MS/MS Fragmentation of peptide **HDSGYEVHHQLVFFAEDVGSNK**

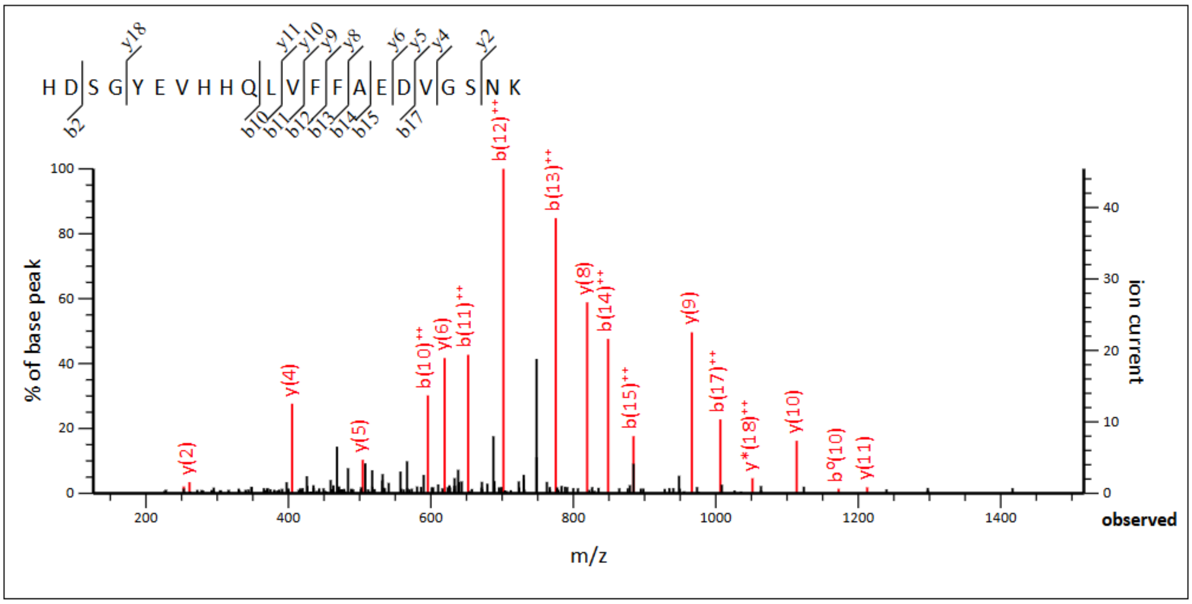


**Monoisotopic mass of neutral peptide Mr(calc):** 2514.1564
**Ions Score:** 65 **Expect:** 3.3e-07
**Matches :** 18/228 fragment ions using 23 most intense peaks

| # | b | $b^{++}$ | $b^{*}$ | $b^{*++}$ | $b^{0}$ | $b^{0++}$ | Seq. | y | $y^{++}$ | $y^{*}$ | $y^{*++}$ | $y^{0}$ | $y^{0++}$ | # |
|---|---|---|---|---|---|---|---|---|---|---|---|---|---|---|
| 1 | 138.0662 | 69.5367 | | | | | H | | | | | | | 22 |
| 2 | **253.0931** | 127.0502 | | | 235.0826 | 118.0449 | D | 2378.1048 | 1189.5560 | 2361.0782 | 1181.0427 | 2360.0942 | 1180.5507 | 21 |
| 3 | 340.1252 | 170.5662 | | | 322.1146 | 161.5609 | S | 2263.0778 | 1132.0425 | 2246.0513 | 1123.5293 | 2245.0673 | 1123.0373 | 20 |
| 4 | 397.1466 | 199.0769 | | | 379.1361 | 190.0717 | G | 2176.0458 | 1088.5265 | 2159.0192 | 1080.0133 | 2158.0352 | 1079.5213 | 19 |
| 5 | 560.2100 | 280.6086 | | | 542.1994 | 271.6033 | Y | 2119.0243 | 1060.0158 | 2101.9978 | 1051.5025 | 2101.0138 | 1051.0105 | 18 |
| 6 | 689.2525 | 345.1299 | | | 671.2420 | 336.1246 | E | 1955.9610 | 978.4841 | 1938.9345 | 969.9709 | 1937.9504 | 969.4789 | 17 |
| 7 | 788.3210 | 394.6641 | | | 770.3104 | 385.6588 | V | 1826.9184 | 913.9628 | 1809.8919 | 905.4496 | 1808.9078 | 904.9576 | 16 |
| 8 | 925.3799 | 463.1936 | | | 907.3693 | 454.1883 | H | 1727.8500 | 864.4286 | 1710.8234 | 855.9154 | 1709.8394 | 855.4234 | 15 |
| 9 | 1062.4388 | 531.7230 | | | 1044.4282 | 522.7177 | H | 1590.7911 | 795.8992 | 1573.7645 | 787.3859 | 1572.7805 | 786.8939 | 14 |
| 10 | 1190.4974 | ***595.7523*** | 1173.4708 | 587.2390 | 1172.4868 | 586.7470 | Q | 1453.7322 | 727.3697 | 1436.7056 | 718.8564 | 1435.7216 | 718.3644 | 13 |
| 11 | 1303.5814 | ***652.2944*** | 1286.5549 | 643.7811 | 1285.5709 | 643.2891 | L | 1325.6736 | 663.3404 | 1308.6470 | 654.8272 | 1307.6630 | 654.3352 | 12 |
| 12 | 1402.6498 | ***701.8286*** | 1385.6233 | 693.3153 | 1384.6393 | 692.8233 | V | ***1212.5895*** | 606.7984 | 1195.5630 | 598.2851 | 1194.5790 | 597.7931 | 11 |
| 13 | 1549.7183 | ***775.3628*** | 1532.6917 | 766.8495 | 1531.7077 | 766.3575 | F | ***1113.5211*** | 557.2642 | 1096.4946 | 548.7509 | 1095.5106 | 548.2589 | 10 |
| 14 | 1696.7867 | ***848.8970*** | 1679.7601 | 840.3837 | 1678.7761 | 839.8917 | F | ***966.4527*** | 483.7300 | 949.4262 | 475.2167 | 948.4421 | 474.7247 | 9 |
| 15 | 1767.8238 | ***884.4155*** | 1750.7972 | 875.9023 | 1749.8132 | 875.4102 | A | ***819.3843*** | 410.1958 | 802.3577 | 401.6825 | 801.3737 | 401.1905 | 8 |
| 16 | 1896.8664 | 948.9368 | 1879.8398 | 940.4236 | 1878.8558 | 939.9315 | E | 748.3472 | 374.6772 | 731.3206 | 366.1639 | 730.3366 | 365.6719 | 7 |
| 17 | 2011.8933 | ***1006.4503*** | 1994.8668 | 997.9370 | 1993.8828 | 997.4450 | D | ***619.3046*** | 310.1559 | 602.2780 | 301.6427 | 601.2940 | 301.1506 | 6 |
| 18 | 2110.9617 | 1055.9845 | 2093.9352 | 1047.4712 | 2092.9512 | 1046.9792 | V | ***504.2776*** | 252.6425 | 487.2511 | 244.1292 | 486.2671 | 243.6372 | 5 |
| 19 | 2167.9832 | 1084.4952 | 2150.9566 | 1075.9820 | 2149.9726 | 1075.4900 | G | ***405.2092*** | 203.1082 | 388.1827 | 194.5950 | 387.1987 | 194.1030 | 4 |
| 20 | 2255.0152 | 1128.0112 | 2237.9887 | 1119.4980 | 2237.0047 | 1119.0060 | S | 348.1878 | 174.5975 | 331.1612 | 166.0842 | 330.1772 | 165.5922 | 3 |
| 21 | 2369.0582 | 1185.0327 | 2352.0316 | 1176.5194 | 2351.0476 | 1176.0274 | N | ***261.1557*** | 131.0815 | 244.1292 | 122.5682 | | | 2 |
| 22 | | | | | | | K | 147.1128 | 74.0600 | 130.0863 | 65.5468 | | | 1 |

## L17

MS/MS Fragmentation of peptide **VFFAEDVGSNK**

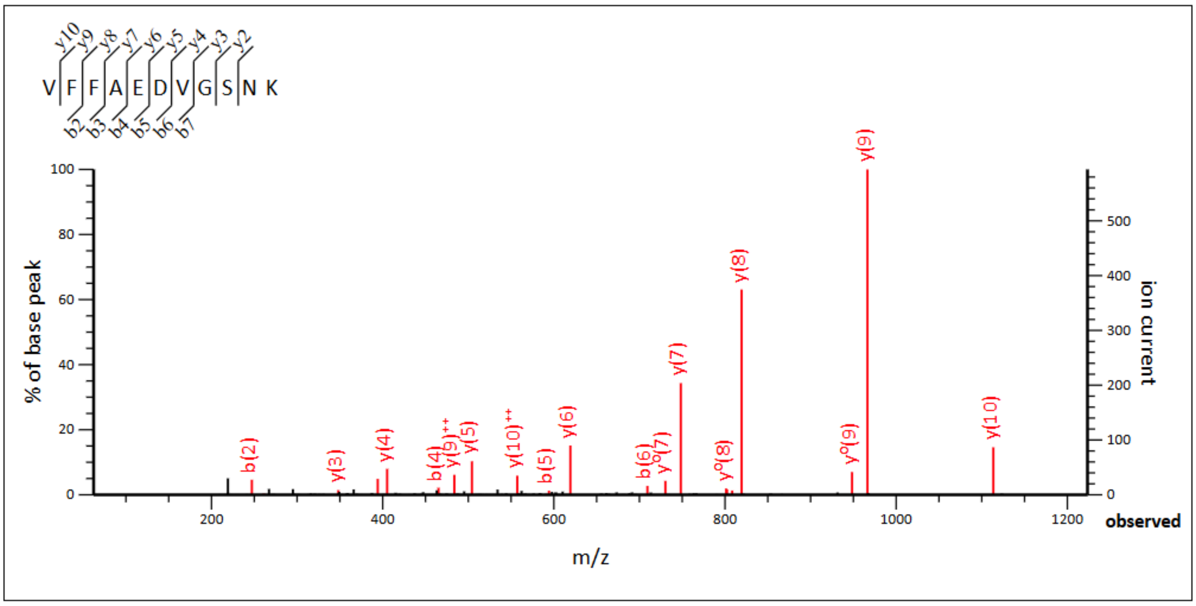


**Monoisotopic mass of neutral peptide Mr(calc):** 1211.5823
**Ions Score:** 69 **Expect:** 1.3e-07
**Matches :** 21/90 fragment ions using 36 most intense peaks

| # | b | $b^{++}$ | b* | $b^{*++}$ | $b^0$ | $b^{0++}$ | Seq. | y | $y^{++}$ | y* | $y^{*++}$ | $y^0$ | $y^{0++}$ | # |
|---|---|---|---|---|---|---|---|---|---|---|---|---|---|---|
| 1 | 100.0757 | 50.5415 | | | | | V | | | | | | | 11 |
| 2 | **247.1441** | 124.0757 | | | | | F | ***1113.5211*** | **557.2642** | 1096.4946 | 548.7509 | 1095.5106 | 548.2589 | 10 |
| 3 | **394.2125** | 197.6099 | | | | | F | ***966.4527*** | **483.7300** | 949.4262 | 475.2167 | 948.4421 | 474.7247 | 9 |
| 4 | **465.2496** | 233.1285 | | | | | A | ***819.3843*** | 410.1958 | 802.3577 | 401.6825 | 801.3737 | 401.1905 | 8 |
| 5 | **594.2922** | 297.6498 | | | 576.2817 | 288.6445 | E | ***748.3472*** | 374.6772 | 731.3206 | 366.1639 | 730.3366 | 365.6719 | 7 |
| 6 | **709.3192** | 355.1632 | | | 691.3086 | 346.1579 | D | ***619.3046*** | 310.1559 | 602.2780 | 301.6427 | 601.2940 | 301.1506 | 6 |
| 7 | **808.3876** | 404.6974 | | | 790.3770 | 395.6921 | V | ***504.2776*** | 252.6425 | 487.2511 | 244.1292 | 486.2671 | 243.6372 | 5 |
| 8 | 865.4090 | 433.2082 | | | 847.3985 | 424.2029 | G | ***405.2092*** | 203.1082 | 388.1827 | 194.5950 | 387.1987 | 194.1030 | 4 |
| 9 | 952.4411 | 476.7242 | | | 934.4305 | 467.7189 | S | ***348.1878*** | 174.5975 | 331.1612 | 166.0842 | 330.1772 | 165.5922 | 3 |
| 10 | 1066.4840 | 533.7456 | 1049.4575 | 525.2324 | 1048.4734 | 524.7404 | N | ***261.1557*** | 131.0815 | 244.1292 | 122.5682 | | | 2 |
| 11 | | | | | | | K | 147.1128 | 74.0600 | 130.0863 | 65.5468 | | | 1 |

## V18

MS/MS Fragmentation of peptide **LFFAEDVGSNK**

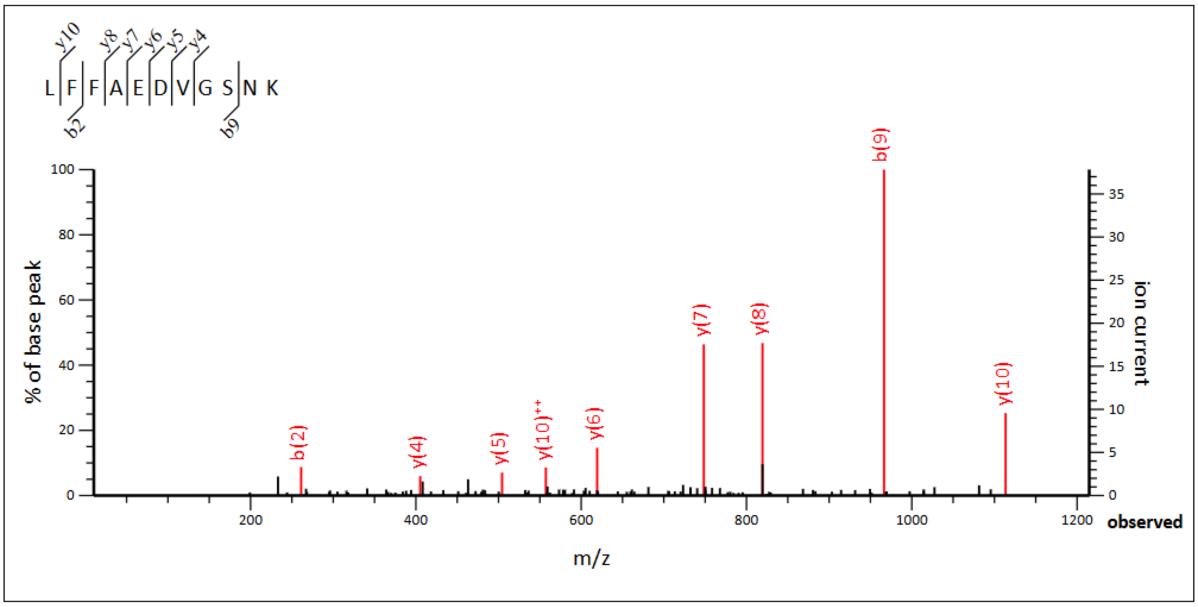


**Monoisotopic mass of neutral peptide Mr(calc):** 1225.5979
**Ions Score:** 78 **Expect:** 1.7e-08
**Matches :** 11/90 fragment ions using 11 most intense peaks

| # | b | $b^{++}$ | b* | $b^{*++}$ | $b^{0}$ | $b^{0++}$ | Seq. | y | $y^{++}$ | y* | $y^{*++}$ | $y^{0}$ | $y^{0++}$ | # |
|---|---|---|---|---|---|---|---|---|---|---|---|---|---|---|
| 1 | 114.0913 | 57.5493 | | | | | L | | | | | | | 11 |
| 2 | **261.1598** | 131.0835 | | | | | F | ***1113.5211*** | **557.2642** | 1096.4946 | 548.7509 | 1095.5106 | 548.2589 | 10 |
| 3 | 408.2282 | 204.6177 | | | | | F | ***966.4527*** | 483.7300 | 949.4262 | 475.2167 | 948.4421 | 474.7247 | 9 |
| 4 | 479.2653 | 240.1363 | | | | | A | ***819.3843*** | 410.1958 | 802.3577 | 401.6825 | 801.3737 | 401.1905 | 8 |
| 5 | 608.3079 | 304.6576 | | | 590.2973 | 295.6523 | E | ***748.3472*** | 374.6772 | 731.3206 | 366.1639 | 730.3366 | 365.6719 | 7 |
| 6 | 723.3348 | 362.1710 | | | 705.3243 | 353.1658 | D | ***619.3046*** | 310.1559 | 602.2780 | 301.6427 | 601.2940 | 301.1506 | 6 |
| 7 | 822.4032 | 411.7053 | | | 804.3927 | 402.7000 | V | ***504.2776*** | 252.6425 | 487.2511 | 244.1292 | 486.2671 | 243.6372 | 5 |
| 8 | 879.4247 | 440.2160 | | | 861.4141 | 431.2107 | G | ***405.2092*** | 203.1082 | 388.1827 | 194.5950 | 387.1987 | 194.1030 | 4 |
| 9 | **966.4567** | 483.7320 | | | 948.4462 | 474.7267 | S | 348.1878 | 174.5975 | 331.1612 | 166.0842 | 330.1772 | 165.5922 | 3 |
| 10 | 1080.4997 | 540.7535 | 1063.4731 | 532.2402 | 1062.4891 | 531.7482 | N | ***261.1557*** | 131.0815 | 244.1292 | 122.5682 | | | 2 |
| 11 | | | | | | | K | 147.1128 | 74.0600 | 130.0863 | 65.5468 | | | 1 |

## F19/20

MS/MS Fragmentation of peptide **LVFAEDVGSNK**

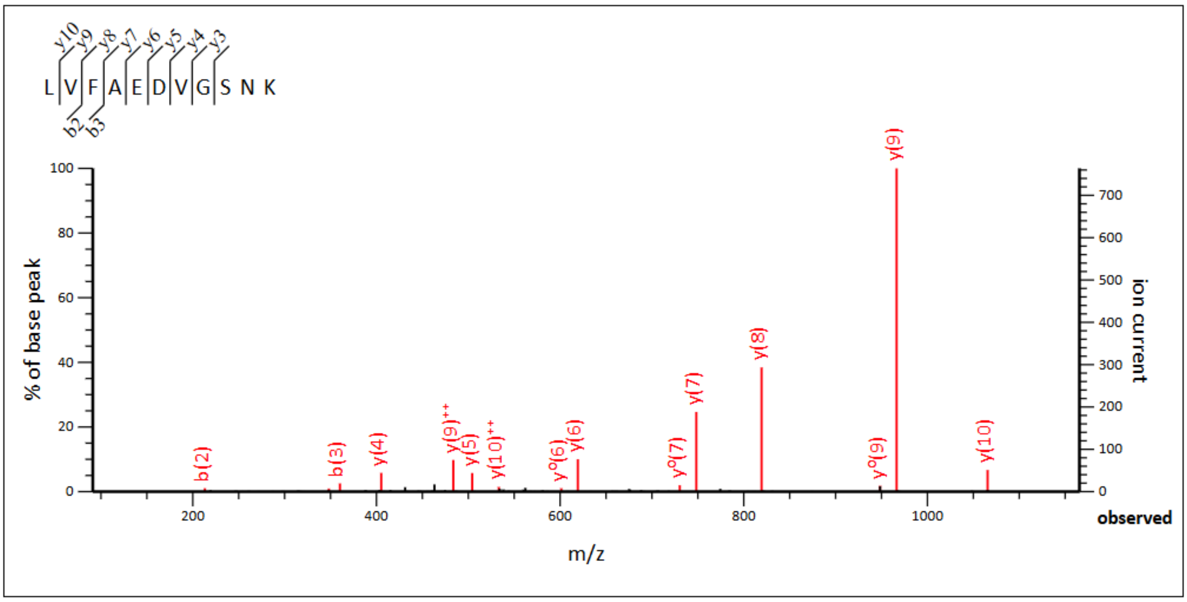


**Monoisotopic mass of neutral peptide Mr(calc):** 1177.5979
**Ions Score:** 79 **Expect:** 1.3e-08
**Matches :** 15/90 fragment ions using 18 most intense peaks

| # | b | $b^{++}$ | b* | $b^{*++}$ | $b^{0}$ | $b^{0++}$ | Seq. | y | $y^{++}$ | y* | $y^{*++}$ | $y^{0}$ | $y^{0++}$ | # |
|---|---|---|---|---|---|---|---|---|---|---|---|---|---|---|
| 1 | 114.0913 | 57.5493 | | | | | L | | | | | | | 11 |
| 2 | **213.1598** | 107.0835 | | | | | V | ***1065.5211*** | **533.2642** | 1048.4946 | 524.7509 | 1047.5106 | 524.2589 | 10 |
| 3 | **360.2282** | 180.6177 | | | | | F | ***966.4527*** | **483.7300** | 949.4262 | 475.2167 | 948.4421 | 474.7247 | 9 |
| 4 | 431.2653 | 216.1363 | | | | | A | ***819.3843*** | 410.1958 | 802.3577 | 401.6825 | 801.3737 | 401.1905 | 8 |
| 5 | 560.3079 | 280.6576 | | | 542.2973 | 271.6523 | E | ***748.3472*** | 374.6772 | 731.3206 | 366.1639 | 730.3366 | 365.6719 | 7 |
| 6 | 675.3348 | 338.1710 | | | 657.3243 | 329.1658 | D | ***619.3046*** | 310.1559 | 602.2780 | 301.6427 | 601.2940 | 301.1506 | 6 |
| 7 | 774.4032 | 387.7053 | | | 756.3927 | 378.7000 | V | ***504.2776*** | 252.6425 | 487.2511 | 244.1292 | 486.2671 | 243.6372 | 5 |
| 8 | 831.4247 | 416.2160 | | | 813.4141 | 407.2107 | G | ***405.2092*** | 203.1082 | 388.1827 | 194.5950 | 387.1987 | 194.1030 | 4 |
| 9 | 918.4567 | 459.7320 | | | 900.4462 | 450.7267 | S | ***348.1878*** | 174.5975 | 331.1612 | 166.0842 | 330.1772 | 165.5922 | 3 |
| 10 | 1032.4997 | 516.7535 | 1015.4731 | 508.2402 | 1014.4891 | 507.7482 | N | 261.1557 | 131.0815 | 244.1292 | 122.5682 | | | 2 |
| 11 | | | | | | | K | 147.1128 | 74.0600 | 130.0863 | 65.5468 | | | 1 |

## A21

MS/MS Fragmentation of peptide **LVFFEDVGSNK**

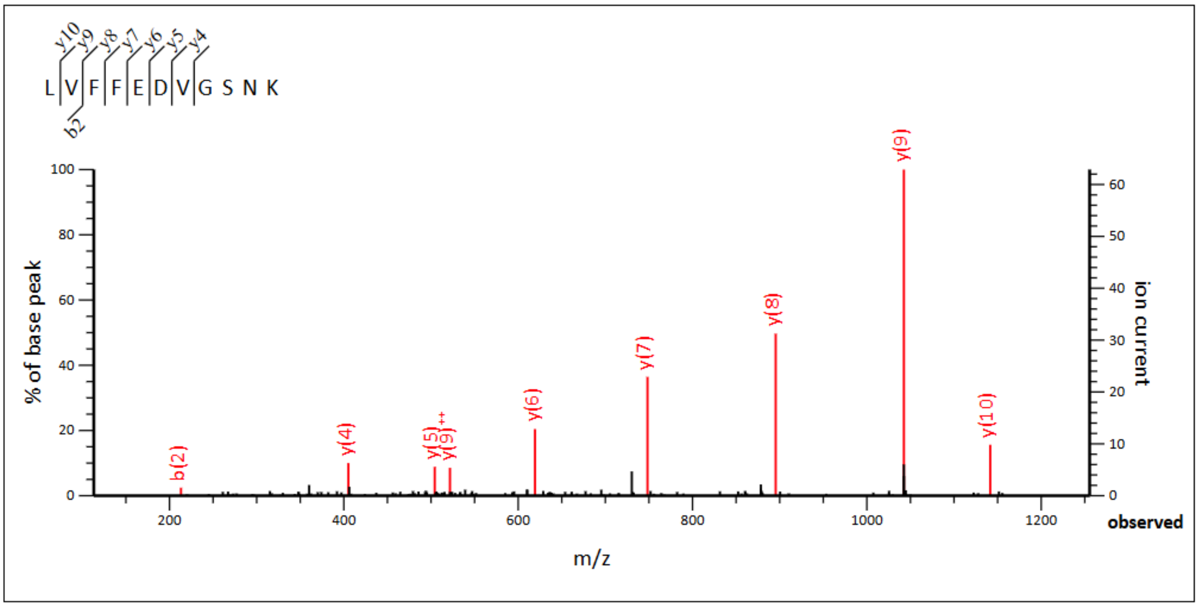


**Monoisotopic mass of neutral peptide Mr(calc):** 1253.6292
**Ions Score:** 70 **Expect:** 1.1e-07
**Matches :** 9/90 fragment ions using 10 most intense peaks

| # | b | $b^{++}$ | b* | $b^{*++}$ | $b^{0}$ | $b^{0++}$ | Seq. | y | $y^{++}$ | y* | $y^{*++}$ | $y^{0}$ | $y^{0++}$ | # |
|---|---|---|---|---|---|---|---|---|---|---|---|---|---|---|
| 1 | 114.0913 | 57.5493 | | | | | L | | | | | | | 11 |
| 2 | 213.1598 | 107.0835 | | | | | V | 1141.5524 | 571.2798 | 1124.5259 | 562.7666 | 1123.5419 | 562.2746 | 10 |
| 3 | 360.2282 | 180.6177 | | | | | F | 1042.4840 | 521.7456 | 1025.4575 | 513.2324 | 1024.4734 | 512.7404 | 9 |
| 4 | 507.2966 | 254.1519 | | | | | F | 895.4156 | 448.2114 | 878.3890 | 439.6982 | 877.4050 | 439.2061 | 8 |
| 5 | 636.3392 | 318.6732 | | | 618.3286 | 309.6679 | E | 748.3472 | 374.6772 | 731.3206 | 366.1639 | 730.3366 | 365.6719 | 7 |
| 6 | 751.3661 | 376.1867 | | | 733.3556 | 367.1814 | D | 619.3046 | 310.1559 | 602.2780 | 301.6427 | 601.2940 | 301.1506 | 6 |
| 7 | 850.4345 | 425.7209 | | | 832.4240 | 416.7156 | V | 504.2776 | 252.6425 | 487.2511 | 244.1292 | 486.2671 | 243.6372 | 5 |
| 8 | 907.4560 | 454.2316 | | | 889.4454 | 445.2264 | G | 405.2092 | 203.1082 | 388.1827 | 194.5950 | 387.1987 | 194.1030 | 4 |
| 9 | 994.4880 | 497.7477 | | | 976.4775 | 488.7424 | S | 348.1878 | 174.5975 | 331.1612 | 166.0842 | 330.1772 | 165.5922 | 3 |
| 10 | 1108.5310 | 554.7691 | 1091.5044 | 546.2558 | 1090.5204 | 545.7638 | N | 261.1557 | 131.0815 | 244.1292 | 122.5682 | | | 2 |
| 11 | | | | | | | K | 147.1128 | 74.0600 | 130.0863 | 65.5468 | | | 1 |

## G25

MS/MS Fragmentation of peptide **LVFFAEDVSNK**

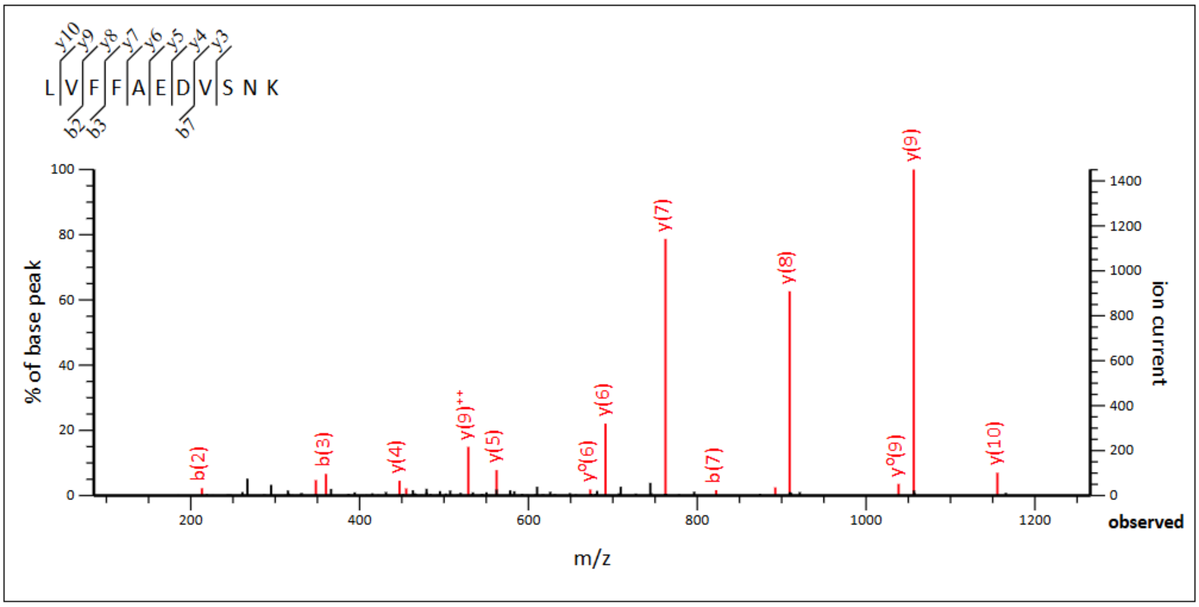


**Monoisotopic mass of neutral peptide Mr(calc):** 1267.6449
**Ions Score:** 73 **Expect:** 4.5e-08
**Matches :** 16/88 fragment ions using 20 most intense peaks

| # | b | $b^{++}$ | $b^*$ | $b^{*++}$ | $b^0$ | $b^{0++}$ | Seq. | y | $y^{++}$ | $y^*$ | $y^{*++}$ | $y^0$ | $y^{0++}$ | # |
|---|---|---|---|---|---|---|---|---|---|---|---|---|---|---|
| 1 | 114.0913 | 57.5493 | | | | | L | | | | | | | 11 |
| 2 | **213.1598** | 107.0835 | | | | | V | ***1155.5681*** | 578.2877 | 1138.5415 | 569.7744 | 1137.5575 | 569.2824 | 10 |
| 3 | **360.2282** | 180.6177 | | | | | F | ***1056.4997*** | **528.7535** | 1039.4731 | 520.2402 | 1038.4891 | 519.7482 | 9 |
| 4 | 507.2966 | 254.1519 | | | | | F | ***909.4312*** | **455.2193** | 892.4047 | 446.7060 | 891.4207 | 446.2140 | 8 |
| 5 | 578.3337 | 289.6705 | | | | | A | ***762.3628*** | 381.6850 | 745.3363 | 373.1718 | 744.3523 | 372.6798 | 7 |
| 6 | 707.3763 | 354.1918 | | | 689.3657 | 345.1865 | E | ***691.3257*** | 346.1665 | 674.2992 | 337.6532 | 673.3151 | 337.1612 | 6 |
| 7 | **822.4032** | 411.7053 | | | 804.3927 | 402.7000 | D | ***562.2831*** | 281.6452 | 545.2566 | 273.1319 | 544.2726 | 272.6399 | 5 |
| 8 | 921.4716 | 461.2395 | | | 903.4611 | 452.2342 | V | ***447.2562*** | 224.1317 | 430.2296 | 215.6184 | 429.2456 | 215.1264 | 4 |
| 9 | 1008.5037 | 504.7555 | | | 990.4931 | 495.7502 | S | ***348.1878*** | 174.5975 | 331.1612 | 166.0842 | 330.1772 | 165.5922 | 3 |
| 10 | 1122.5466 | 561.7769 | 1105.5201 | 553.2637 | 1104.5360 | 552.7717 | N | 261.1557 | 131.0815 | 244.1292 | 122.5682 | | | 2 |
| 11 | | | | | | | K | 147.1128 | 74.0600 | 130.0863 | 65.5468 | | | 1 |

# S26

MS/MS Fragmentation of peptide **LVFFAEDVGNK**

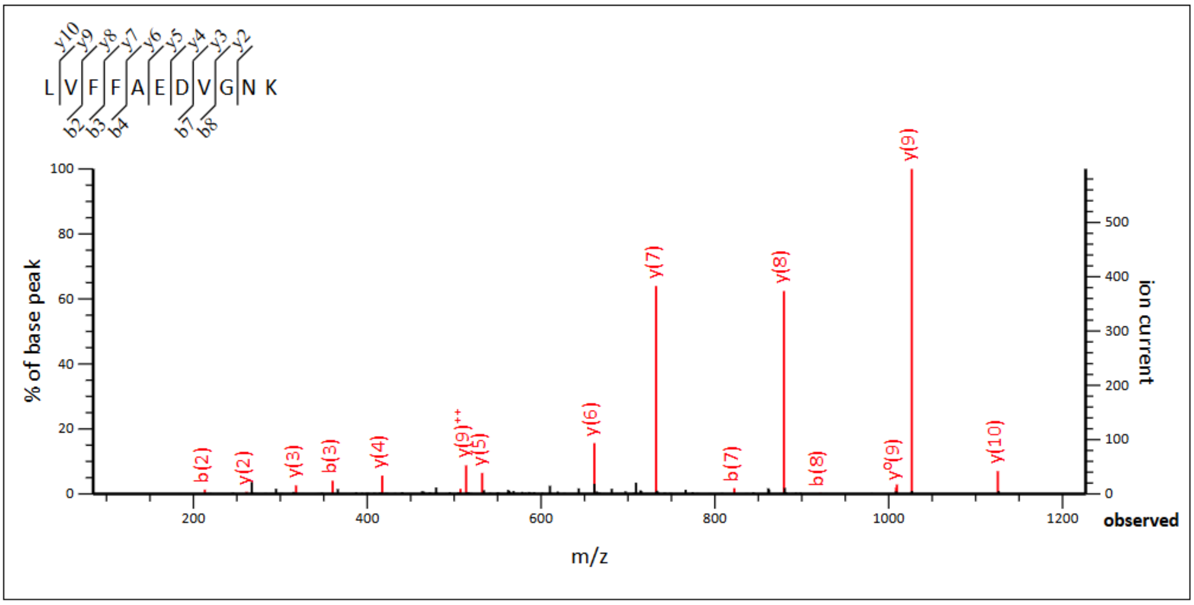


**Monoisotopic mass of neutral peptide Mr(calc):** 1237.6343
**Ions Score:** 71 **Expect:** 8e-08
**Matches :** 17/84 fragment ions using 29 most intense peaks

| # | b | $b^{++}$ | b* | $b^{*++}$ | $b^0$ | $b^{0++}$ | Seq. | y | $y^{++}$ | y* | $y^{*++}$ | $y^0$ | $y^{0++}$ | # |
|---|---|---|---|---|---|---|---|---|---|---|---|---|---|---|
| 1 | 114.0913 | 57.5493 | | | | | L | | | | | | | 11 |
| 2 | **213.1598** | 107.0835 | | | | | V | ***1125.5575*** | 563.2824 | 1108.5310 | 554.7691 | 1107.5469 | 554.2771 | 10 |
| 3 | **360.2282** | 180.6177 | | | | | F | ***1026.4891*** | **513.7482** | 1009.4625 | 505.2349 | 1008.4785 | 504.7429 | 9 |
| 4 | **507.2966** | 254.1519 | | | | | F | ***879.4207*** | 440.2140 | 862.3941 | 431.7007 | 861.4101 | 431.2087 | 8 |
| 5 | 578.3337 | 289.6705 | | | | | A | ***732.3523*** | 366.6798 | 715.3257 | 358.1665 | 714.3417 | 357.6745 | 7 |
| 6 | 707.3763 | 354.1918 | | | 689.3657 | 345.1865 | E | ***661.3151*** | 331.1612 | 644.2886 | 322.6479 | 643.3046 | 322.1559 | 6 |
| 7 | **822.4032** | 411.7053 | | | 804.3927 | 402.7000 | D | ***532.2726*** | 266.6399 | 515.2460 | 258.1266 | 514.2620 | 257.6346 | 5 |
| 8 | **921.4716** | 461.2395 | | | 903.4611 | 452.2342 | V | ***417.2456*** | 209.1264 | 400.2191 | 200.6132 | | | 4 |
| 9 | 978.4931 | 489.7502 | | | 960.4825 | 480.7449 | G | ***318.1772*** | 159.5922 | 301.1506 | 151.0790 | | | 3 |
| 10 | 1092.5360 | 546.7717 | 1075.5095 | 538.2584 | 1074.5255 | 537.7664 | N | ***261.1557*** | 131.0815 | 244.1292 | 122.5682 | | | 2 |
| 11 | | | | | | | K | 147.1128 | 74.0600 | 130.0863 | 65.5468 | | | 1 |

## N27

MS/MS Fragmentation of peptide **LVFFAEDVGSK**

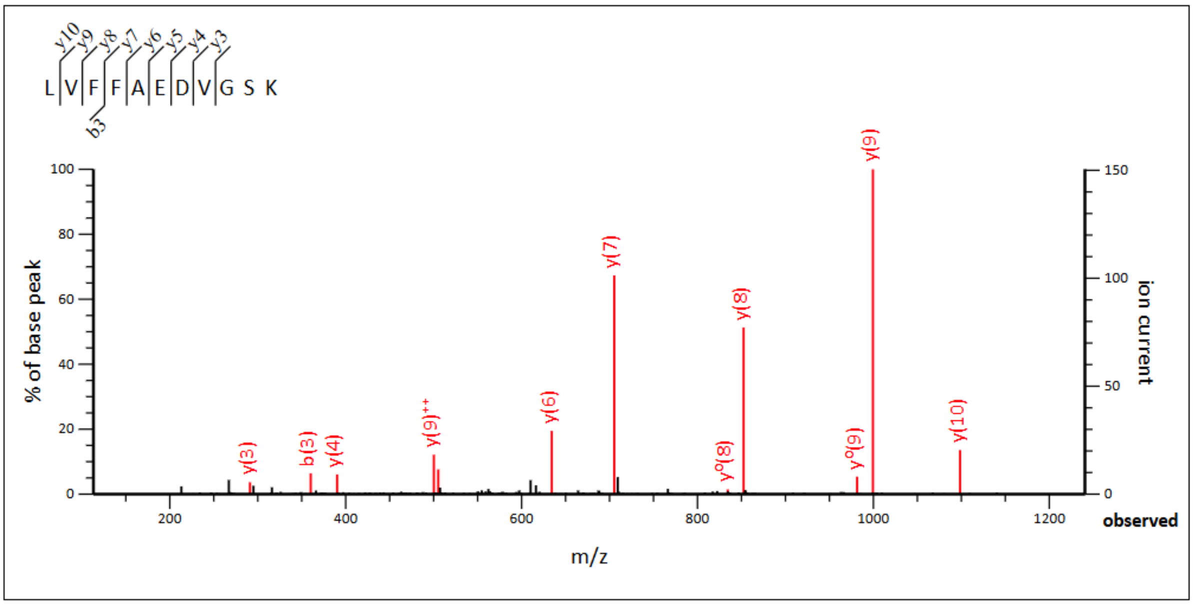


**Monoisotopic mass of neutral peptide Mr(calc):** 1210.6234
**Ions Score:** 66 **Expect:** 2.3e-07
**Matches :** 12/88 fragment ions using 19 most intense peaks

| # | b | b$^{++}$ | b$^{0}$ | b$^{0++}$ | Seq. | y | y$^{++}$ | y* | y*$^{++}$ | y$^{0}$ | y$^{0++}$ | # |
|---|---|---|---|---|---|---|---|---|---|---|---|---|
| 1 | 114.0913 | 57.5493 | | | L | | | | | | | 11 |
| 2 | 213.1598 | 107.0835 | | | V | ***1098.5466*** | 549.7769 | 1081.5201 | 541.2637 | 1080.5360 | 540.7717 | 10 |
| 3 | **360.2282** | 180.6177 | | | F | ***999.4782*** | **500.2427** | 982.4516 | 491.7295 | 981.4676 | 491.2374 | 9 |
| 4 | 507.2966 | 254.1519 | | | F | ***852.4098*** | 426.7085 | 835.3832 | 418.1953 | 834.3992 | 417.7032 | 8 |
| 5 | 578.3337 | 289.6705 | | | A | ***705.3414*** | 353.1743 | 688.3148 | 344.6610 | 687.3308 | 344.1690 | 7 |
| 6 | 707.3763 | 354.1918 | 689.3657 | 345.1865 | E | ***634.3042*** | 317.6558 | 617.2777 | 309.1425 | 616.2937 | 308.6505 | 6 |
| 7 | 822.4032 | 411.7053 | 804.3927 | 402.7000 | D | ***505.2617*** | 253.1345 | 488.2351 | 244.6212 | 487.2511 | 244.1292 | 5 |
| 8 | 921.4716 | 461.2395 | 903.4611 | 452.2342 | V | ***390.2347*** | 195.6210 | 373.2082 | 187.1077 | 372.2241 | 186.6157 | 4 |
| 9 | 978.4931 | 489.7502 | 960.4825 | 480.7449 | G | ***291.1663*** | 146.0868 | 274.1397 | 137.5735 | 273.1557 | 137.0815 | 3 |
| 10 | 1065.5251 | 533.2662 | 1047.5146 | 524.2609 | S | 234.1448 | 117.5761 | 217.1183 | 109.0628 | 216.1343 | 108.5708 | 2 |
| 11 | | | | | K | 147.1128 | 74.0600 | 130.0863 | 65.5468 | | | 1 |

#### In rAβ42

### H6

No identifications in rAβ42

### H13/14

No identifications in rAβ42

### K16

No identifications in rAβ42

## L17

MS/MS Fragmentation of peptide **VFFAEDVGSNK**

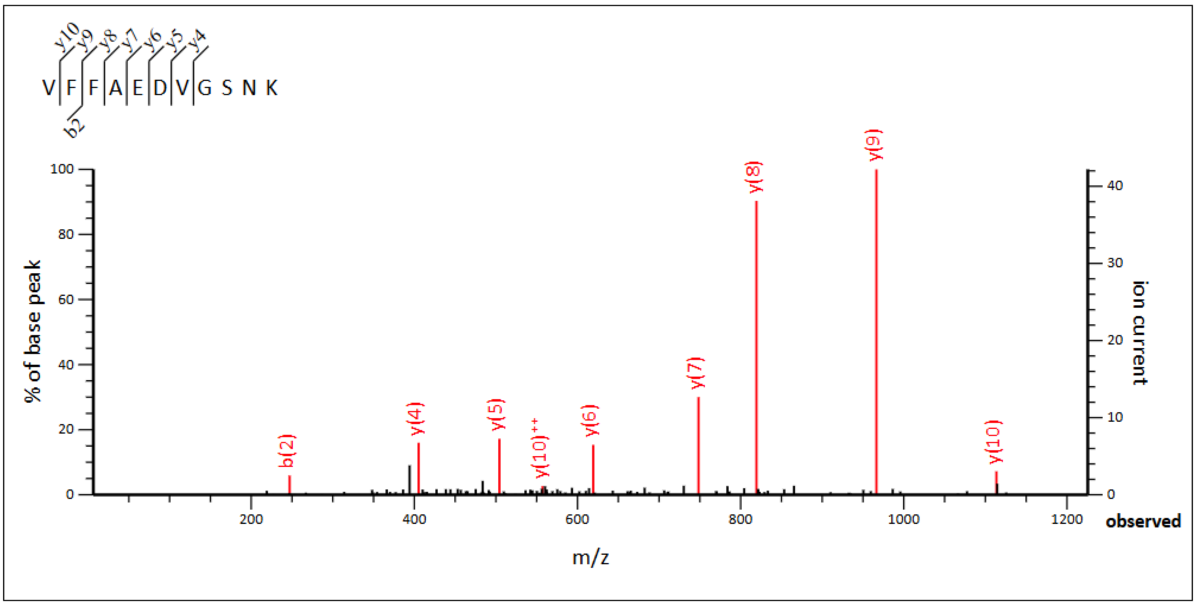


**Monoisotopic mass of neutral peptide Mr(calc):** 1211.5823
**Ions Score:** 67 **Expect:** 2e-07
**Matches :** 9/90 fragment ions using 11 most intense peaks

| # | b | $b^{++}$ | b* | $b^{*++}$ | $b^{0}$ | $b^{0++}$ | Seq. | y | $y^{++}$ | y* | $y^{*++}$ | $y^{0}$ | $y^{0++}$ | # |
|---|---|---|---|---|---|---|---|---|---|---|---|---|---|---|
| 1 | 100.0757 | 50.5415 | | | | | V | | | | | | | 11 |
| 2 | **247.1441** | 124.0757 | | | | | F | ***1113.5211*** | **557.2642** | 1096.4946 | 548.7509 | 1095.5106 | 548.2589 | 10 |
| 3 | 394.2125 | 197.6099 | | | | | F | ***966.4527*** | 483.7300 | 949.4262 | 475.2167 | 948.4421 | 474.7247 | 9 |
| 4 | 465.2496 | 233.1285 | | | | | A | ***819.3843*** | 410.1958 | 802.3577 | 401.6825 | 801.3737 | 401.1905 | 8 |
| 5 | 594.2922 | 297.6498 | | | 576.2817 | 288.6445 | E | ***748.3472*** | 374.6772 | 731.3206 | 366.1639 | 730.3366 | 365.6719 | 7 |
| 6 | 709.3192 | 355.1632 | | | 691.3086 | 346.1579 | D | ***619.3046*** | 310.1559 | 602.2780 | 301.6427 | 601.2940 | 301.1506 | 6 |
| 7 | 808.3876 | 404.6974 | | | 790.3770 | 395.6921 | V | ***504.2776*** | 252.6425 | 487.2511 | 244.1292 | 486.2671 | 243.6372 | 5 |
| 8 | 865.4090 | 433.2082 | | | 847.3985 | 424.2029 | G | ***405.2092*** | 203.1082 | 388.1827 | 194.5950 | 387.1987 | 194.1030 | 4 |
| 9 | 952.4411 | 476.7242 | | | 934.4305 | 467.7189 | S | 348.1878 | 174.5975 | 331.1612 | 166.0842 | 330.1772 | 165.5922 | 3 |
| 10 | 1066.4840 | 533.7456 | 1049.4575 | 525.2324 | 1048.4734 | 524.7404 | N | 261.1557 | 131.0815 | 244.1292 | 122.5682 | | | 2 |
| 11 | | | | | | | K | 147.1128 | 74.0600 | 130.0863 | 65.5468 | | | 1 |

# V18

MS/MS Fragmentation of peptide **LFFAEDVGSNK**

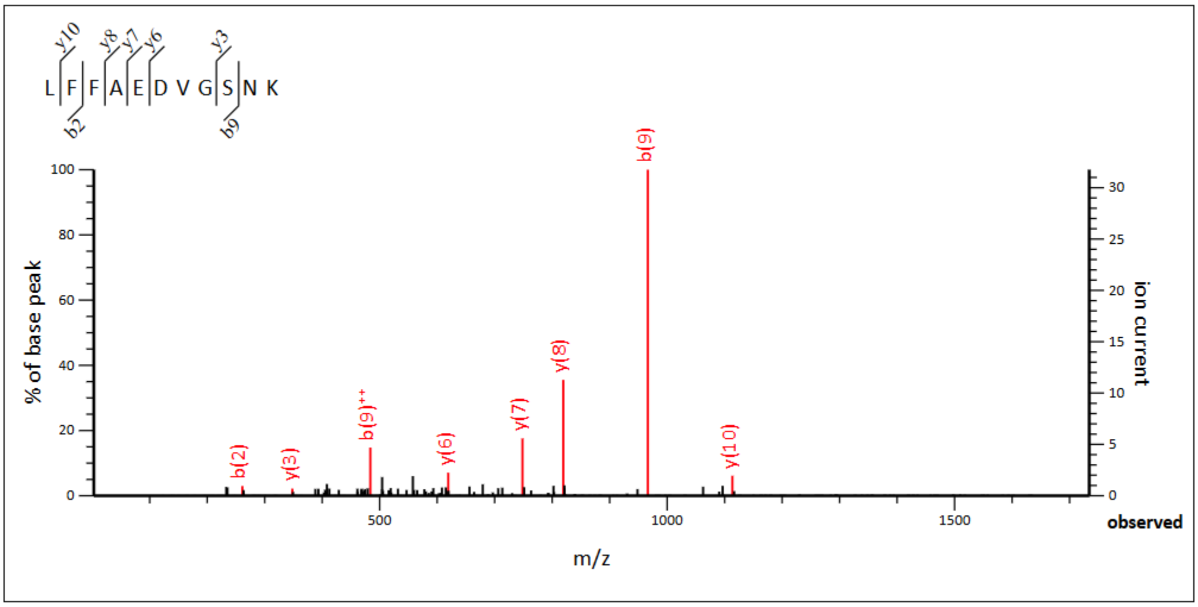


**Monoisotopic mass of neutral peptide Mr(calc):** 1225.5979
**Ions Score:** 57 **Expect:** 2e-06
**Matches :** 11/90 fragment ions using 12 most intense peaks

| # | b | $b^{++}$ | b* | $b^{*++}$ | $b^{0}$ | $b^{0++}$ | Seq. | y | $y^{++}$ | y* | $y^{*++}$ | $y^{0}$ | $y^{0++}$ | # |
|---|---|---|---|---|---|---|---|---|---|---|---|---|---|---|
| 1 | 114.0913 | 57.5493 | | | | | L | | | | | | | 11 |
| 2 | **261.1598** | 131.0835 | | | | | F | ***1113.5211*** | 557.2642 | 1096.4946 | 548.7509 | 1095.5106 | 548.2589 | 10 |
| 3 | 408.2282 | 204.6177 | | | | | F | ***966.4527*** | **483.7300** | 949.4262 | 475.2167 | 948.4421 | 474.7247 | 9 |
| 4 | 479.2653 | 240.1363 | | | | | A | ***819.3843*** | 410.1958 | 802.3577 | 401.6825 | 801.3737 | 401.1905 | 8 |
| 5 | 608.3079 | 304.6576 | | | 590.2973 | 295.6523 | E | ***748.3472*** | 374.6772 | 731.3206 | 366.1639 | 730.3366 | 365.6719 | 7 |
| 6 | 723.3348 | 362.1710 | | | 705.3243 | 353.1658 | D | ***619.3046*** | 310.1559 | 602.2780 | 301.6427 | 601.2940 | 301.1506 | 6 |
| 7 | 822.4032 | 411.7053 | | | 804.3927 | 402.7000 | V | 504.2776 | 252.6425 | 487.2511 | 244.1292 | 486.2671 | 243.6372 | 5 |
| 8 | 879.4247 | 440.2160 | | | 861.4141 | 431.2107 | G | 405.2092 | 203.1082 | 388.1827 | 194.5950 | 387.1987 | 194.1030 | 4 |
| 9 | **966.4567** | **483.7320** | | | 948.4462 | 474.7267 | S | ***348.1878*** | 174.5975 | 331.1612 | 166.0842 | 330.1772 | 165.5922 | 3 |
| 10 | 1080.4997 | 540.7535 | 1063.4731 | 532.2402 | 1062.4891 | 531.7482 | N | ***261.1557*** | 131.0815 | 244.1292 | 122.5682 | | | 2 |
| 11 | | | | | | | K | 147.1128 | 74.0600 | 130.0863 | 65.5468 | | | 1 |

## F19/20

MS/MS Fragmentation of peptide **LVFAEDVGSNK**

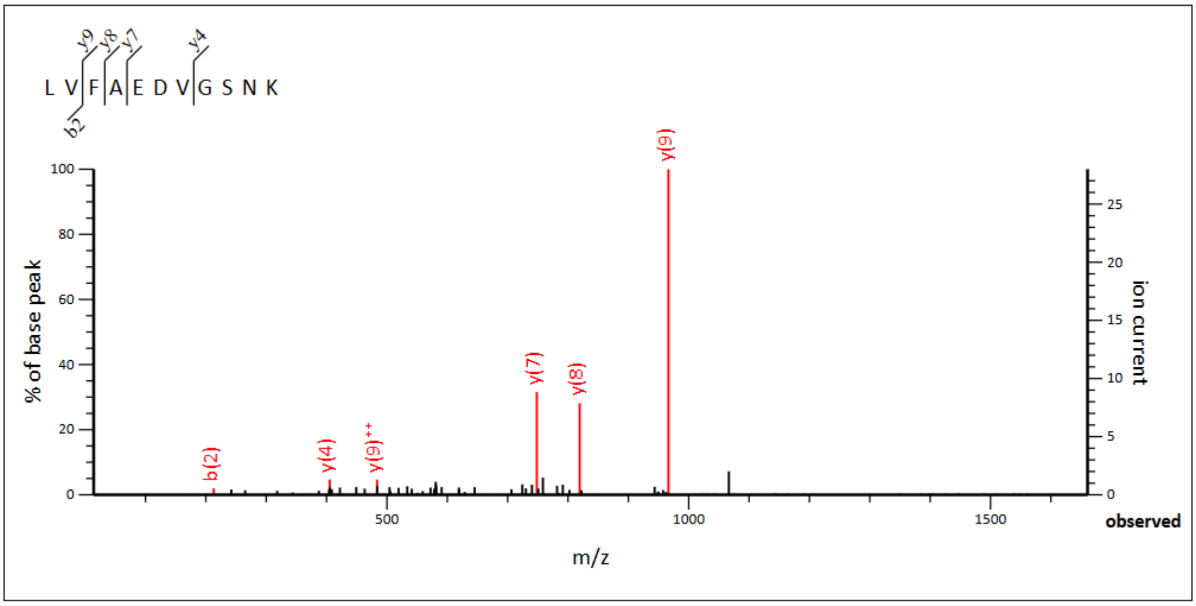


**Monoisotopic mass of neutral peptide Mr(calc):** 1177.5979
**Ions Score:** 26 **Expect:** 0.0025
**Matches :** 6/90 fragment ions using 11 most intense peaks

| # | b | $b^{++}$ | b* | $b^{*++}$ | $b^{0}$ | $b^{0++}$ | Seq. | y | $y^{++}$ | y* | $y^{*++}$ | $y^{0}$ | $y^{0++}$ | # |
|---|---|---|---|---|---|---|---|---|---|---|---|---|---|---|
| 1 | 114.0913 | 57.5493 | | | | | L | | | | | | | 11 |
| 2 | **213.1598** | 107.0835 | | | | | V | 1065.5211 | 533.2642 | 1048.4946 | 524.7509 | 1047.5106 | 524.2589 | 10 |
| 3 | 360.2282 | 180.6177 | | | | | F | ***966.4527*** | **483.7300** | 949.4262 | 475.2167 | 948.4421 | 474.7247 | 9 |
| 4 | 431.2653 | 216.1363 | | | | | A | ***819.3843*** | 410.1958 | 802.3577 | 401.6825 | 801.3737 | 401.1905 | 8 |
| 5 | 560.3079 | 280.6576 | | | 542.2973 | 271.6523 | E | ***748.3472*** | 374.6772 | 731.3206 | 366.1639 | 730.3366 | 365.6719 | 7 |
| 6 | 675.3348 | 338.1710 | | | 657.3243 | 329.1658 | D | 619.3046 | 310.1559 | 602.2780 | 301.6427 | 601.2940 | 301.1506 | 6 |
| 7 | 774.4032 | 387.7053 | | | 756.3927 | 378.7000 | V | 504.2776 | 252.6425 | 487.2511 | 244.1292 | 486.2671 | 243.6372 | 5 |
| 8 | 831.4247 | 416.2160 | | | 813.4141 | 407.2107 | G | ***405.2092*** | 203.1082 | 388.1827 | 194.5950 | 387.1987 | 194.1030 | 4 |
| 9 | 918.4567 | 459.7320 | | | 900.4462 | 450.7267 | S | 348.1878 | 174.5975 | 331.1612 | 166.0842 | 330.1772 | 165.5922 | 3 |
| 10 | 1032.4997 | 516.7535 | 1015.4731 | 508.2402 | 1014.4891 | 507.7482 | N | 261.1557 | 131.0815 | 244.1292 | 122.5682 | | | 2 |
| 11 | | | | | | | K | 147.1128 | 74.0600 | 130.0863 | 65.5468 | | | 1 |

## A21

MS/MS Fragmentation of peptide **LVFFEDVGSNK**

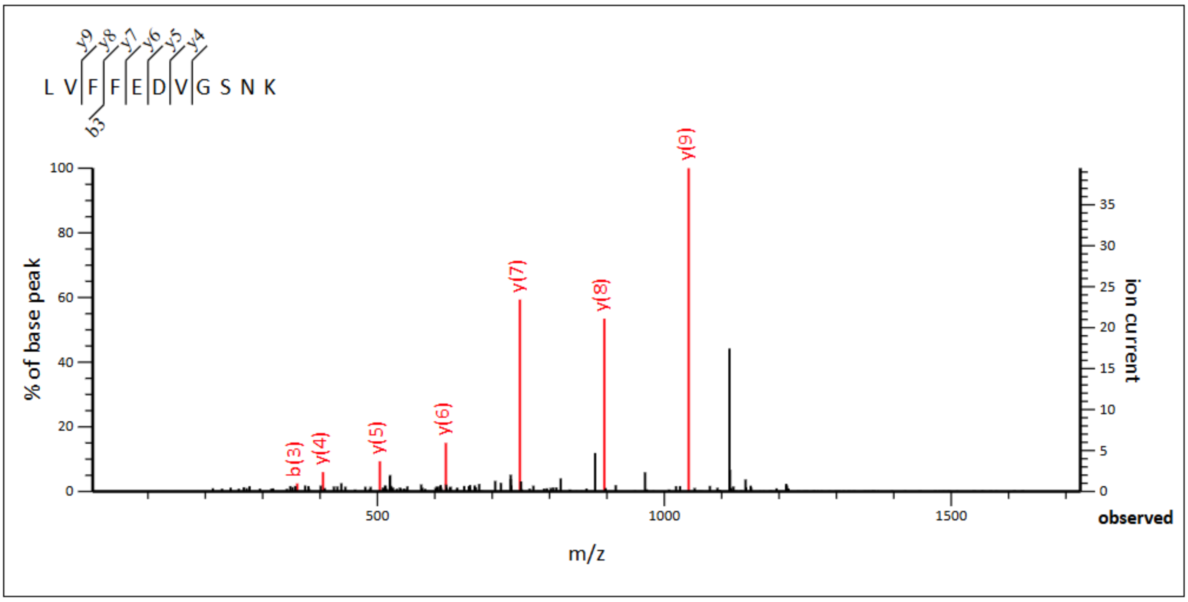


**Monoisotopic mass of neutral peptide Mr(calc):** 1253.6292
**Ions Score:** 42 **Expect:** 6.4e-05
**Matches :** 7/90 fragment ions using 12 most intense peaks

| # | b | $b^{++}$ | b* | $b^{*++}$ | $b^0$ | $b^{0++}$ | Seq. | y | $y^{++}$ | y* | $y^{*++}$ | $y^0$ | $y^{0++}$ | # |
|---|---|---|---|---|---|---|---|---|---|---|---|---|---|---|
| 1 | 114.0913 | 57.5493 | | | | | L | | | | | | | 11 |
| 2 | 213.1598 | 107.0835 | | | | | V | 1141.5524 | 571.2798 | 1124.5259 | 562.7666 | 1123.5419 | 562.2746 | 10 |
| 3 | **360.2282** | 180.6177 | | | | | F | ***1042.4840*** | 521.7456 | 1025.4575 | 513.2324 | 1024.4734 | 512.7404 | 9 |
| 4 | 507.2966 | 254.1519 | | | | | F | ***895.4156*** | 448.2114 | 878.3890 | 439.6982 | 877.4050 | 439.2061 | 8 |
| 5 | 636.3392 | 318.6732 | | | 618.3286 | 309.6679 | E | ***748.3472*** | 374.6772 | 731.3206 | 366.1639 | 730.3366 | 365.6719 | 7 |
| 6 | 751.3661 | 376.1867 | | | 733.3556 | 367.1814 | D | ***619.3046*** | 310.1559 | 602.2780 | 301.6427 | 601.2940 | 301.1506 | 6 |
| 7 | 850.4345 | 425.7209 | | | 832.4240 | 416.7156 | V | ***504.2776*** | 252.6425 | 487.2511 | 244.1292 | 486.2671 | 243.6372 | 5 |
| 8 | 907.4560 | 454.2316 | | | 889.4454 | 445.2264 | G | ***405.2092*** | 203.1082 | 388.1827 | 194.5950 | 387.1987 | 194.1030 | 4 |
| 9 | 994.4880 | 497.7477 | | | 976.4775 | 488.7424 | S | 348.1878 | 174.5975 | 331.1612 | 166.0842 | 330.1772 | 165.5922 | 3 |
| 10 | 1108.5310 | 554.7691 | 1091.5044 | 546.2558 | 1090.5204 | 545.7638 | N | 261.1557 | 131.0815 | 244.1292 | 122.5682 | | | 2 |
| 11 | | | | | | | K | 147.1128 | 74.0600 | 130.0863 | 65.5468 | | | 1 |

## G25

MS/MS Fragmentation of peptide **LVFFAEDVSNK**

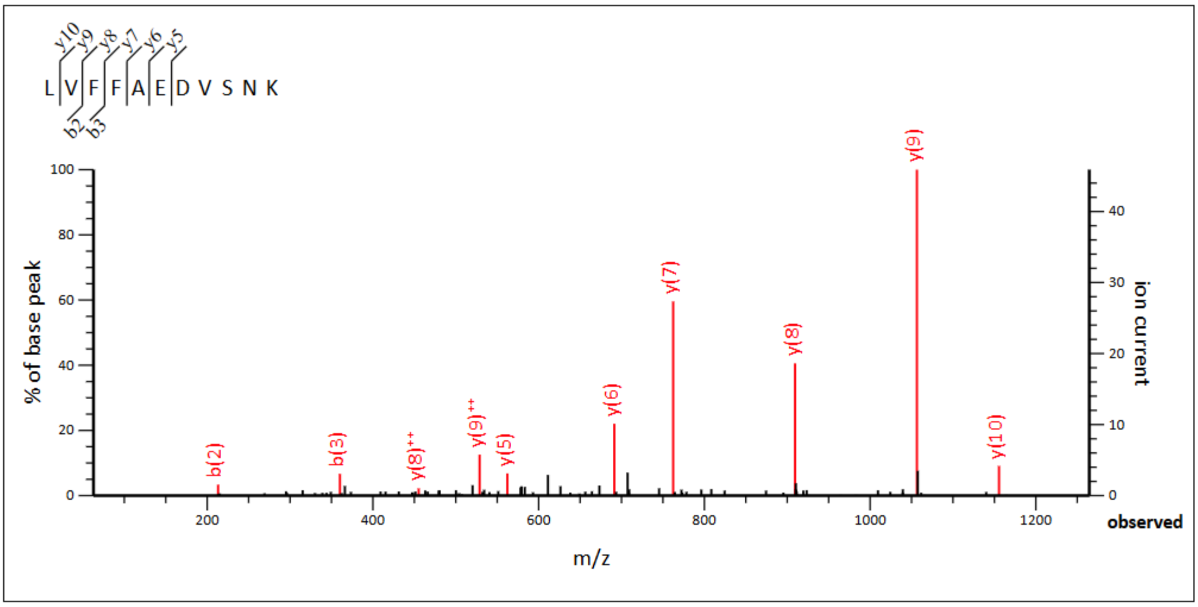


**Monoisotopic mass of neutral peptide Mr(calc):** 1267.6449
**Ions Score:** 36 **Expect:** 0.00024
**Matches :** 10/88 fragment ions using 21 most intense peaks

| # | b | $b^{++}$ | $b^{*}$ | $b^{*++}$ | $b^{0}$ | $b^{0++}$ | Seq. | y | $y^{++}$ | $y^{*}$ | $y^{*++}$ | $y^{0}$ | $y^{0++}$ | # |
|---|---|---|---|---|---|---|---|---|---|---|---|---|---|---|
| 1 | 114.0913 | 57.5493 | | | | | L | | | | | | | 11 |
| 2 | **213.1598** | 107.0835 | | | | | V | ***1155.5681*** | 578.2877 | 1138.5415 | 569.7744 | 1137.5575 | 569.2824 | 10 |
| 3 | **360.2282** | 180.6177 | | | | | F | ***1056.4997*** | **528.7535** | 1039.4731 | 520.2402 | 1038.4891 | 519.7482 | 9 |
| 4 | 507.2966 | 254.1519 | | | | | F | ***909.4312*** | **455.2193** | 892.4047 | 446.7060 | 891.4207 | 446.2140 | 8 |
| 5 | 578.3337 | 289.6705 | | | | | A | ***762.3628*** | 381.6850 | 745.3363 | 373.1718 | 744.3523 | 372.6798 | 7 |
| 6 | 707.3763 | 354.1918 | | | 689.3657 | 345.1865 | E | ***691.3257*** | 346.1665 | 674.2992 | 337.6532 | 673.3151 | 337.1612 | 6 |
| 7 | 822.4032 | 411.7053 | | | 804.3927 | 402.7000 | D | ***562.2831*** | 281.6452 | 545.2566 | 273.1319 | 544.2726 | 272.6399 | 5 |
| 8 | 921.4716 | 461.2395 | | | 903.4611 | 452.2342 | V | 447.2562 | 224.1317 | 430.2296 | 215.6184 | 429.2456 | 215.1264 | 4 |
| 9 | 1008.5037 | 504.7555 | | | 990.4931 | 495.7502 | S | 348.1878 | 174.5975 | 331.1612 | 166.0842 | 330.1772 | 165.5922 | 3 |
| 10 | 1122.5466 | 561.7769 | 1105.5201 | 553.2637 | 1104.5360 | 552.7717 | N | 261.1557 | 131.0815 | 244.1292 | 122.5682 | | | 2 |
| 11 | | | | | | | K | 147.1128 | 74.0600 | 130.0863 | 65.5468 | | | 1 |

## S26

MS/MS Fragmentation of peptide **LVFFAEDVGNK**

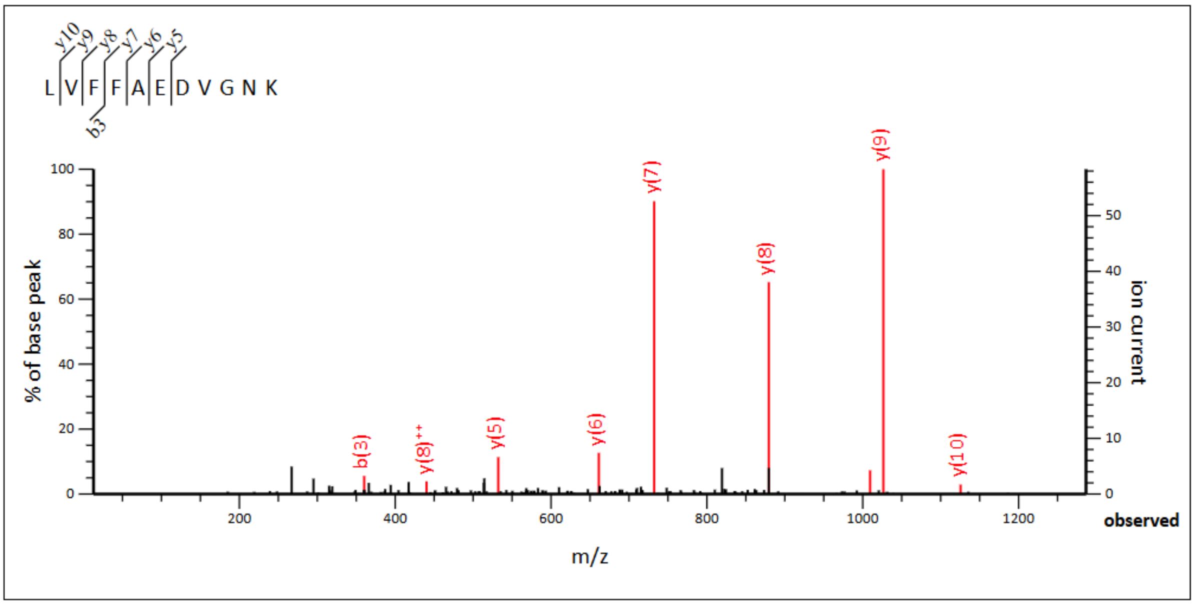


**Monoisotopic mass of neutral peptide Mr(calc):** 1237.6343
**Ions Score:** 53 **Expect:** 5.1e-06
**Matches :** 9/84 fragment ions using 11 most intense peaks

| # | b | $b^{++}$ | b* | $b^{*++}$ | $b^0$ | $b^{0++}$ | Seq. | y | $y^{++}$ | y* | $y^{*++}$ | $y^0$ | $y^{0++}$ | # |
|---|---|---|---|---|---|---|---|---|---|---|---|---|---|---|
| 1 | 114.0913 | 57.5493 | | | | | L | | | | | | | 11 |
| 2 | 213.1598 | 107.0835 | | | | | V | 1125.5575 | 563.2824 | 1108.5310 | 554.7691 | 1107.5469 | 554.2771 | 10 |
| 3 | 360.2282 | 180.6177 | | | | | F | 1026.4891 | 513.7482 | 1009.4625 | 505.2349 | 1008.4785 | 504.7429 | 9 |
| 4 | 507.2966 | 254.1519 | | | | | F | 879.4207 | 440.2140 | 862.3941 | 431.7007 | 861.4101 | 431.2087 | 8 |
| 5 | 578.3337 | 289.6705 | | | | | A | 732.3523 | 366.6798 | 715.3257 | 358.1665 | 714.3417 | 357.6745 | 7 |
| 6 | 707.3763 | 354.1918 | | | 689.3657 | 345.1865 | E | 661.3151 | 331.1612 | 644.2886 | 322.6479 | 643.3046 | 322.1559 | 6 |
| 7 | 822.4032 | 411.7053 | | | 804.3927 | 402.7000 | D | 532.2726 | 266.6399 | 515.2460 | 258.1266 | 514.2620 | 257.6346 | 5 |
| 8 | 921.4716 | 461.2395 | | | 903.4611 | 452.2342 | V | 417.2456 | 209.1264 | 400.2191 | 200.6132 | | | 4 |
| 9 | 978.4931 | 489.7502 | | | 960.4825 | 480.7449 | G | 318.1772 | 159.5922 | 301.1506 | 151.0790 | | | 3 |
| 10 | 1092.5360 | 546.7717 | 1075.5095 | 538.2584 | 1074.5255 | 537.7664 | N | 261.1557 | 131.0815 | 244.1292 | 122.5682 | | | 2 |
| 11 | | | | | | | K | 147.1128 | 74.0600 | 130.0863 | 65.5468 | | | 1 |

## N27

MS/MS Fragmentation of peptide **LVFFAEDVGSK**

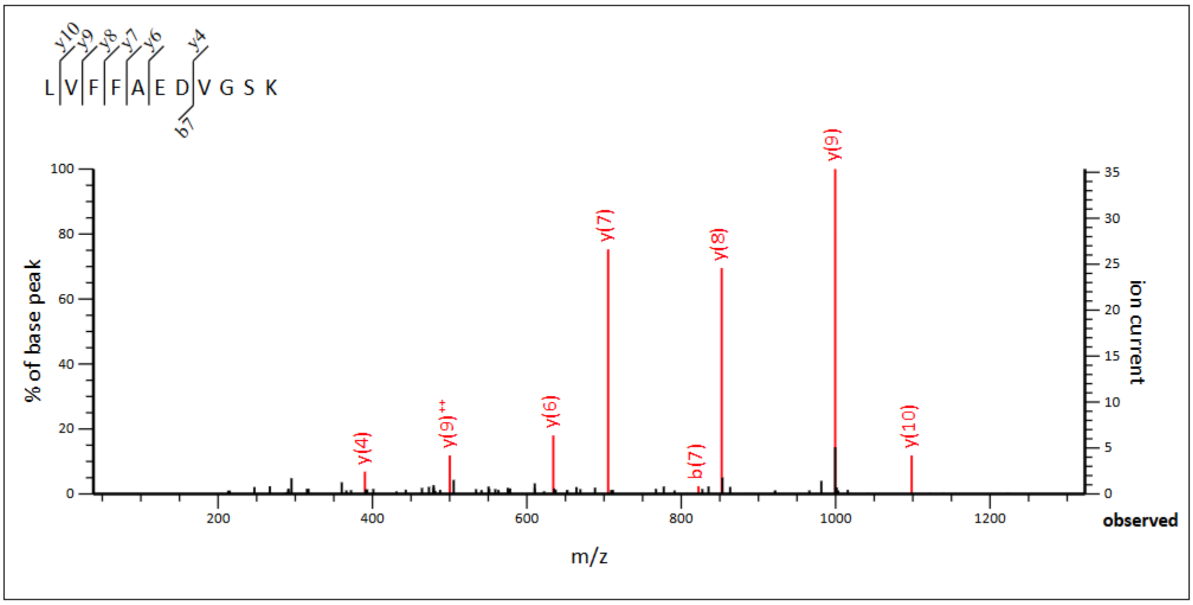


**Monoisotopic mass of neutral peptide Mr(calc):** 1210.6234
**Ions Score:** 51 **Expect:** 8.6e-06
**Matches :** 8/88 fragment ions using 11 most intense peaks

| # | b | $b^{++}$ | $b^0$ | $b^{0++}$ | Seq. | y | $y^{++}$ | $y^*$ | $y^{*++}$ | $y^0$ | $y^{0++}$ | # |
|---|---|---|---|---|---|---|---|---|---|---|---|---|
| 1 | 114.0913 | 57.5493 | | | L | | | | | | | 11 |
| 2 | 213.1598 | 107.0835 | | | V | ***1098.5466*** | 549.7769 | 1081.5201 | 541.2637 | 1080.5360 | 540.7717 | 10 |
| 3 | 360.2282 | 180.6177 | | | F | ***999.4782*** | **500.2427** | 982.4516 | 491.7295 | 981.4676 | 491.2374 | 9 |
| 4 | 507.2966 | 254.1519 | | | F | ***852.4098*** | 426.7085 | 835.3832 | 418.1953 | 834.3992 | 417.7032 | 8 |
| 5 | 578.3337 | 289.6705 | | | A | ***705.3414*** | 353.1743 | 688.3148 | 344.6610 | 687.3308 | 344.1690 | 7 |
| 6 | 707.3763 | 354.1918 | 689.3657 | 345.1865 | E | ***634.3042*** | 317.6558 | 617.2777 | 309.1425 | 616.2937 | 308.6505 | 6 |
| 7 | **822.4032** | 411.7053 | 804.3927 | 402.7000 | D | 505.2617 | 253.1345 | 488.2351 | 244.6212 | 487.2511 | 244.1292 | 5 |
| 8 | 921.4716 | 461.2395 | 903.4611 | 452.2342 | V | ***390.2347*** | 195.6210 | 373.2082 | 187.1077 | 372.2241 | 186.6157 | 4 |
| 9 | 978.4931 | 489.7502 | 960.4825 | 480.7449 | G | 291.1663 | 146.0868 | 274.1397 | 137.5735 | 273.1557 | 137.0815 | 3 |
| 10 | 1065.5251 | 533.2662 | 1047.5146 | 524.2609 | S | 234.1448 | 117.5761 | 217.1183 | 109.0628 | 216.1343 | 108.5708 | 2 |
| 11 | | | | | K | 147.1128 | 74.0600 | 130.0863 | 65.5468 | | | 1 |

### DUPLICATION Aβ42 peptide detections

## In s1Aβ42

### Y10

MS/MS Fragmentation of peptide **HDSGYYEVHHQK**

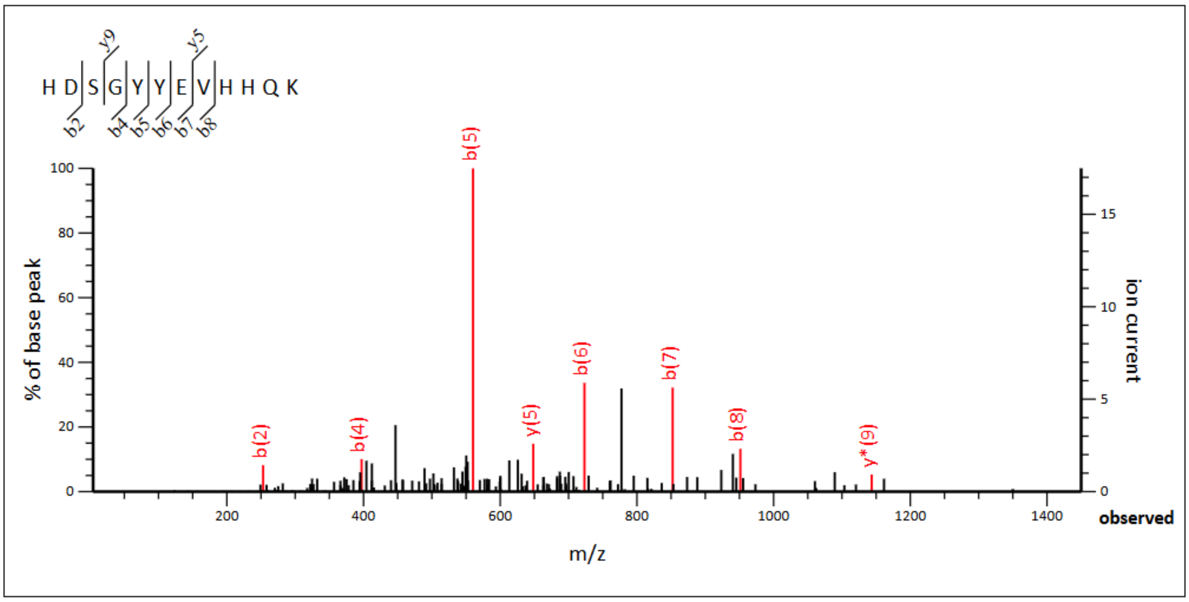


**Monoisotopic mass of neutral peptide Mr(calc):** 1498.6589
**Ions Score:** 41 **Expect:** 7.3e-05
**Matches :** 8/100 fragment ions using 12 most intense peaks

| # | b | $b^{++}$ | $b^{*}$ | $b^{*++}$ | $b^{0}$ | $b^{0++}$ | Seq. | y | $y^{++}$ | $y^{*}$ | $y^{*++}$ | $y^{0}$ | $y^{0++}$ | # |
|---|---|---|---|---|---|---|---|---|---|---|---|---|---|---|
| 1 | 138.0662 | 69.5367 | | | | | H | | | | | | | 12 |
| 2 | ***253.0931*** | 127.0502 | | | 235.0826 | 118.0449 | D | 1362.6073 | 681.8073 | 1345.5808 | 673.2940 | 1344.5967 | 672.8020 | 11 |
| 3 | 340.1252 | 170.5662 | | | 322.1146 | 161.5609 | S | 1247.5804 | 624.2938 | 1230.5538 | 615.7805 | 1229.5698 | 615.2885 | 10 |
| 4 | ***397.1466*** | 199.0769 | | | 379.1361 | 190.0717 | G | 1160.5483 | 580.7778 | 1143.5218 | 572.2645 | 1142.5378 | 571.7725 | 9 |
| 5 | ***560.2100*** | 280.6086 | | | 542.1994 | 271.6033 | Y | 1103.5269 | 552.2671 | 1086.5003 | 543.7538 | 1085.5163 | 543.2618 | 8 |
| 6 | ***723.2733*** | 362.1403 | | | 705.2627 | 353.1350 | Y | 940.4635 | 470.7354 | 923.4370 | 462.2221 | 922.4530 | 461.7301 | 7 |
| 7 | ***852.3159*** | 426.6616 | | | 834.3053 | 417.6563 | E | 777.4002 | 389.2037 | 760.3737 | 380.6905 | 759.3896 | 380.1985 | 6 |
| 8 | ***951.3843*** | 476.1958 | | | 933.3737 | 467.1905 | V | **648.3576** | 324.6824 | 631.3311 | 316.1692 | | | 5 |
| 9 | 1088.4432 | 544.7252 | | | 1070.4326 | 535.7200 | H | 549.2892 | 275.1482 | 532.2627 | 266.6350 | | | 4 |
| 10 | 1225.5021 | 613.2547 | | | 1207.4915 | 604.2494 | H | 412.2303 | 206.6188 | 395.2037 | 198.1055 | | | 3 |
| 11 | 1353.5607 | 677.2840 | 1336.5341 | 668.7707 | 1335.5501 | 668.2787 | Q | 275.1714 | 138.0893 | 258.1448 | 129.5761 | | | 2 |
| 12 | | | | | | | K | 147.1128 | 74.0600 | 130.0863 | 65.5468 | | | 1 |

## V12

MS/MS Fragmentation of peptide **HDSGYEVVHHQK**

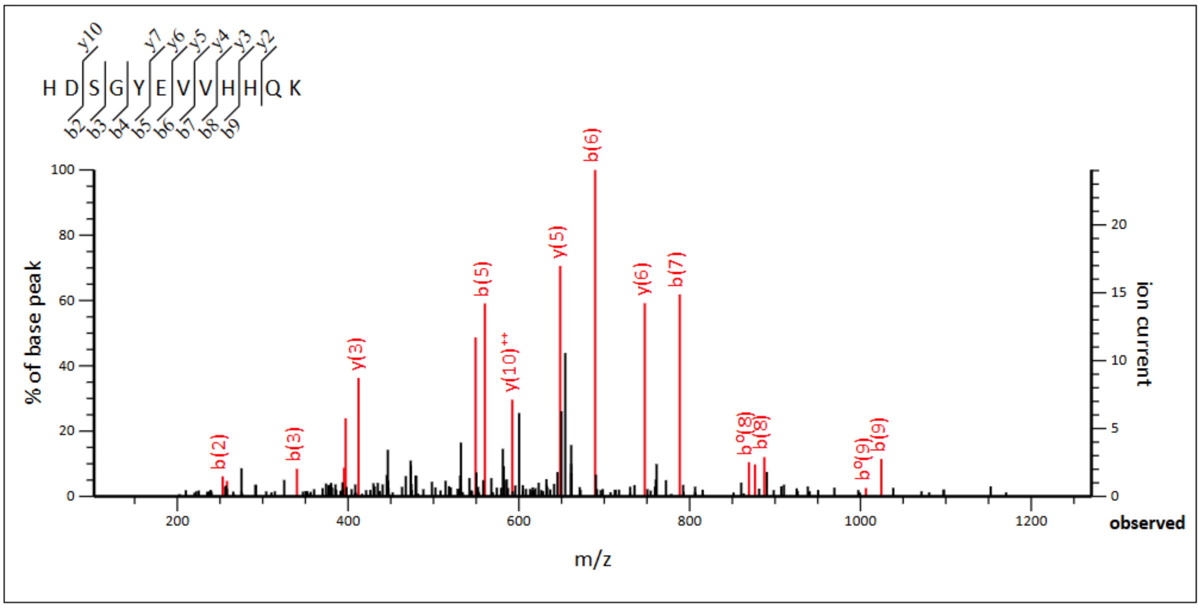


**Monoisotopic mass of neutral peptide Mr(calc):** 1434.6640
**Ions Score:** 56 **Expect:** 2.8e-06
**Matches :** 18/98 fragment ions using 29 most intense peaks

| # | b | $b^{++}$ | $b^{*}$ | $b^{*++}$ | $b^{0}$ | $b^{0++}$ | Seq. | y | $y^{++}$ | $y^{*}$ | $y^{*++}$ | $y^{0}$ | $y^{0++}$ | # |
|---|---|---|---|---|---|---|---|---|---|---|---|---|---|---|
| 1 | 138.0662 | 69.5367 | | | | | H | | | | | | | 12 |
| 2 | ***253.0931*** | 127.0502 | | | 235.0826 | 118.0449 | D | 1298.6124 | 649.8098 | 1281.5858 | 641.2966 | 1280.6018 | 640.8046 | 11 |
| 3 | ***340.1252*** | 170.5662 | | | 322.1146 | 161.5609 | S | 1183.5854 | **592.2964** | 1166.5589 | 583.7831 | 1165.5749 | 583.2911 | 10 |
| 4 | ***397.1466*** | 199.0769 | | | 379.1361 | 190.0717 | G | 1096.5534 | 548.7803 | 1079.5269 | 540.2671 | 1078.5429 | 539.7751 | 9 |
| 5 | ***560.2100*** | 280.6086 | | | 542.1994 | 271.6033 | Y | 1039.5320 | 520.2696 | 1022.5054 | 511.7563 | 1021.5214 | 511.2643 | 8 |
| 6 | ***689.2525*** | 345.1299 | | | 671.2420 | 336.1246 | E | **876.4686** | 438.7380 | 859.4421 | 430.2247 | 858.4581 | 429.7327 | 7 |
| 7 | ***788.3210*** | 394.6641 | | | 770.3104 | 385.6588 | V | **747.4260** | 374.2167 | 730.3995 | 365.7034 | | | 6 |
| 8 | ***887.3894*** | 444.1983 | | | 869.3788 | 435.1930 | V | **648.3576** | 324.6824 | 631.3311 | 316.1692 | | | 5 |
| 9 | ***1024.4483*** | 512.7278 | | | 1006.4377 | 503.7225 | H | **549.2892** | 275.1482 | 532.2627 | 266.6350 | | | 4 |
| 10 | 1161.5072 | 581.2572 | | | 1143.4966 | 572.2520 | H | **412.2303** | 206.6188 | 395.2037 | 198.1055 | | | 3 |
| 11 | 1289.5658 | 645.2865 | 1272.5392 | 636.7733 | 1271.5552 | 636.2812 | Q | 275.1714 | 138.0893 | 258.1448 | 129.5761 | | | 2 |
| 12 | | | | | | | K | 147.1128 | 74.0600 | 130.0863 | 65.5468 | | | 1 |

## L17

MS/MS Fragmentation of peptide **LLVFFAEDVGSNK**

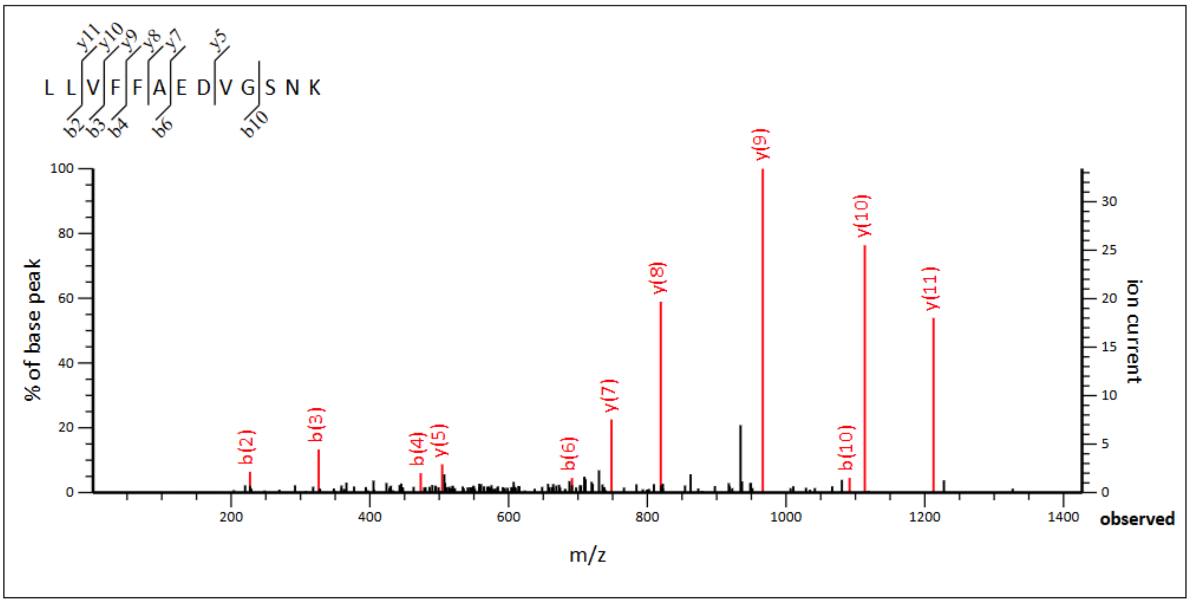


**Monoisotopic mass of neutral peptide Mr(calc):** 1437.7504
**Ions Score:** 55 **Expect:** 3e-06
**Matches :** 11/106 fragment ions using 13 most intense peaks

| # | b | $b^{++}$ | b* | $b^{*++}$ | $b^0$ | $b^{0++}$ | Seq. | y | $y^{++}$ | y* | $y^{*++}$ | $y^0$ | $y^{0++}$ | # |
|---|---|---|---|---|---|---|---|---|---|---|---|---|---|---|
| 1 | 114.0913 | 57.5493 | | | | | L | | | | | | | 13 |
| 2 | *227.1754* | 114.0913 | | | | | L | 1325.6736 | 663.3404 | 1308.6470 | 654.8272 | 1307.6630 | 654.3352 | 12 |
| 3 | *326.2438* | 163.6255 | | | | | V | *1212.5895* | 606.7984 | 1195.5630 | 598.2851 | 1194.5790 | 597.7931 | 11 |
| 4 | *473.3122* | 237.1598 | | | | | F | *1113.5211* | 557.2642 | 1096.4946 | 548.7509 | 1095.5106 | 548.2589 | 10 |
| 5 | 620.3806 | 310.6940 | | | | | F | *966.4527* | 483.7300 | 949.4262 | 475.2167 | 948.4421 | 474.7247 | 9 |
| 6 | *691.4178* | 346.2125 | | | | | A | *819.3843* | 410.1958 | 802.3577 | 401.6825 | 801.3737 | 401.1905 | 8 |
| 7 | 820.4604 | 410.7338 | | | 802.4498 | 401.7285 | E | *748.3472* | 374.6772 | 731.3206 | 366.1639 | 730.3366 | 365.6719 | 7 |
| 8 | 935.4873 | 468.2473 | | | 917.4767 | 459.2420 | D | 619.3046 | 310.1559 | 602.2780 | 301.6427 | 601.2940 | 301.1506 | 6 |
| 9 | 1034.5557 | 517.7815 | | | 1016.5451 | 508.7762 | V | *504.2776* | 252.6425 | 487.2511 | 244.1292 | 486.2671 | 243.6372 | 5 |
| 10 | *1091.5772* | 546.2922 | | | 1073.5666 | 537.2869 | G | 405.2092 | 203.1082 | 388.1827 | 194.5950 | 387.1987 | 194.1030 | 4 |
| 11 | 1178.6092 | 589.8082 | | | 1160.5986 | 580.8030 | S | 348.1878 | 174.5975 | 331.1612 | 166.0842 | 330.1772 | 165.5922 | 3 |
| 12 | 1292.6521 | 646.8297 | 1275.6256 | 638.3164 | 1274.6416 | 637.8244 | N | 261.1557 | 131.0815 | 244.1292 | 122.5682 | | | 2 |
| 13 | | | | | | | K | 147.1128 | 74.0600 | 130.0863 | 65.5468 | | | 1 |

## V18

MS/MS Fragmentation of peptide **LVVFFAEDVGSNK**

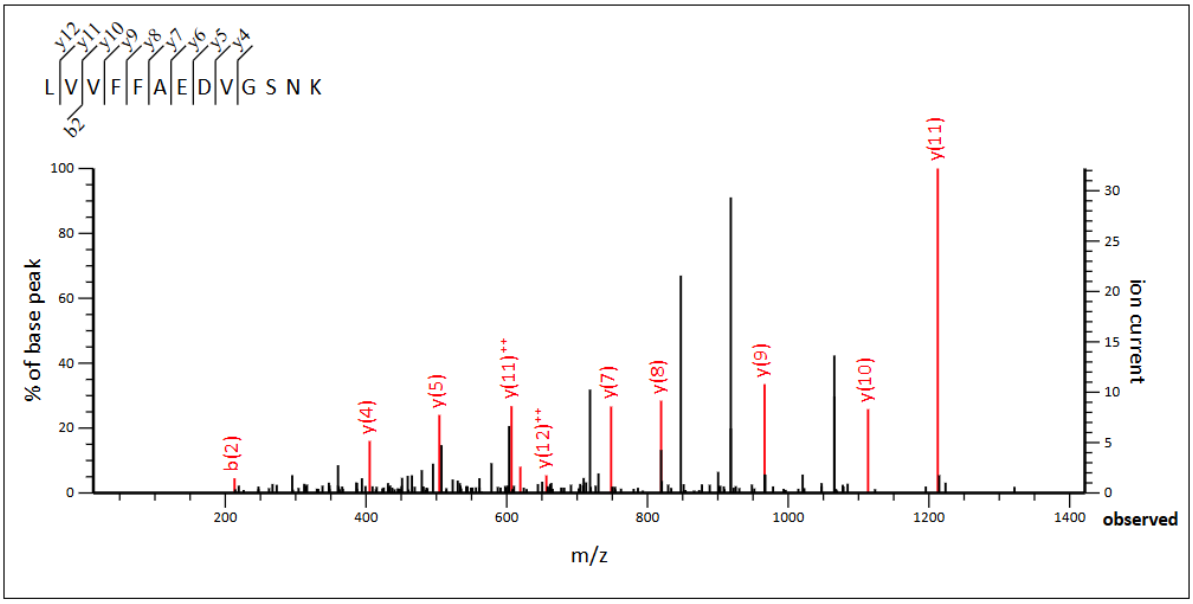


**Monoisotopic mass of neutral peptide Mr(calc):** 1423.7347
**Ions Score:** 45 **Expect:** 3.2e-05
**Matches :** 11/106 fragment ions using 25 most intense peaks

| # | b | $b^{++}$ | b* | $b^{*++}$ | $b^{0}$ | $b^{0++}$ | Seq. | y | $y^{++}$ | y* | $y^{*++}$ | $y^{0}$ | $y^{0++}$ | # |
|---|---|---|---|---|---|---|---|---|---|---|---|---|---|---|
| 1 | 114.0913 | 57.5493 | | | | | L | | | | | | | 13 |
| 2 | **213.1598** | 107.0835 | | | | | V | 1311.6579 | **656.3326** | 1294.6314 | 647.8193 | 1293.6474 | 647.3273 | 12 |
| 3 | 312.2282 | 156.6177 | | | | | V | ***1212.5895*** | **606.7984** | 1195.5630 | 598.2851 | 1194.5790 | 597.7931 | 11 |
| 4 | 459.2966 | 230.1519 | | | | | F | ***1113.5211*** | 557.2642 | 1096.4946 | 548.7509 | 1095.5106 | 548.2589 | 10 |
| 5 | 606.3650 | 303.6861 | | | | | F | ***966.4527*** | 483.7300 | 949.4262 | 475.2167 | 948.4421 | 474.7247 | 9 |
| 6 | 677.4021 | 339.2047 | | | | | A | ***819.3843*** | 410.1958 | 802.3577 | 401.6825 | 801.3737 | 401.1905 | 8 |
| 7 | 806.4447 | 403.7260 | | | 788.4341 | 394.7207 | E | ***748.3472*** | 374.6772 | 731.3206 | 366.1639 | 730.3366 | 365.6719 | 7 |
| 8 | 921.4716 | 461.2395 | | | 903.4611 | 452.2342 | D | ***619.3046*** | 310.1559 | 602.2780 | 301.6427 | 601.2940 | 301.1506 | 6 |
| 9 | 1020.5401 | 510.7737 | | | 1002.5295 | 501.7684 | V | ***504.2776*** | 252.6425 | 487.2511 | 244.1292 | 486.2671 | 243.6372 | 5 |
| 10 | 1077.5615 | 539.2844 | | | 1059.5510 | 530.2791 | G | ***405.2092*** | 203.1082 | 388.1827 | 194.5950 | 387.1987 | 194.1030 | 4 |
| 11 | 1164.5936 | 582.8004 | | | 1146.5830 | 573.7951 | S | 348.1878 | 174.5975 | 331.1612 | 166.0842 | 330.1772 | 165.5922 | 3 |
| 12 | 1278.6365 | 639.8219 | 1261.6099 | 631.3086 | 1260.6259 | 630.8166 | N | 261.1557 | 131.0815 | 244.1292 | 122.5682 | | | 2 |
| 13 | | | | | | | K | 147.1128 | 74.0600 | 130.0863 | 65.5468 | | | 1 |

## F19/20

MS/MS Fragmentation of peptide **LVFFFAEDVGSNK**

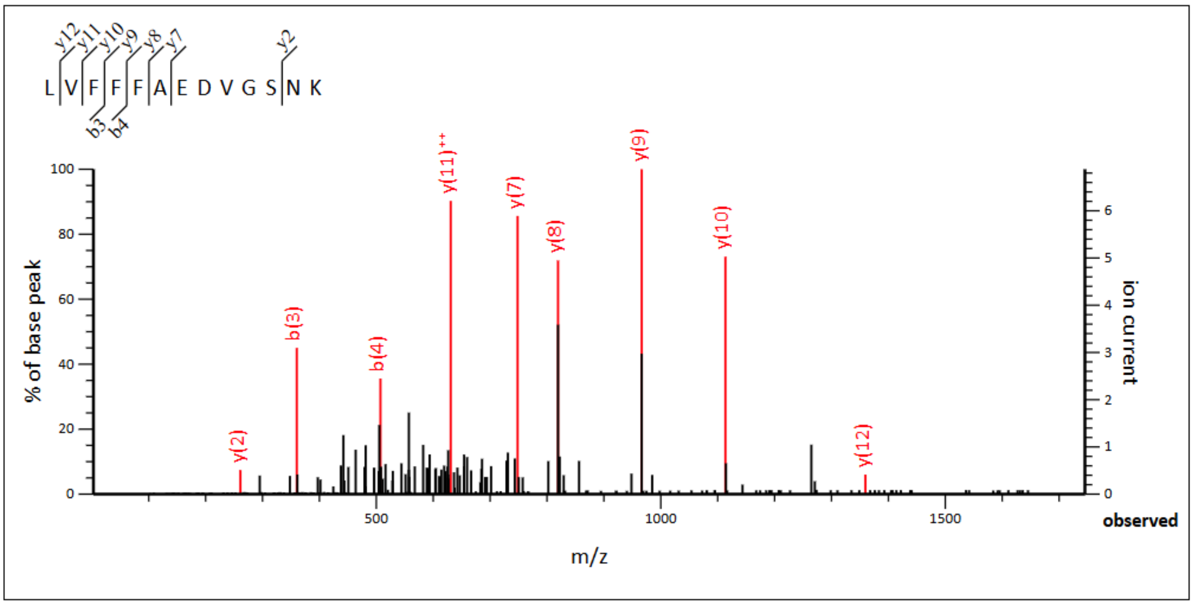


**Monoisotopic mass of neutral peptide Mr(calc):** 1471.7347
**Ions Score:** 40 **Expect:** 9.7e-05
**Matches :** 9/106 fragment ions using 14 most intense peaks

| # | b | $b^{++}$ | $b^{*}$ | $b^{*++}$ | $b^{0}$ | $b^{0++}$ | Seq. | y | $y^{++}$ | $y^{*}$ | $y^{*++}$ | $y^{0}$ | $y^{0++}$ | # |
|---|---|---|---|---|---|---|---|---|---|---|---|---|---|---|
| 1 | 114.0913 | 57.5493 | | | | | L | | | | | | | 13 |
| 2 | 213.1598 | 107.0835 | | | | | V | ***1359.6579*** | 680.3326 | 1342.6314 | 671.8193 | 1341.6474 | 671.3273 | 12 |
| 3 | **360.2282** | 180.6177 | | | | | F | 1260.5895 | **630.7984** | 1243.5630 | 622.2851 | 1242.5790 | 621.7931 | 11 |
| 4 | **507.2966** | 254.1519 | | | | | F | ***1113.5211*** | 557.2642 | 1096.4946 | 548.7509 | 1095.5106 | 548.2589 | 10 |
| 5 | 654.3650 | 327.6861 | | | | | F | ***966.4527*** | 483.7300 | 949.4262 | 475.2167 | 948.4421 | 474.7247 | 9 |
| 6 | 725.4021 | 363.2047 | | | | | A | ***819.3843*** | 410.1958 | 802.3577 | 401.6825 | 801.3737 | 401.1905 | 8 |
| 7 | 854.4447 | 427.7260 | | | 836.4341 | 418.7207 | E | ***748.3472*** | 374.6772 | 731.3206 | 366.1639 | 730.3366 | 365.6719 | 7 |
| 8 | 969.4716 | 485.2395 | | | 951.4611 | 476.2342 | D | 619.3046 | 310.1559 | 602.2780 | 301.6427 | 601.2940 | 301.1506 | 6 |
| 9 | 1068.5401 | 534.7737 | | | 1050.5295 | 525.7684 | V | 504.2776 | 252.6425 | 487.2511 | 244.1292 | 486.2671 | 243.6372 | 5 |
| 10 | 1125.5615 | 563.2844 | | | 1107.5510 | 554.2791 | G | 405.2092 | 203.1082 | 388.1827 | 194.5950 | 387.1987 | 194.1030 | 4 |
| 11 | 1212.5936 | 606.8004 | | | 1194.5830 | 597.7951 | S | 348.1878 | 174.5975 | 331.1612 | 166.0842 | 330.1772 | 165.5922 | 3 |
| 12 | 1326.6365 | 663.8219 | 1309.6099 | 655.3086 | 1308.6259 | 654.8166 | N | ***261.1557*** | 131.0815 | 244.1292 | 122.5682 | | | 2 |
| 13 | | | | | | | K | 147.1128 | 74.0600 | 130.0863 | 65.5468 | | | 1 |

## A21

MS/MS Fragmentation of peptide **LVFFAAEDVGSNK**

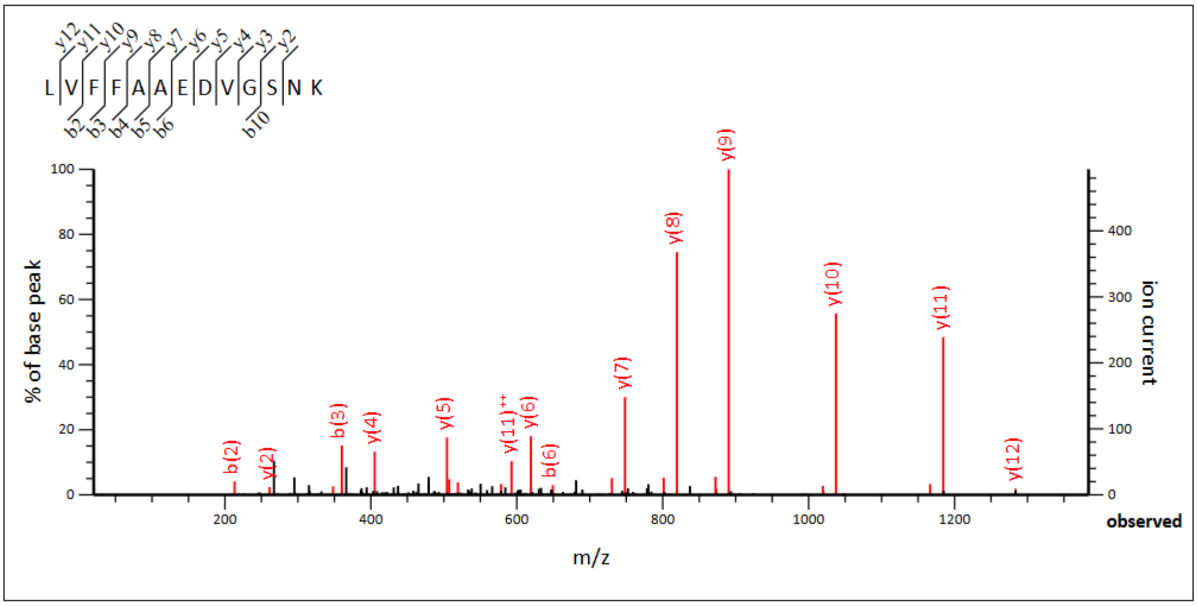


**Monoisotopic mass of neutral peptide Mr(calc):** 1395.7034
**Ions Score:** 85 **Expect:** 3.1e-09
**Matches :** 26/106 fragment ions using 45 most intense peaks

| # | b | b$^{++}$ | b* | b$^{*++}$ | b$^{0}$ | b$^{0++}$ | Seq. | y | y$^{++}$ | y* | y$^{*++}$ | y$^{0}$ | y$^{0++}$ | # |
|---|---|---|---|---|---|---|---|---|---|---|---|---|---|---|
| 1 | 114.0913 | 57.5493 | | | | | L | | | | | | | 13 |
| 2 | **213.1598** | 107.0835 | | | | | V | ***1283.6266*** | 642.3170 | 1266.6001 | 633.8037 | 1265.6161 | 633.3117 | 12 |
| 3 | **360.2282** | 180.6177 | | | | | F | ***1184.5582*** | **592.7828** | 1167.5317 | 584.2695 | 1166.5477 | 583.7775 | 11 |
| 4 | **507.2966** | 254.1519 | | | | | F | ***1037.4898*** | **519.2485** | 1020.4633 | 510.7353 | 1019.4793 | 510.2433 | 10 |
| 5 | **578.3337** | 289.6705 | | | | | A | ***890.4214*** | 445.7143 | 873.3949 | 437.2011 | 872.4108 | 436.7091 | 9 |
| 6 | **649.3708** | 325.1890 | | | | | A | ***819.3843*** | 410.1958 | 802.3577 | 401.6825 | 801.3737 | 401.1905 | 8 |
| 7 | 778.4134 | 389.7103 | | | 760.4028 | 380.7051 | E | ***748.3472*** | 374.6772 | 731.3206 | 366.1639 | 730.3366 | 365.6719 | 7 |
| 8 | 893.4403 | 447.2238 | | | 875.4298 | 438.2185 | D | ***619.3046*** | 310.1559 | 602.2780 | 301.6427 | 601.2940 | 301.1506 | 6 |
| 9 | 992.5088 | 496.7580 | | | 974.4982 | 487.7527 | V | ***504.2776*** | 252.6425 | 487.2511 | 244.1292 | 486.2671 | 243.6372 | 5 |
| 10 | **1049.5302** | 525.2688 | | | 1031.5197 | 516.2635 | G | ***405.2092*** | 203.1082 | 388.1827 | 194.5950 | 387.1987 | 194.1030 | 4 |
| 11 | 1136.5623 | 568.7848 | | | 1118.5517 | 559.7795 | S | ***348.1878*** | 174.5975 | 331.1612 | 166.0842 | 330.1772 | 165.5922 | 3 |
| 12 | 1250.6052 | 625.8062 | 1233.5786 | 617.2930 | 1232.5946 | 616.8009 | N | ***261.1557*** | 131.0815 | 244.1292 | 122.5682 | | | 2 |
| 13 | | | | | | | K | 147.1128 | 74.0600 | 130.0863 | 65.5468 | | | 1 |

## E22

MS/MS Fragmentation of peptide **LVFFAEEDVGSNK**

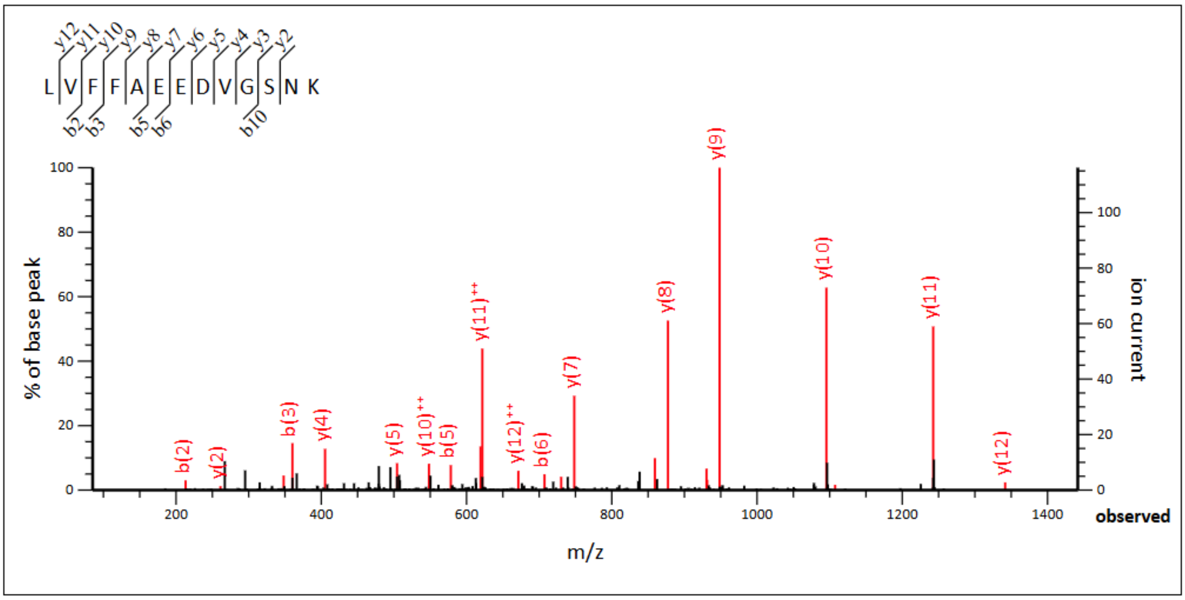


**Monoisotopic mass of neutral peptide Mr(calc):** 1453.7089
**Ions Score:** 76 **Expect:** 2.3e-08
**Matches :** 23/108 fragment ions using 45 most intense peaks

| # | b | $b^{++}$ | b* | $b^{*++}$ | $b^0$ | $b^{0++}$ | Seq. | y | $y^{++}$ | y* | $y^{*++}$ | $y^0$ | $y^{0++}$ | # |
|---|---|---|---|---|---|---|---|---|---|---|---|---|---|---|
| 1 | 114.0913 | 57.5493 | | | | | L | | | | | | | 13 |
| 2 | **213.1598** | 107.0835 | | | | | V | ***1341.6321*** | **671.3197** | 1324.6056 | 662.8064 | 1323.6216 | 662.3144 | 12 |
| 3 | **360.2282** | 180.6177 | | | | | F | ***1242.5637*** | **621.7855** | 1225.5372 | 613.2722 | 1224.5531 | 612.7802 | 11 |
| 4 | 507.2966 | 254.1519 | | | | | F | ***1095.4953*** | **548.2513** | 1078.4687 | 539.7380 | 1077.4847 | 539.2460 | 10 |
| 5 | **578.3337** | 289.6705 | | | | | A | ***948.4269*** | 474.7171 | 931.4003 | 466.2038 | 930.4163 | 465.7118 | 9 |
| 6 | **707.3763** | 354.1918 | | | 689.3657 | 345.1865 | E | ***877.3898*** | 439.1985 | 860.3632 | 430.6852 | 859.3792 | 430.1932 | 8 |
| 7 | 836.4189 | 418.7131 | | | 818.4083 | 409.7078 | E | ***748.3472*** | 374.6772 | 731.3206 | 366.1639 | 730.3366 | 365.6719 | 7 |
| 8 | 951.4458 | 476.2266 | | | 933.4353 | 467.2213 | D | ***619.3046*** | 310.1559 | 602.2780 | 301.6427 | 601.2940 | 301.1506 | 6 |
| 9 | 1050.5142 | 525.7608 | | | 1032.5037 | 516.7555 | V | ***504.2776*** | 252.6425 | 487.2511 | 244.1292 | 486.2671 | 243.6372 | 5 |
| 10 | **1107.5357** | 554.2715 | | | 1089.5251 | 545.2662 | G | ***405.2092*** | 203.1082 | 388.1827 | 194.5950 | 387.1987 | 194.1030 | 4 |
| 11 | 1194.5677 | 597.7875 | | | 1176.5572 | 588.7822 | S | ***348.1878*** | 174.5975 | 331.1612 | 166.0842 | 330.1772 | 165.5922 | 3 |
| 12 | 1308.6107 | 654.8090 | 1291.5841 | 646.2957 | 1290.6001 | 645.8037 | N | ***261.1557*** | 131.0815 | 244.1292 | 122.5682 | | | 2 |
| 13 | | | | | | | K | 147.1128 | 74.0600 | 130.0863 | 65.5468 | | | 1 |

## V24

MS/MS Fragmentation of peptide **LVFFAEDVVGSNK**

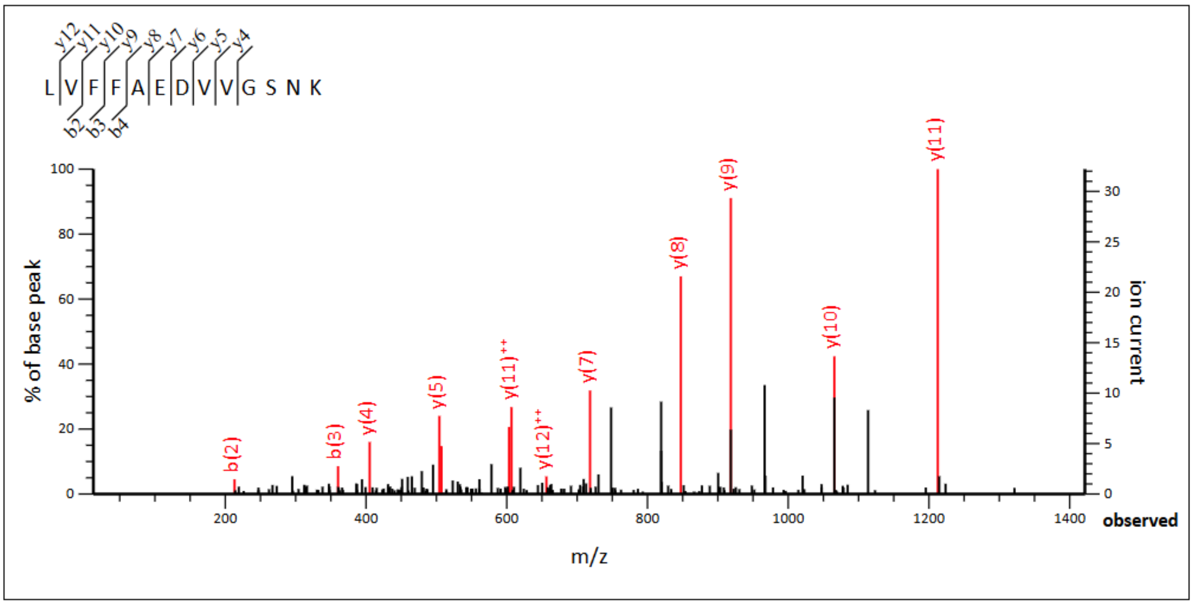


**Monoisotopic mass of neutral peptide Mr(calc):** 1423.7347
**Ions Score:** 54  **Expect:** 4.2e-06
**Matches :** 13/108 fragment ions using 25 most intense peaks

| # | b | $b^{++}$ | b* | $b^{*++}$ | $b^{0}$ | $b^{0++}$ | Seq. | y | $y^{++}$ | y* | $y^{*++}$ | $y^{0}$ | $y^{0++}$ | # |
|---|---|---|---|---|---|---|---|---|---|---|---|---|---|---|
| 1 | 114.0913 | 57.5493 | | | | | L | | | | | | | 13 |
| 2 | **213.1598** | 107.0835 | | | | | V | 1311.6579 | **656.3326** | 1294.6314 | 647.8193 | 1293.6474 | 647.3273 | 12 |
| 3 | **360.2282** | 180.6177 | | | | | F | ***1212.5895*** | **606.7984** | 1195.5630 | 598.2851 | 1194.5790 | 597.7931 | 11 |
| 4 | **507.2966** | 254.1519 | | | | | F | ***1065.5211*** | 533.2642 | 1048.4946 | 524.7509 | 1047.5106 | 524.2589 | 10 |
| 5 | 578.3337 | 289.6705 | | | | | A | ***918.4527*** | 459.7300 | 901.4262 | 451.2167 | 900.4421 | 450.7247 | 9 |
| 6 | 707.3763 | 354.1918 | | | 689.3657 | 345.1865 | E | ***847.4156*** | 424.2114 | 830.3890 | 415.6982 | 829.4050 | 415.2061 | 8 |
| 7 | 822.4032 | 411.7053 | | | 804.3927 | 402.7000 | D | ***718.3730*** | 359.6901 | 701.3464 | 351.1769 | 700.3624 | 350.6849 | 7 |
| 8 | 921.4716 | 461.2395 | | | 903.4611 | 452.2342 | V | ***603.3461*** | 302.1767 | 586.3195 | 293.6634 | 585.3355 | 293.1714 | 6 |
| 9 | 1020.5401 | 510.7737 | | | 1002.5295 | 501.7684 | V | ***504.2776*** | 252.6425 | 487.2511 | 244.1292 | 486.2671 | 243.6372 | 5 |
| 10 | 1077.5615 | 539.2844 | | | 1059.5510 | 530.2791 | G | ***405.2092*** | 203.1082 | 388.1827 | 194.5950 | 387.1987 | 194.1030 | 4 |
| 11 | 1164.5936 | 582.8004 | | | 1146.5830 | 573.7951 | S | 348.1878 | 174.5975 | 331.1612 | 166.0842 | 330.1772 | 165.5922 | 3 |
| 12 | 1278.6365 | 639.8219 | 1261.6099 | 631.3086 | 1260.6259 | 630.8166 | N | 261.1557 | 131.0815 | 244.1292 | 122.5682 | | | 2 |
| 13 | | | | | | | K | 147.1128 | 74.0600 | 130.0863 | 65.5468 | | | 1 |

## G25

MS/MS Fragmentation of peptide **LVFFAEDVGGSNK**

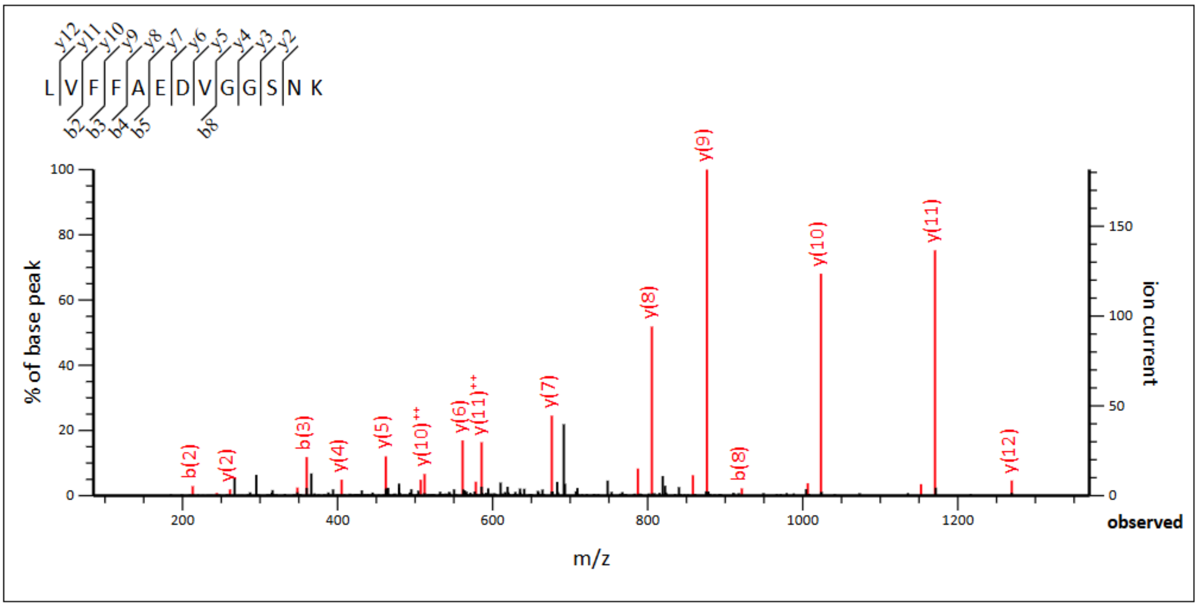


**Monoisotopic mass of neutral peptide Mr(calc):** 1381.6878
**Ions Score:** 80 **Expect:** 9e-09
**Matches :** 23/108 fragment ions using 43 most intense peaks

| # | b | $b^{++}$ | b* | $b^{*++}$ | $b^0$ | $b^{0++}$ | Seq. | y | $y^{++}$ | y* | $y^{*++}$ | $y^0$ | $y^{0++}$ | # |
|---|---|---|---|---|---|---|---|---|---|---|---|---|---|---|
| 1 | 114.0913 | 57.5493 | | | | | L | | | | | | | 13 |
| 2 | 213.1598 | 107.0835 | | | | | V | 1269.6110 | 635.3091 | 1252.5844 | 626.7959 | 1251.6004 | 626.3039 | 12 |
| 3 | 360.2282 | 180.6177 | | | | | F | 1170.5426 | 585.7749 | 1153.5160 | 577.2617 | 1152.5320 | 576.7696 | 11 |
| 4 | 507.2966 | 254.1519 | | | | | F | 1023.4742 | 512.2407 | 1006.4476 | 503.7274 | 1005.4636 | 503.2354 | 10 |
| 5 | 578.3337 | 289.6705 | | | | | A | 876.4058 | 438.7065 | 859.3792 | 430.1932 | 858.3952 | 429.7012 | 9 |
| 6 | 707.3763 | 354.1918 | | | 689.3657 | 345.1865 | E | 805.3686 | 403.1880 | 788.3421 | 394.6747 | 787.3581 | 394.1827 | 8 |
| 7 | 822.4032 | 411.7053 | | | 804.3927 | 402.7000 | D | 676.3260 | 338.6667 | 659.2995 | 330.1534 | 658.3155 | 329.6614 | 7 |
| 8 | 921.4716 | 461.2395 | | | 903.4611 | 452.2342 | V | 561.2991 | 281.1532 | 544.2726 | 272.6399 | 543.2885 | 272.1479 | 6 |
| 9 | 978.4931 | 489.7502 | | | 960.4825 | 480.7449 | G | 462.2307 | 231.6190 | 445.2041 | 223.1057 | 444.2201 | 222.6137 | 5 |
| 10 | 1035.5146 | 518.2609 | | | 1017.5040 | 509.2556 | G | 405.2092 | 203.1082 | 388.1827 | 194.5950 | 387.1987 | 194.1030 | 4 |
| 11 | 1122.5466 | 561.7769 | | | 1104.5360 | 552.7717 | S | 348.1878 | 174.5975 | 331.1612 | 166.0842 | 330.1772 | 165.5922 | 3 |
| 12 | 1236.5895 | 618.7984 | 1219.5630 | 610.2851 | 1218.5790 | 609.7931 | N | 261.1557 | 131.0815 | 244.1292 | 122.5682 | | | 2 |
| 13 | | | | | | | K | 147.1128 | 74.0600 | 130.0863 | 65.5468 | | | 1 |

## N27

MS/MS Fragmentation of peptide **LVFFAEDVGSNNK**

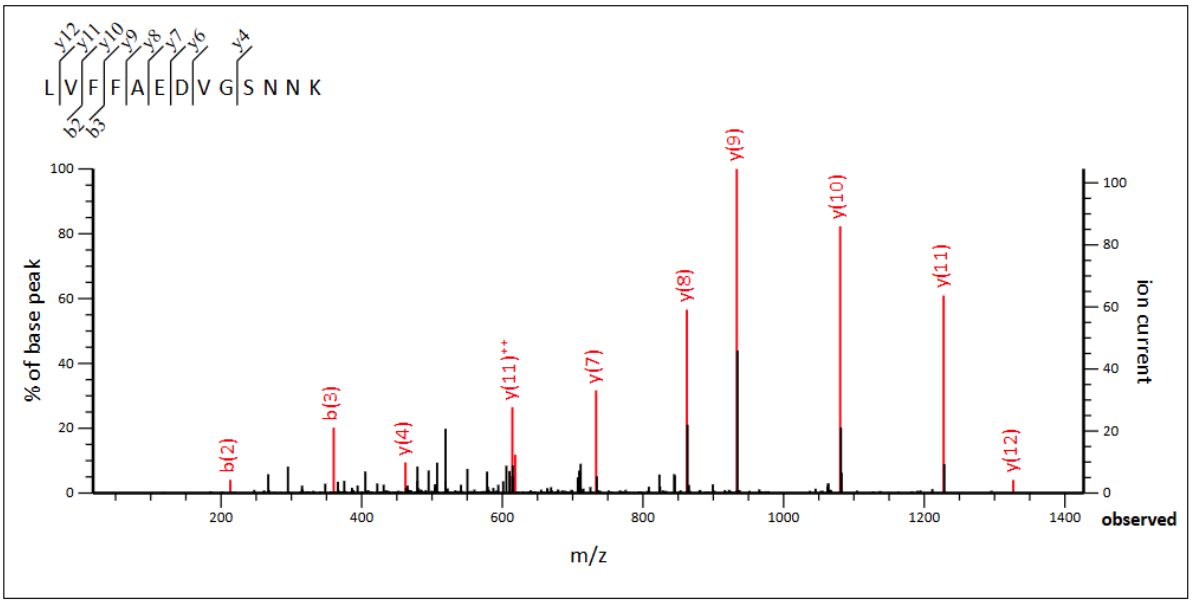


**Monoisotopic mass of neutral peptide Mr(calc):** 1438.7092
**Ions Score:** 76 **Expect:** 2.8e-08
**Matches :** 11/108 fragment ions using 13 most intense peaks

| # | b | $b^{++}$ | b* | $b^{*++}$ | $b^{0}$ | $b^{0++}$ | Seq. | y | $y^{++}$ | y* | $y^{*++}$ | $y^{0}$ | $y^{0++}$ | # |
|---|---|---|---|---|---|---|---|---|---|---|---|---|---|---|
| 1 | 114.0913 | 57.5493 | | | | | L | | | | | | | 13 |
| 2 | **213.1598** | 107.0835 | | | | | V | ***1326.6325*** | 663.8199 | 1309.6059 | 655.3066 | 1308.6219 | 654.8146 | 12 |
| 3 | **360.2282** | 180.6177 | | | | | F | ***1227.5640*** | **614.2857** | 1210.5375 | 605.7724 | 1209.5535 | 605.2804 | 11 |
| 4 | 507.2966 | 254.1519 | | | | | F | ***1080.4956*** | 540.7515 | 1063.4691 | 532.2382 | 1062.4851 | 531.7462 | 10 |
| 5 | 578.3337 | 289.6705 | | | | | A | ***933.4272*** | 467.2172 | 916.4007 | 458.7040 | 915.4166 | 458.2120 | 9 |
| 6 | 707.3763 | 354.1918 | | | 689.3657 | 345.1865 | E | ***862.3901*** | 431.6987 | 845.3636 | 423.1854 | 844.3795 | 422.6934 | 8 |
| 7 | 822.4032 | 411.7053 | | | 804.3927 | 402.7000 | D | ***733.3475*** | 367.1774 | 716.3210 | 358.6641 | 715.3369 | 358.1721 | 7 |
| 8 | 921.4716 | 461.2395 | | | 903.4611 | 452.2342 | V | ***618.3206*** | 309.6639 | 601.2940 | 301.1506 | 600.3100 | 300.6586 | 6 |
| 9 | 978.4931 | 489.7502 | | | 960.4825 | 480.7449 | G | 519.2521 | 260.1297 | 502.2256 | 251.6164 | 501.2416 | 251.1244 | 5 |
| 10 | 1065.5251 | 533.2662 | | | 1047.5146 | 524.2609 | S | ***462.2307*** | 231.6190 | 445.2041 | 223.1057 | 444.2201 | 222.6137 | 4 |
| 11 | 1179.5681 | 590.2877 | 1162.5415 | 581.7744 | 1161.5575 | 581.2824 | N | 375.1987 | 188.1030 | 358.1721 | 179.5897 | | | 3 |
| 12 | 1293.6110 | 647.3091 | 1276.5844 | 638.7959 | 1275.6004 | 638.3039 | N | 261.1557 | 131.0815 | 244.1292 | 122.5682 | | | 2 |
| 13 | | | | | | | K | 147.1128 | 74.0600 | 130.0863 | 65.5468 | | | 1 |

#### In rAβ42

### Y10

No identifications in rAβ42

### V12

No identifications in rAβ42

### L17

No identifications in rAβ42

### V18

No identifications in rAβ42

## F19/20

MS/MS Fragmentation of peptide **LVFFFAEDVGSNK**

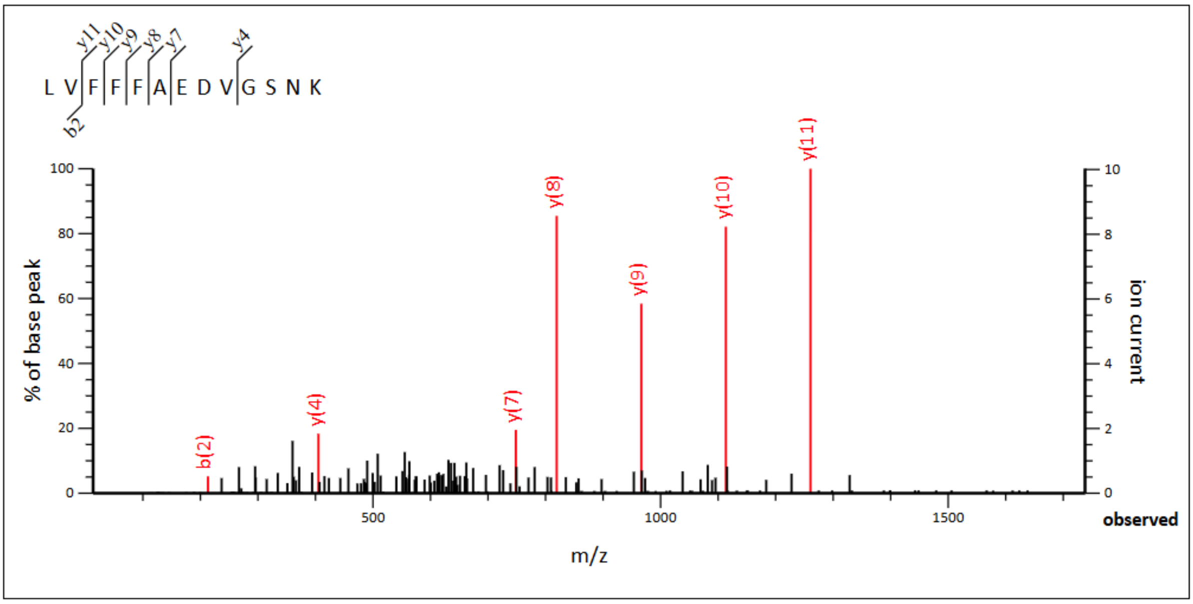


**Monoisotopic mass of neutral peptide Mr(calc):** 1471.7347
**Ions Score:** 35  **Expect:** 0.00035
**Matches :** 7/106 fragment ions using 14 most intense peaks

| # | b | $b^{++}$ | $b^{*}$ | $b^{*++}$ | $b^{0}$ | $b^{0++}$ | Seq. | y | $y^{++}$ | $y^{*}$ | $y^{*++}$ | $y^{0}$ | $y^{0++}$ | # |
|---|---|---|---|---|---|---|---|---|---|---|---|---|---|---|
| 1 | 114.0913 | 57.5493 | | | | | L | | | | | | | 13 |
| 2 | **213.1598** | 107.0835 | | | | | V | 1359.6579 | 680.3326 | 1342.6314 | 671.8193 | 1341.6474 | 671.3273 | 12 |
| 3 | 360.2282 | 180.6177 | | | | | F | ***1260.5895*** | 630.7984 | 1243.5630 | 622.2851 | 1242.5790 | 621.7931 | 11 |
| 4 | 507.2966 | 254.1519 | | | | | F | ***1113.5211*** | 557.2642 | 1096.4946 | 548.7509 | 1095.5106 | 548.2589 | 10 |
| 5 | 654.3650 | 327.6861 | | | | | F | ***966.4527*** | 483.7300 | 949.4262 | 475.2167 | 948.4421 | 474.7247 | 9 |
| 6 | 725.4021 | 363.2047 | | | | | A | ***819.3843*** | 410.1958 | 802.3577 | 401.6825 | 801.3737 | 401.1905 | 8 |
| 7 | 854.4447 | 427.7260 | | | 836.4341 | 418.7207 | E | ***748.3472*** | 374.6772 | 731.3206 | 366.1639 | 730.3366 | 365.6719 | 7 |
| 8 | 969.4716 | 485.2395 | | | 951.4611 | 476.2342 | D | 619.3046 | 310.1559 | 602.2780 | 301.6427 | 601.2940 | 301.1506 | 6 |
| 9 | 1068.5401 | 534.7737 | | | 1050.5295 | 525.7684 | V | 504.2776 | 252.6425 | 487.2511 | 244.1292 | 486.2671 | 243.6372 | 5 |
| 10 | 1125.5615 | 563.2844 | | | 1107.5510 | 554.2791 | G | ***405.2092*** | 203.1082 | 388.1827 | 194.5950 | 387.1987 | 194.1030 | 4 |
| 11 | 1212.5936 | 606.8004 | | | 1194.5830 | 597.7951 | S | 348.1878 | 174.5975 | 331.1612 | 166.0842 | 330.1772 | 165.5922 | 3 |
| 12 | 1326.6365 | 663.8219 | 1309.6099 | 655.3086 | 1308.6259 | 654.8166 | N | 261.1557 | 131.0815 | 244.1292 | 122.5682 | | | 2 |
| 13 | | | | | | | K | 147.1128 | 74.0600 | 130.0863 | 65.5468 | | | 1 |

## A21

MS/MS Fragmentation of peptide **LVFFAAEDVGSNK**

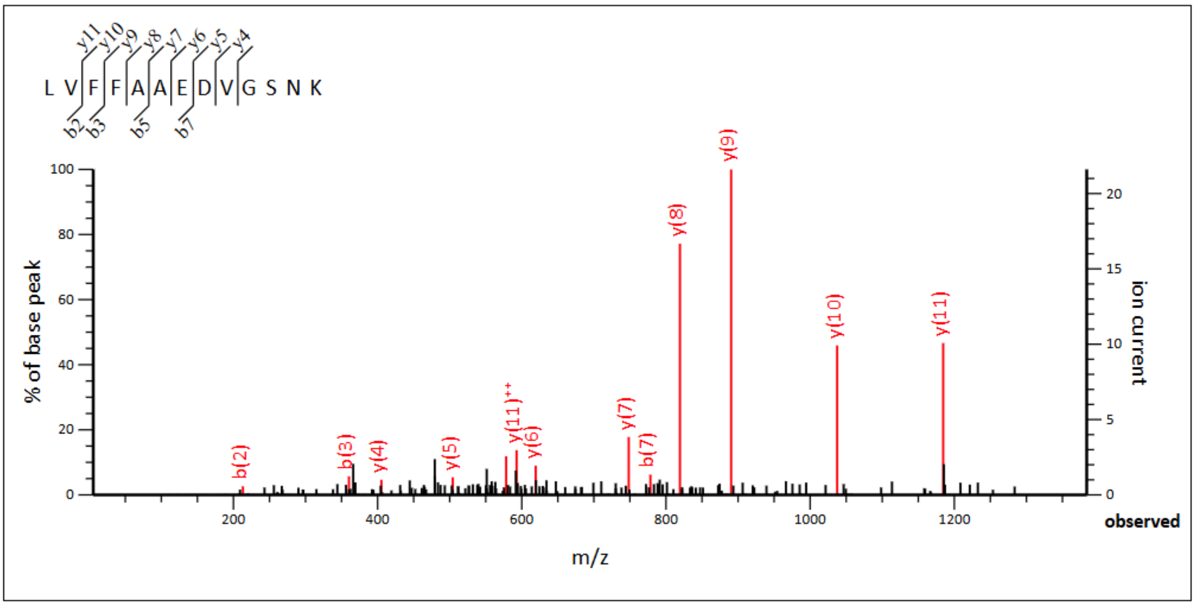


**Monoisotopic mass of neutral peptide Mr(calc):** 1395.7034
**Ions Score:** 38 **Expect:** 0.00016
**Matches :** 13/106 fragment ions using 36 most intense peaks

| # | b | b$^{++}$ | b* | b*$^{++}$ | b$^{0}$ | b$^{0++}$ | Seq. | y | y$^{++}$ | y* | y*$^{++}$ | y$^{0}$ | y$^{0++}$ | # |
|---|---|---|---|---|---|---|---|---|---|---|---|---|---|---|
| 1 | 114.0913 | 57.5493 | | | | | L | | | | | | | 13 |
| 2 | **213.1598** | 107.0835 | | | | | V | 1283.6266 | 642.3170 | 1266.6001 | 633.8037 | 1265.6161 | 633.3117 | 12 |
| 3 | **360.2282** | 180.6177 | | | | | F | ***1184.5582*** | **592.7828** | 1167.5317 | 584.2695 | 1166.5477 | 583.7775 | 11 |
| 4 | 507.2966 | 254.1519 | | | | | F | ***1037.4898*** | 519.2485 | 1020.4633 | 510.7353 | 1019.4793 | 510.2433 | 10 |
| 5 | **578.3337** | 289.6705 | | | | | A | ***890.4214*** | 445.7143 | 873.3949 | 437.2011 | 872.4108 | 436.7091 | 9 |
| 6 | 649.3708 | 325.1890 | | | | | A | ***819.3843*** | 410.1958 | 802.3577 | 401.6825 | 801.3737 | 401.1905 | 8 |
| 7 | **778.4134** | 389.7103 | | | 760.4028 | 380.7051 | E | ***748.3472*** | 374.6772 | 731.3206 | 366.1639 | 730.3366 | 365.6719 | 7 |
| 8 | 893.4403 | 447.2238 | | | 875.4298 | 438.2185 | D | ***619.3046*** | 310.1559 | 602.2780 | 301.6427 | 601.2940 | 301.1506 | 6 |
| 9 | 992.5088 | 496.7580 | | | 974.4982 | 487.7527 | V | ***504.2776*** | 252.6425 | 487.2511 | 244.1292 | 486.2671 | 243.6372 | 5 |
| 10 | 1049.5302 | 525.2688 | | | 1031.5197 | 516.2635 | G | ***405.2092*** | 203.1082 | 388.1827 | 194.5950 | 387.1987 | 194.1030 | 4 |
| 11 | 1136.5623 | 568.7848 | | | 1118.5517 | 559.7795 | S | 348.1878 | 174.5975 | 331.1612 | 166.0842 | 330.1772 | 165.5922 | 3 |
| 12 | 1250.6052 | 625.8062 | 1233.5786 | 617.2930 | 1232.5946 | 616.8009 | N | 261.1557 | 131.0815 | 244.1292 | 122.5682 | | | 2 |
| 13 | | | | | | | K | 147.1128 | 74.0600 | 130.0863 | 65.5468 | | | 1 |

# E22

MS/MS Fragmentation of peptide **LVFFAEEDVGSNK**

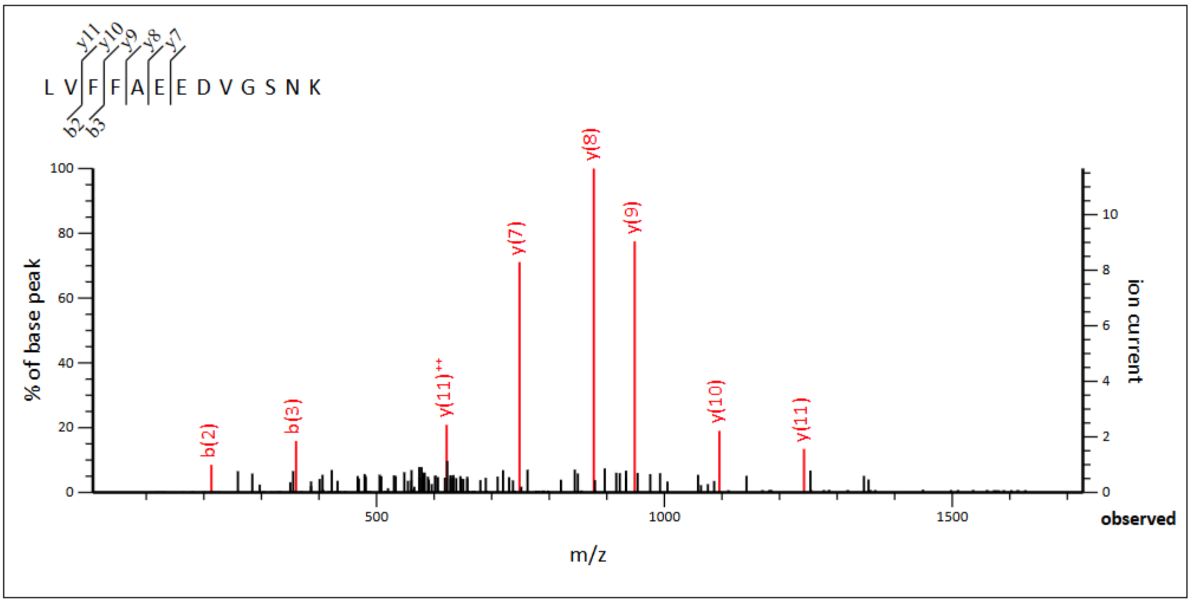


**Monoisotopic mass of neutral peptide Mr(calc):** 1453.7089
**Ions Score:** 28 **Expect:** 0.0014
**Matches :** 8/108 fragment ions using 14 most intense peaks

| # | b | b$^{++}$ | b* | b*$^{++}$ | b$^{0}$ | b$^{0++}$ | Seq. | y | y$^{++}$ | y* | y*$^{++}$ | y$^{0}$ | y$^{0++}$ | # |
|---|---|---|---|---|---|---|---|---|---|---|---|---|---|---|
| 1 | 114.0913 | 57.5493 | | | | | L | | | | | | | 13 |
| 2 | **213.1598** | 107.0835 | | | | | V | 1341.6321 | 671.3197 | 1324.6056 | 662.8064 | 1323.6216 | 662.3144 | 12 |
| 3 | **360.2282** | 180.6177 | | | | | F | ***1242.5637*** | **621.7855** | 1225.5372 | 613.2722 | 1224.5531 | 612.7802 | 11 |
| 4 | 507.2966 | 254.1519 | | | | | F | ***1095.4953*** | 548.2513 | 1078.4687 | 539.7380 | 1077.4847 | 539.2460 | 10 |
| 5 | 578.3337 | 289.6705 | | | | | A | ***948.4269*** | 474.7171 | 931.4003 | 466.2038 | 930.4163 | 465.7118 | 9 |
| 6 | 707.3763 | 354.1918 | | | 689.3657 | 345.1865 | E | ***877.3898*** | 439.1985 | 860.3632 | 430.6852 | 859.3792 | 430.1932 | 8 |
| 7 | 836.4189 | 418.7131 | | | 818.4083 | 409.7078 | E | ***748.3472*** | 374.6772 | 731.3206 | 366.1639 | 730.3366 | 365.6719 | 7 |
| 8 | 951.4458 | 476.2266 | | | 933.4353 | 467.2213 | D | 619.3046 | 310.1559 | 602.2780 | 301.6427 | 601.2940 | 301.1506 | 6 |
| 9 | 1050.5142 | 525.7608 | | | 1032.5037 | 516.7555 | V | 504.2776 | 252.6425 | 487.2511 | 244.1292 | 486.2671 | 243.6372 | 5 |
| 10 | 1107.5357 | 554.2715 | | | 1089.5251 | 545.2662 | G | 405.2092 | 203.1082 | 388.1827 | 194.5950 | 387.1987 | 194.1030 | 4 |
| 11 | 1194.5677 | 597.7875 | | | 1176.5572 | 588.7822 | S | 348.1878 | 174.5975 | 331.1612 | 166.0842 | 330.1772 | 165.5922 | 3 |
| 12 | 1308.6107 | 654.8090 | 1291.5841 | 646.2957 | 1290.6001 | 645.8037 | N | 261.1557 | 131.0815 | 244.1292 | 122.5682 | | | 2 |
| 13 | | | | | | | K | 147.1128 | 74.0600 | 130.0863 | 65.5468 | | | 1 |

## V24

MS/MS Fragmentation of peptide **LVFFAEDVVGSNK**

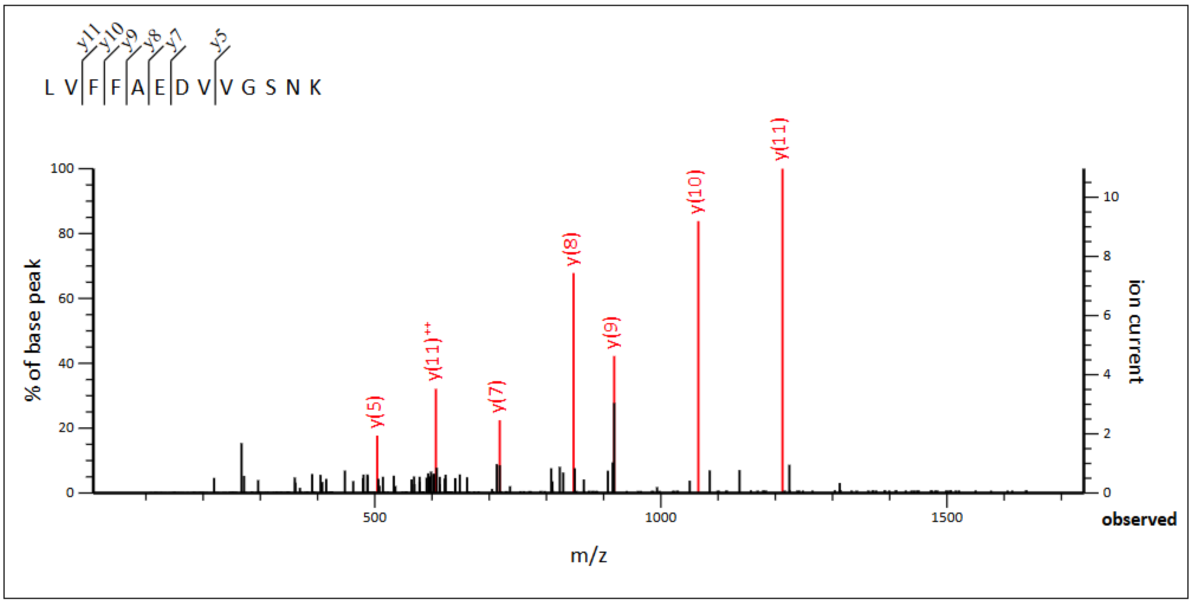


**Monoisotopic mass of neutral peptide Mr(calc):** 1423.7347
**Ions Score:** 36 **Expect:** 0.00025
**Matches :** 7/108 fragment ions using 14 most intense peaks

| # | b | $b^{++}$ | b* | $b^{*++}$ | $b^0$ | $b^{0++}$ | Seq. | y | $y^{++}$ | y* | $y^{*++}$ | $y^0$ | $y^{0++}$ | # |
|---|---|---|---|---|---|---|---|---|---|---|---|---|---|---|
| 1 | 114.0913 | 57.5493 | | | | | L | | | | | | | 13 |
| 2 | 213.1598 | 107.0835 | | | | | V | 1311.6579 | 656.3326 | 1294.6314 | 647.8193 | 1293.6474 | 647.3273 | 12 |
| 3 | 360.2282 | 180.6177 | | | | | F | ***1212.5895*** | **606.7984** | 1195.5630 | 598.2851 | 1194.5790 | 597.7931 | 11 |
| 4 | 507.2966 | 254.1519 | | | | | F | ***1065.5211*** | 533.2642 | 1048.4946 | 524.7509 | 1047.5106 | 524.2589 | 10 |
| 5 | 578.3337 | 289.6705 | | | | | A | ***918.4527*** | 459.7300 | 901.4262 | 451.2167 | 900.4421 | 450.7247 | 9 |
| 6 | 707.3763 | 354.1918 | | | 689.3657 | 345.1865 | E | ***847.4156*** | 424.2114 | 830.3890 | 415.6982 | 829.4050 | 415.2061 | 8 |
| 7 | 822.4032 | 411.7053 | | | 804.3927 | 402.7000 | D | ***718.3730*** | 359.6901 | 701.3464 | 351.1769 | 700.3624 | 350.6849 | 7 |
| 8 | 921.4716 | 461.2395 | | | 903.4611 | 452.2342 | V | 603.3461 | 302.1767 | 586.3195 | 293.6634 | 585.3355 | 293.1714 | 6 |
| 9 | 1020.5401 | 510.7737 | | | 1002.5295 | 501.7684 | V | ***504.2776*** | 252.6425 | 487.2511 | 244.1292 | 486.2671 | 243.6372 | 5 |
| 10 | 1077.5615 | 539.2844 | | | 1059.5510 | 530.2791 | G | 405.2092 | 203.1082 | 388.1827 | 194.5950 | 387.1987 | 194.1030 | 4 |
| 11 | 1164.5936 | 582.8004 | | | 1146.5830 | 573.7951 | S | 348.1878 | 174.5975 | 331.1612 | 166.0842 | 330.1772 | 165.5922 | 3 |
| 12 | 1278.6365 | 639.8219 | 1261.6099 | 631.3086 | 1260.6259 | 630.8166 | N | 261.1557 | 131.0815 | 244.1292 | 122.5682 | | | 2 |
| 13 | | | | | | | K | 147.1128 | 74.0600 | 130.0863 | 65.5468 | | | 1 |

## G25

MS/MS Fragmentation of peptide **LVFFAEDVGGSNK**

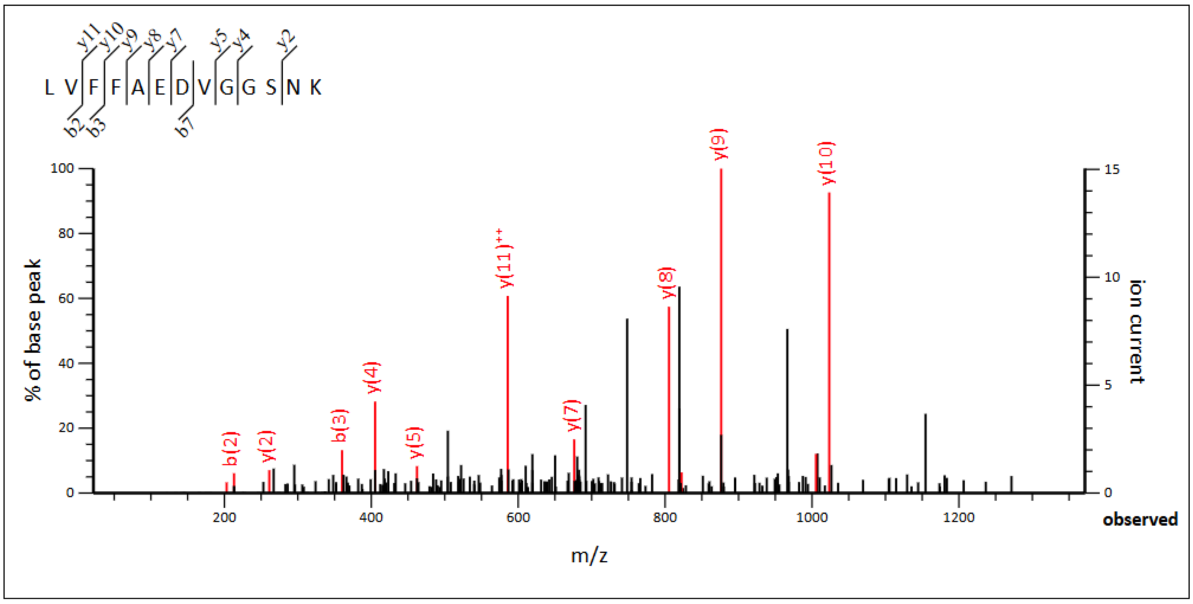


**Monoisotopic mass of neutral peptide Mr(calc):** 1381.6878
**Ions Score:** 32 **Expect:** 0.00063
**Matches :** 13/108 fragment ions using 35 most intense peaks

| # | b | b++ | b* | b*++ | b0 | b0++ | Seq. | y | y++ | y* | y*++ | y0 | y0++ | # |
|---|---|---|---|---|---|---|---|---|---|---|---|---|---|---|
| 1 | 114.0913 | 57.5493 | | | | | L | | | | | | | 13 |
| 2 | **213.1598** | 107.0835 | | | | | V | 1269.6110 | 635.3091 | 1252.5844 | 626.7959 | 1251.6004 | 626.3039 | 12 |
| 3 | **360.2282** | 180.6177 | | | | | F | 1170.5426 | **585.7749** | 1153.5160 | 577.2617 | 1152.5320 | 576.7696 | 11 |
| 4 | 507.2966 | 254.1519 | | | | | F | ***1023.4742*** | 512.2407 | 1006.4476 | 503.7274 | 1005.4636 | 503.2354 | 10 |
| 5 | 578.3337 | 289.6705 | | | | | A | ***876.4058*** | 438.7065 | 859.3792 | 430.1932 | 858.3952 | 429.7012 | 9 |
| 6 | 707.3763 | 354.1918 | | | 689.3657 | 345.1865 | E | ***805.3686*** | 403.1880 | 788.3421 | 394.6747 | 787.3581 | 394.1827 | 8 |
| 7 | **822.4032** | 411.7053 | | | 804.3927 | 402.7000 | D | ***676.3260*** | 338.6667 | 659.2995 | 330.1534 | 658.3155 | 329.6614 | 7 |
| 8 | 921.4716 | 461.2395 | | | 903.4611 | 452.2342 | V | 561.2991 | 281.1532 | 544.2726 | 272.6399 | 543.2885 | 272.1479 | 6 |
| 9 | 978.4931 | 489.7502 | | | 960.4825 | 480.7449 | G | ***462.2307*** | 231.6190 | 445.2041 | 223.1057 | 444.2201 | 222.6137 | 5 |
| 10 | 1035.5146 | 518.2609 | | | 1017.5040 | 509.2556 | G | ***405.2092*** | **203.1082** | 388.1827 | 194.5950 | 387.1987 | 194.1030 | 4 |
| 11 | 1122.5466 | 561.7769 | | | 1104.5360 | 552.7717 | S | 348.1878 | 174.5975 | 331.1612 | 166.0842 | 330.1772 | 165.5922 | 3 |
| 12 | 1236.5895 | 618.7984 | 1219.5630 | 610.2851 | 1218.5790 | 609.7931 | N | ***261.1557*** | 131.0815 | 244.1292 | 122.5682 | | | 2 |
| 13 | | | | | | | K | 147.1128 | 74.0600 | 130.0863 | 65.5468 | | | 1 |

### N27

No identifications in rAβ42

**Supplementary References**

1. Meisl, G. *et al.* Molecular mechanisms of protein aggregation from global fitting of kinetic models. *Nat. Protoc.* **11**, 252–272 (2016).

2. Willander, H. *et al.* BRICHOS domains efficiently delay fibrillation of amyloid β-peptide. *Journal of Biological Chemistry* **287**, 31608–31617 (2012).
